# Integrative HDX-MS enables quantification of the conformational landscape of the sugar transporter XylE

**DOI:** 10.1101/2022.07.11.499559

**Authors:** Ruyu Jia, Richard T. Bradshaw, Valeria Calvaresi, Argyris Politis

**Author notes:** Authors contributed equally (Ruyu Jia, Richard T. Bradshaw).

## Abstract

A yet unresolved challenge in structural biology is to quantify conformational states of proteins underpinning function. This challenge is particularly acute for membrane proteins owing to the difficulties in stabilising them for i*n vitro* studies. To address this challenge, we present here an integrative strategy that combines hydrogen-deuterium exchange mass spectrometry (HDX-MS) with ensemble modelling. We benchmark our strategy on wild type and mutant conformers of XylE, a prototypical member of the ubiquitous Major Facilitator Superfamily (MFS) of transporters. Next, we apply our strategy to quantify conformational ensembles of XylE embedded in different lipid environments and identify key lipid contacts that modulate protein conformations. Further application of our integrative strategy to substrate-bound and inhibitor-bound ensembles, allowed us to unravel protein-ligand interactions contributing to the alternating access mechanism of secondary transport in atomistic detail. Overall, our study highlights the potential of integrative HDX-MS modelling to capture, accurately quantify and subsequently visualise co-populated states of membrane proteins in association with mutations and diverse substrates and inhibitors.

**For Table of Content Only:** 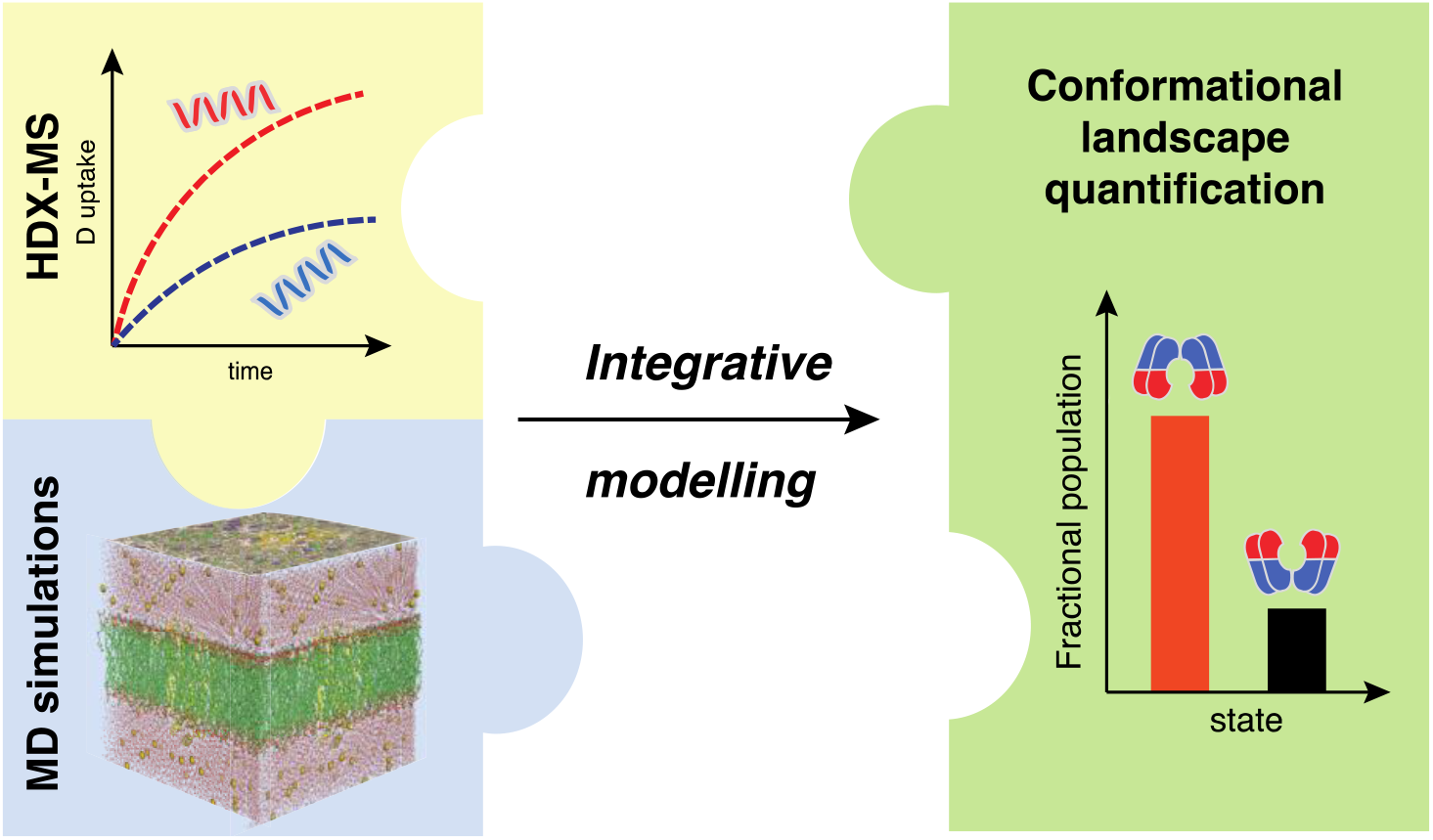

## Introduction

One of the challenges for structural biology for the upcoming decades is to evolve from assigning static structural snapshots, to characterising *dynamical ensembles*. Novel methodologies are required for the interrogation of membrane proteins, where existing tools, such as crystallography and cryogenic electron microscopy (cryo-EM), are often unsuccessful in probing the intricate interplay of protein and membrane dynamics underlying function. Furthermore, nuclear magnetic resonance (NMR) spectroscopy is limited to membrane-bound systems, where protein expression remains a challenging task. In contrast, HDX-MS is a well-established technique to probe the conformational dynamics of soluble proteins, which has recently emerged also for interrogating more complex protein systems^1^, such as membrane proteins^2, 3^. HDX-MS reports upon the deuterium exchange rates of backbone amide groups, averaged over oligopeptide protein segments. This technique offers advantages over traditional structural approaches, namely tolerance for complex, heterogeneous environments (e.g. lipids, detergents, native membranes), low sample requirements, and no need for bio-orthogonal labels. Importantly, HDX-MS data report on the equilibrium of protein conformational ensembles including all relevant populations. Therefore, HDX-MS has become a particularly powerful tool to study dynamical mechanisms inaccessible to other structural techniques, also in systems such as membrane proteins.

To infer structural information from HDX-MS data, the peptide-level exchange is often unsophisticatedly correlated to molecular simulations, to give qualitative insights into regions of relative flexibility. However, these simple insights neglect the full structural detail of the information that HDX-MS can provide, whereas the magnitude of dynamical changes that computational interpretations can study is not well understood. The simplicity of these analyses has in the past constrained HDX-MS to qualitative studies, complementary to more technically challenging higher-resolution methods. More advanced HDX-MS analyses instead promise to uncover information that existing crystallographic, or solution structural techniques are unable to provide - the full range of conformational states underpinning function in atomistic detail^4^.

A key issue in the interpretation of HDX-MS data, however, is the fact that the measured values of deuterium (D) uptake over time represents a conformationally-averaged exchange rate for the peptide of interest. Attempts to connect the observed HDX to structure (e.g. a single crystal structure, or single modelled conformational state) are therefore qualitative at best. They also risk being subjective, based on the availability of structural data or preconceived ideas of protein conformational states, rather than an objective selection of a model that best fits the HDX data. The structural context of HDX-MS data is, therefore, better represented as an ensemble of structures, such as those generated by the HDXer method^5^, or the Bayesian^6^ approach employed in HDX data intepretation^7^.

If HDX-MS experiments probe a highly diverse conformational ensemble, i.e. one interconverting between multiple conformational states, each with independent H-D exchange rates, then the ensemble-averaged HDX-MS signal may not be sufficient to unambiguously deconvolute the full extent of structural variation present. In studies of conformational mechanisms, this can be avoided by restricting proteins to particular conformational states as much as possible. In our own work, we have made use of single point mutations and ligand binding to probe single states of the XylE transport cycle independently of one another and describe the conformational process and key determinants of transport^2, 8^.

Here, we demonstrate an integrative approach to accurately describe conformational populations present in HDX-MS experiments. First, we benchmark our approach by calculating the populations present in the wild type (WT) apo XylE protein ensemble, and in an ensemble-driven towards an outward-facing (OF) conformational state by single point mutation (G58W). We find that the predicted conformational populations are relatively invariant to the predictive model used to connect protein structure to residue protection factors and exchange rate. Next, we conformationally describe subtle HDX differences observed between XylE ensembles in different lipid environments and identify specific protein-lipid contacts that discriminate between the two lipid conditions. Finally, we characterise the conformational effects of substrate binding to XylE and contrast the calculated substrate-bound ensemble with those of inhibitor-bound states. In doing so we provide an atomistic description of the mechanism of substrate and inhibitor recognition in XylE and demonstrate an important improvement in making useful structural interpretations with HDX-MS data.

## Results

### Benchmarking

We developed a three-step integrative approach that brings together experiments and computations. The steps involve, **a**) Gather and analyse HDX-MS data, **b**) Generate candidate model structures using MD simulations and subsequently extract simulated HDX-MS data, and **c**) Reweight the simulated data to fit a candidate model ensemble that conforms to the target data. Ultimately, the approach allows for quantifying the conformational populations of individual ensemble structures. In the first step, differential HDX-MS are carried out using previously described protocols^9^ (**Figure 1a**). This enables deuterium uptake measurements to be determined for identified peptides over various time intervals (ranging from seconds to hours). Differential HDX-MS experiments compare protein dynamics between two or more states, with improved precision compared to absolute HDX-MS, since experimental uncertainties cancel across the multiple protein measurements. Such datasets are ideal for qualitative analysis, but no computational approaches have been developed to predict or analyse ΔHDX-MS data directly. As a consequence, ΔHDX-MS can currently only describe qualitative structural trends between protein states, instead of precise conformational changes. Here, we normalised our ΔHDX-MS data to absolute HDX-MS data against a maximally deuterated (MaxD) sample, allowing for quantification of HDX-MS measurements. In the subsequent step, we deploy μs-long MD simulations to generate candidate structures of the protein under investigation (**Figure 1b**). We compute deuterium uptake associated with the generated model structures, using a common empirical model to estimate HDX protection factors (PF) from MD simulations. The model itself (**Equation 1**), estimates structural protection as an ensemble average function of the hydrogen bonds, *N*_*H*_, and heavy-atom interatomic contacts, *N*_*C*_, involving each amide.

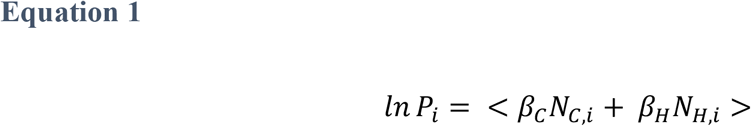

**Figure 1.**
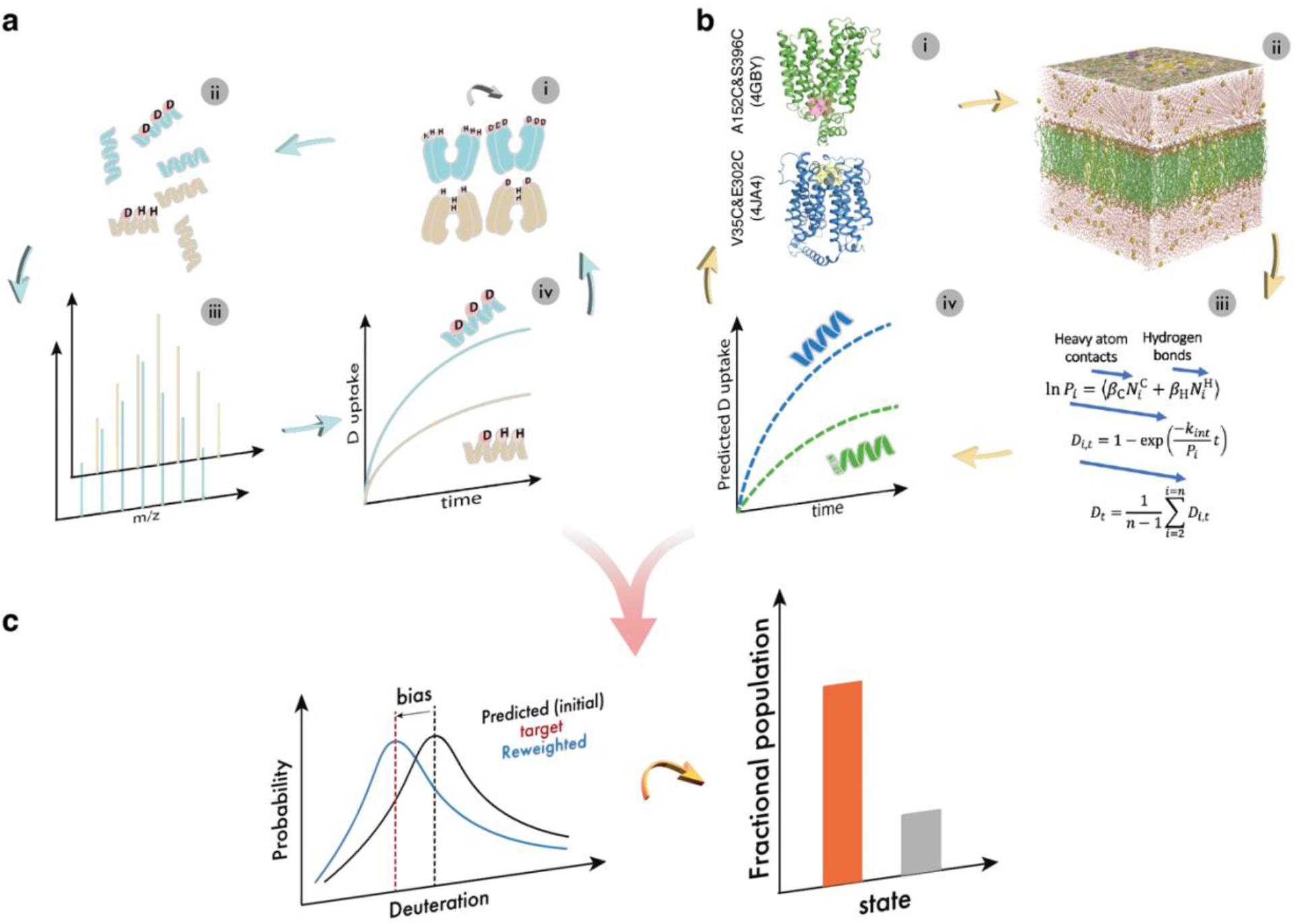
Integrative modelling workflow with HDX-MS data and molecular simulations. **(a,i)** The H-D exchange of protein backbone amides occurs spontaneously in deuterated solution. **(a,ii)** The exchange can be quenched at various time points and measured at peptide level resolution when coupled to enzymatic digestion. **(a,iii&iv)** Deuterium uptake over time is determined by measuring the peptide mass by LC-MS. **(b,i)** Generate initial protein structures. **(b,ii)** Ensemble sampling of protein structures via MD simulations. **(b,iii&iv)** Compute deuterium uptake based on an empirical model. **(c)** Fit predicted deuteration to target data to find an atomistic structural ensemble that best fits target HDX-MS data. Then quantify fractional population in the final reweighting ensemble structure.

In the final step, with the experimental and simulated HDX data in hand, we fit simulated deuteration to target experiments. We use our predictive model to adjust (‘reweight’) the relative populations of models to fit individual absolute HDX-MS data (**Figure 1c**). Overall, this step allows us to find an atomistic structural ensemble that best fits target HDX-MS data and to quantify the fractional population of individual conformational states in the final reweighted ensemble.

Initially, we benchmarked our strategy using existing data generated in our group^2^. In previous studies, we have shown that the equilibrium conformational populations of states in the transport cycle of XylE may be affected by mutations, proton- or ligand-binding, and detergent or lipid environment^28^. WT apo XylE has been crystallised only in an inward-open state, while observation of an outward-open state required a double mutation (G58W/L315W) of residues lining the extracellular vestibule, sterically hindering the transition to an inward-facing (IF) state. To compare experimental and predicted HDX-MS data, back-exchange corrected HDX-MS data is required by HDXer. After reanalysing the original differential HDX-MS data obtained by Martens *et al* in order to ensure consistent peptides could be studied across all subsequent states, we performed a maximally deuterated (MaxD) control to obtain the absolute HDX-MS data of WT and the single mutant G58W XylE. In WT XylE, peptides at the intracellular face of the protein exhibit significant deuteration while peptides at the extracellular face are comparatively highly protected. In G58W XylE the pattern is reversed and peptides at the extracellular face exhibit high deuteration, commensurate with a flexible, solvent-accessible conformational state (**Figure S1**). Differential HDX-MS analysis highlighted 35 peptides from previous results with significant differences in uptake between WT and G58W. Significant differences were entirely localised to the solvent-facing surfaces of the protein and were qualitatively consistent with those originally analysed^2^, suggesting that the G58W protein shifted the relative conformational populations of XylE towards more OF states (**Figure 2a**). Next, to quantify the magnitude of the shift in conformational populations represented by the previous experimental WT and G58W data^2^, we performed ensemble reweighting. A mixed candidate ensemble, initially of 50% OF and 50% IF structures, was fitted using HDXer to target each of the absolute HDX-MS datasets separately. Targeting the WT HDX-MS data resulted in a final reweighted ensemble of 4.4% OF, and 95.6% IF structures. In contrast, targeting the G58W HDX-MS data resulted in a final ensemble of 80.6% OF, and 19.4% IF structures (**Figure 2a**).

**Figure 2.**
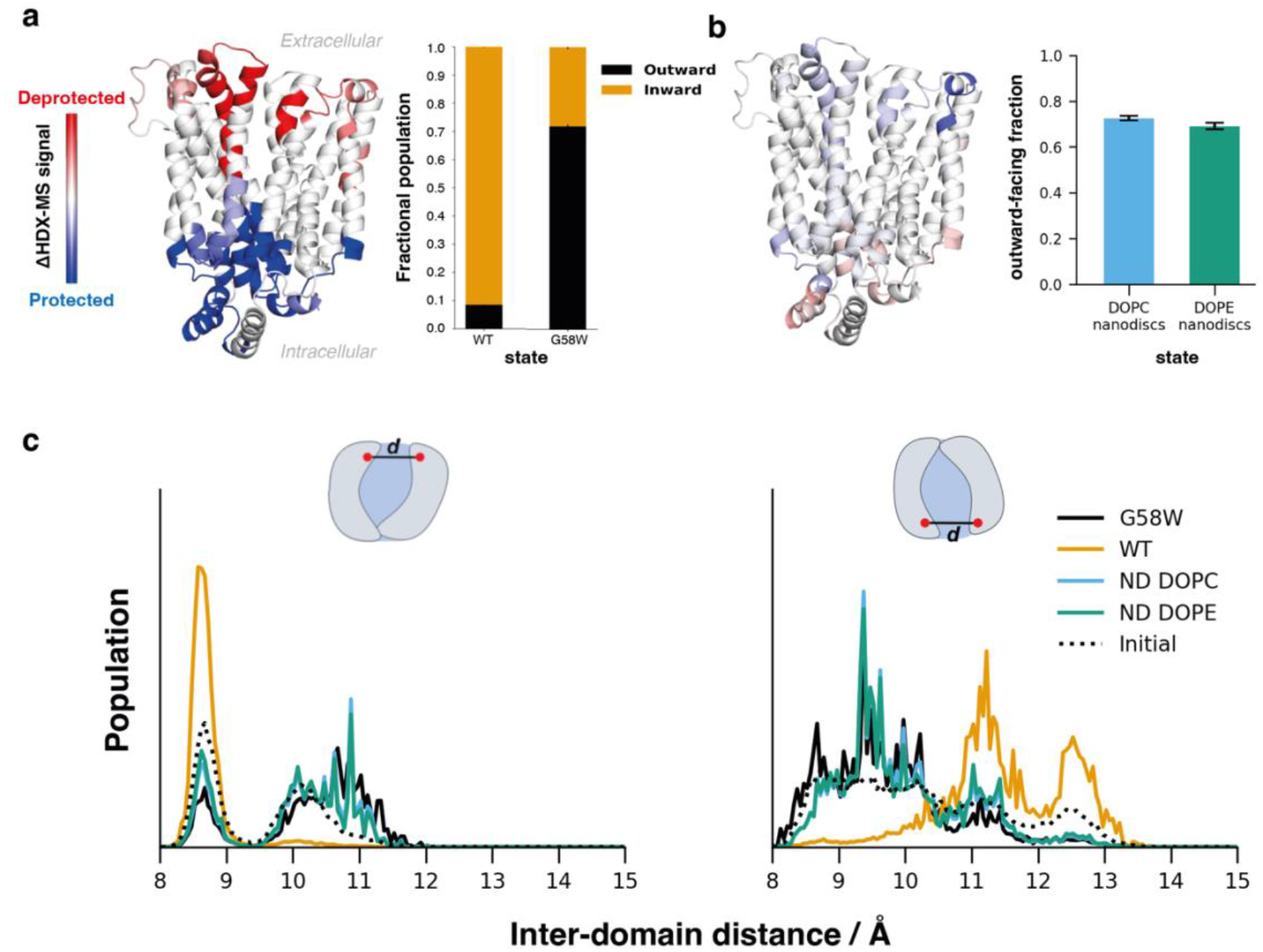
Benchmarking on XylE WT, G58W mutant and lipid nanodisc. **(a)**. Differential HDX-MS and reweighting results for XylE WT and G58W solution ensembles highlight significant differences in deuterium uptake between WT & G58W XylE, localised to the extracellular and intracellular protein faces. The results indicate that the G58W mutation shifts the XylE conformational ensemble to a more OF population than WT. Computational reweighting of a mixed OF/IF ensemble to fit the experimental WT and G58W. HDX-MS data results in a clear separation of the structures present in each experimental dataset. WT XylE is mostly IF (orange), and G58W is mostly OF (black)**. (b)** Differential HDX-MS and reweighting results for XylE WT in DOPC-based and DOPE-based nanodisc. Structural overlay of DOPC – DOPE ΔHDX-MS. After reweighting, both nanodisc environments consist of a mixed OF and IF ensemble, with populations intermediate between those of WT and G58W XylE in detergent micelles. DOPE-based nanodisc shift towards a slightly more IF ensemble than DOPC-based**. (c)** Effect of reweighting on the XylE inter-domain distance distributions on the extracellular and intracellular face of the protein. Larger distances correspond to a more ‘open’ structure.

The bias applied to (i.e. the final relative weight of) each structure in the reweighted ensemble is inextricably linked to the model used to predict residue protection factors from the structure. The Best-Vendruscolo model is parameterised to estimate the conformational free energy change of ‘opening’ (ΔG_op_) from simulations that predominantly sample the protein exchange- non-competent or ‘closed’ (C) state. Our simulations are approximately 10^3^-fold longer than the simulations used to parameterise the original Best-Vendruscolo model, and this additional sampling of structural fluctuations may reduce the accuracy of the parameterised model. We therefore investigated the sensitivity of the reweighted conformational populations to changes in the scaling parameters (*β*_*C*_ and *β*_*H*_) of the Best-Vendruscolo predictive model. Scaling factors were re-optimised for µs-scale dynamics, resulting in *β*_*C*_ = 0.29 (original *β*_*C*_ = 0.35), *β*_*H*_ = 3.9 (original *β*_*H*_ = 2.0), and reweighting was again applied to quantify the conformational populations present in the WT and G58W experimental data. The WT ensemble was calculated with final weights of 8.5% OF, and 91.5% IF, while the G58W ensemble was calculated to be 71.9% OF, and 28.1% IF (**Figure S2**).

Although the absolute magnitudes of the final reweighted populations do change between the two models, the overall structural interpretation that the G58W mutation shifts the conformational equilibrium from almost completely IF to almost completely OF is unchanged. The structural interpretations that reweighting can provide for XylE are therefore relatively robust to small uncertainties in the predictive model. Large conformational effects, e.g. associated with point mutations, are therefore clearly amenable to reweighting interpretations, and consistently determined. With that in mind, we pose the question: are smaller conformational/dynamical differences also interpretable?

Differential HDX signals between DOPC-based and DOPE-based nanodisc (ND) are largely non-significant (**Figure 2b**) when compared at the individual peptide level but may cumulatively, and following HDXer reweighting, the HDX measurements reflect a small but significant difference in global OF/IF populations between the PC- and PE-based lipidic environments.

The conformational shifts associated with each experimental dataset may be explored in greater detail by examining the inter-domain distance distributions for each reweighted ensemble. Here, a distinct difference between the WT and G58W or ND distributions is observed, particularly in the distances observed at the intracellular face of the protein (**Figure 2c**). After reweighting to the WT data, a substantial population is observed at an intracellular inter-domain distance of ∼13 Å, corresponding to a ‘fully-open’ conformation, while in reweighting to G58W or ND data, only a ‘partially-open’ conformation (inter-domain distance ∼ 11 Å) is observed.

To visualise the structural differences between partially and fully open conformations in more detail, we extracted representative structures from the intracellular distance distributions after reweighting to either the WT or G58W (**Figure S3 & S4**) HDX data. The larger inter-domain distance observed in the fully open conformation appeared to arise from the motion of the intracellular helical bundle (residues 220-270), which lies away from the intracellular vestibule, packed against the C-terminal intracellular helix. In contrast, in the partially open conformation, the intracellular bundle is swung back towards the vestibule opening, although the intracellular pathway to the binding site remains accessible. Although the fully open conformation is more consistent with an ‘inward-open’ state, we note that the intracellular helices are often only partly resolved in available crystal structures, thus supporting the idea of a high degree of flexibility in this region.

After reweighting to the G58W HDX-MS data, only small populations of IF structures remained. However, the OF conformations of XylE (corresponding to intracellular inter-domain distances of 8.0 – 10.6 Å) also showed substantial flexibility in the intracellular helical bundle. Although these four intracellular helices form a cap to the intracellular vestibule in the outward-open crystal structure, the flexibility of these regions observed in MD simulations appears to be consistent with our HDX-MS data. The final ensembles after reweighting to DOPC or DOPE-based nanodisc, however, were very similar in terms of OF/IF populations (**Figure 2b**) and inter-domain distances (**Figure 2c**). The number of intracellular lipid contacts to E153, D337, and E397, which had previously been identified as key residues controlling the inward-outward conformational preference^2^, also showed no appreciable difference after reweighting to either DOPC or DOPE nanodisc HDX-MS (**Figure S5**).

Overall, our analysis implies that the effects of lipid type upon XylE conformational preference are very subtle, although consistent with previous hypotheses. The similarity between the final ensemble populations highlights a limit of the structural fidelity provided by the ensemble reweighting approach – in particular, our target dataset for reweighting does not include peptides covering the E153, D337, or E397 residues, any (or all) of which may be crucial to characterise the subtle effects of lipid composition upon structure.

### HDX-MS reveals distinct protein dynamics upon substrate/ inhibitor binding

Having established the feasibility of our strategy in benchmarking XylE WT and G58W mutant using previously published differential HDX-MS data obtained^2^, we turned our attention to understanding distinct dynamics of XylE transporter upon substrate and inhibitor binding. We chose to use the substrate xylose and endogenous inhibitor, glucose, as well as exogenous ligands phloretin^10^ and phloridzin^11^, which are known inhibitors of glucose transmembrane transport^12^. A previous study has established XylE as a surrogate for the mechanistic understanding of GLUT1 via a conserved mechanism of ligand binding^13^.

To dissect the structural changes made by substrate and inhibitor binding before transport, we performed a new set of differential HDX-MS experiments, together with MaxD experiments, comparing XylE WT apo and in ligand-bound (xylose-, glucose-, phloretin- and phloridzin-bound) states (**Figure 3a**). Prior to experiments, the protein and ligands were incubated to achieve a binding occupancy of approximately 90% according to the binding affinity of each ligand (**Table S1**). We obtained 82% sequence coverage, allowing us to interrogate the dynamics of XylE in its ligand-bound and ligand-free states along most of the protein sequence (**Figure S6 & S7 & Table S2**).

**Figure 3.**
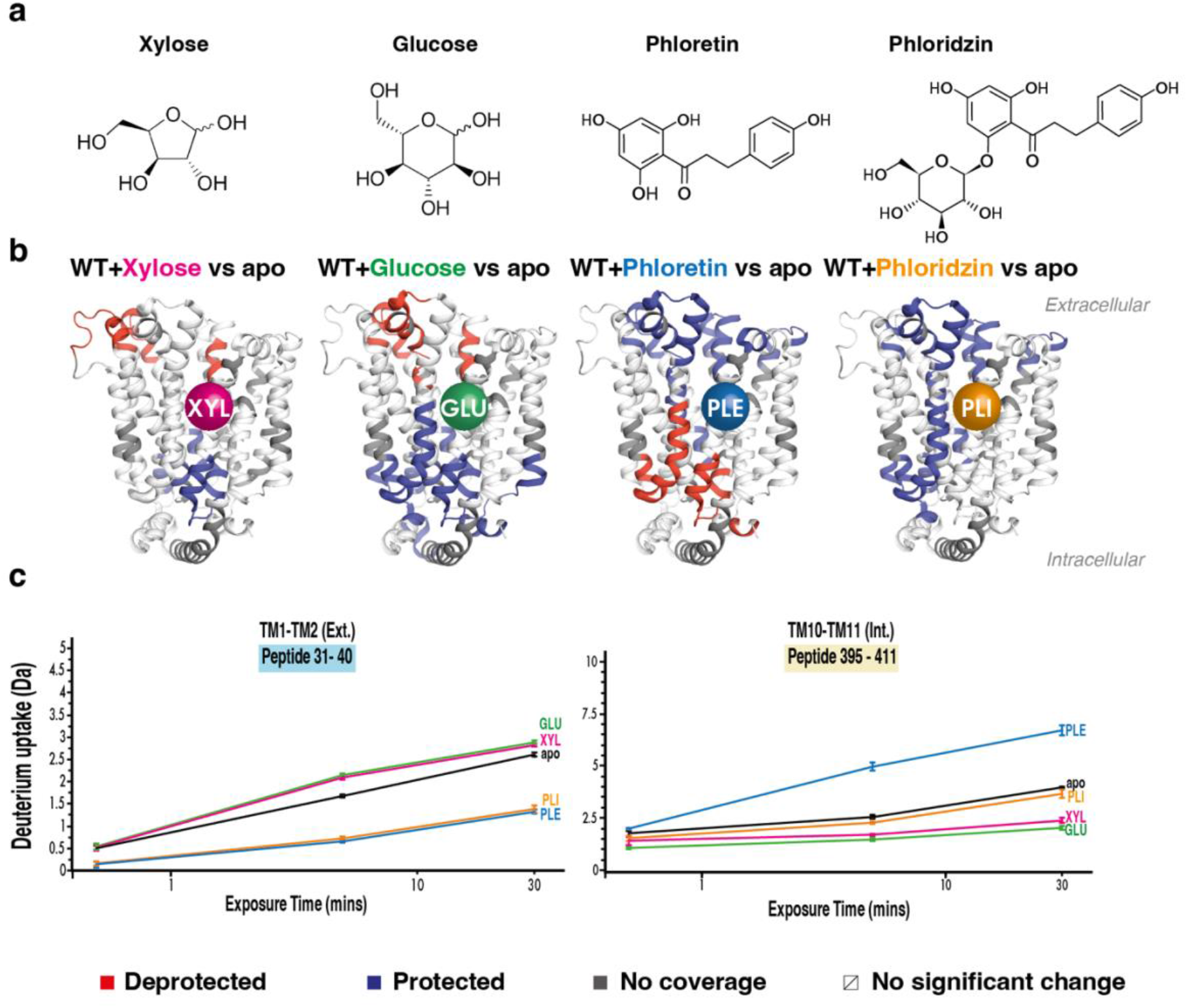
Differential HDX-MS experiments comparing XylE-ligand bound states to the apo state. **(a)** Chemical structures of ligands: Xylose, Glucose, Phloretin and Phloridzin. **(b)** Differential HDX-MS uptake pattern between WT and XylE-ligand bound structures. **(c)** Representative peptide deuterium uptake plots between WT and XylE-ligand bound structures (peptide 31- 40 on the extracellular side and 395- 411 on the intracellular side). Figures are plotted onto a 3D protein structure (PDB:4GBY). Blue and red regions indicate negative (protected) or positive (deprotected) deuterium uptake differences between states, respectively. SDs for each time point are plotted as error bars (n=3).

We began by carrying out differential HDX-MS experiments with the substrate xylose and inhibitor glucose. To obtain comparable conditions with other ligands, which required to be solubilised in DMSO, for each experiment we equilibrated the protein and ligand together in 10% DMSO solution prior to labelling. To ensure the presence of DMSO did not adversely affect protein dynamics, we compared these experiments to our previously published data^8^, and we observed analogous conformational fingerprints to the previous results (**Figure S8**). Indeed, in these new experiments, the presence of xylose and glucose leads to an increase in deuterium uptake on the extracellular side (e.g. peptide 31-40) coupled with a decrease on the intracellular side (e.g. peptide 396-411) (**Figure 3b, c**). Such deuterium uptake difference is a typical ΔHDX pattern of a transition towards an OF conformation.

Next, we explored the conformational landscape of XylE WT upon binding to phloretin and phloridzin. Interestingly, we observed a substantially different ΔHDX pattern in the presence of these GLUT inhibitors. Unlike xylose and glucose, phloretin-bound HDX fingerprint shows a decrease of deuterium uptake on the extracellular side (e.g. peptide 31-40) with an increase on the intracellular side (e.g. peptide 396-411), a ΔHDX pattern typical for the transition of transporter towards an IF conformation (**Figure 3b, c**). Intriguingly, compared to unbound XylE, the presence of phloridzin causes an overall decrease in deuterium uptake on both extracellular (e.g. peptide 31-40) and intracellular (e.g. peptide 396-411) sides (**Figure 3b, c**). For the phloretin-bound structure, the data suggest that phloretin binding drives the protein towards a more IF ensemble than the apo state. This likely precludes other ligands to bind to the extracellular side of the protein and eventually being transported. In the presence of phloridzin, a decrease in deuterium uptake from both extracellular and intracellular sides suggests an overall protection of the whole protein, consistent with an occluded-like state. It is interesting to speculate that such differences may be indicative of different inhibitory pathways, likely reflecting differences in the effectiveness in inhibiting the glucose transport, as previously suggested^14, 15^, although detailed biophysical and structural studies of these tool compounds are few in number. Our HDX-MS data suggests a “combinatorial” effect for the sugar (glucose) and aglucone (phloretin) moiety upon inhibitor phloridzin binding to the protein. To explore this further and to gain structural insight into the conformational landscape of the inhibitor-bound states, we proceeded to computational analyses using our integrative HDX-MS approach presented above.

### MD simulations to predict protein-inhibitor binding modes

Integrative ensemble reweighting to each protein-ligand HDX-MS dataset first required a comprehensive candidate ensemble incorporating both OF and IF ligand-bound structures for each ligand. However, crystal structures are only available for OF xylose-bound (4GBY), OF glucose-bound (4GBZ) and IF apo (4JA4) structures. Therefore, to generate ensembles for the remaining protein-ligand states, we first generated conformationally locked protein structures - OF and IF mutants of XylE restricted by cysteine cross-linking mutations (OF: A152C/S396C and IF: V35C/E302C) based on available crystal structures (4GBY and 4JA4), and performed MD simulations to sample the conformationally-locked apo-state pocket. Subsequently, we extracted representative pocket (and receptor) structures from each apo simulation and identified docked poses for each ligand using rigid receptor docking in Autodock Vina^16^. Protein flexibility was instead incorporated by docking to multiple representative structures from apo-state MD simulations, rather than a single crystal structure. We then subjected the docked poses to Principal Component Analysis (PCA) and clustering in the reduced dimensions to identify highly populated binding modes suitable to initiate MD simulations. Finally, a two-step MD simulation (100 ns and 1 µs) allowed us to validate the stability of the selected bound structures (**Figure 4**).

**Figure 4.**
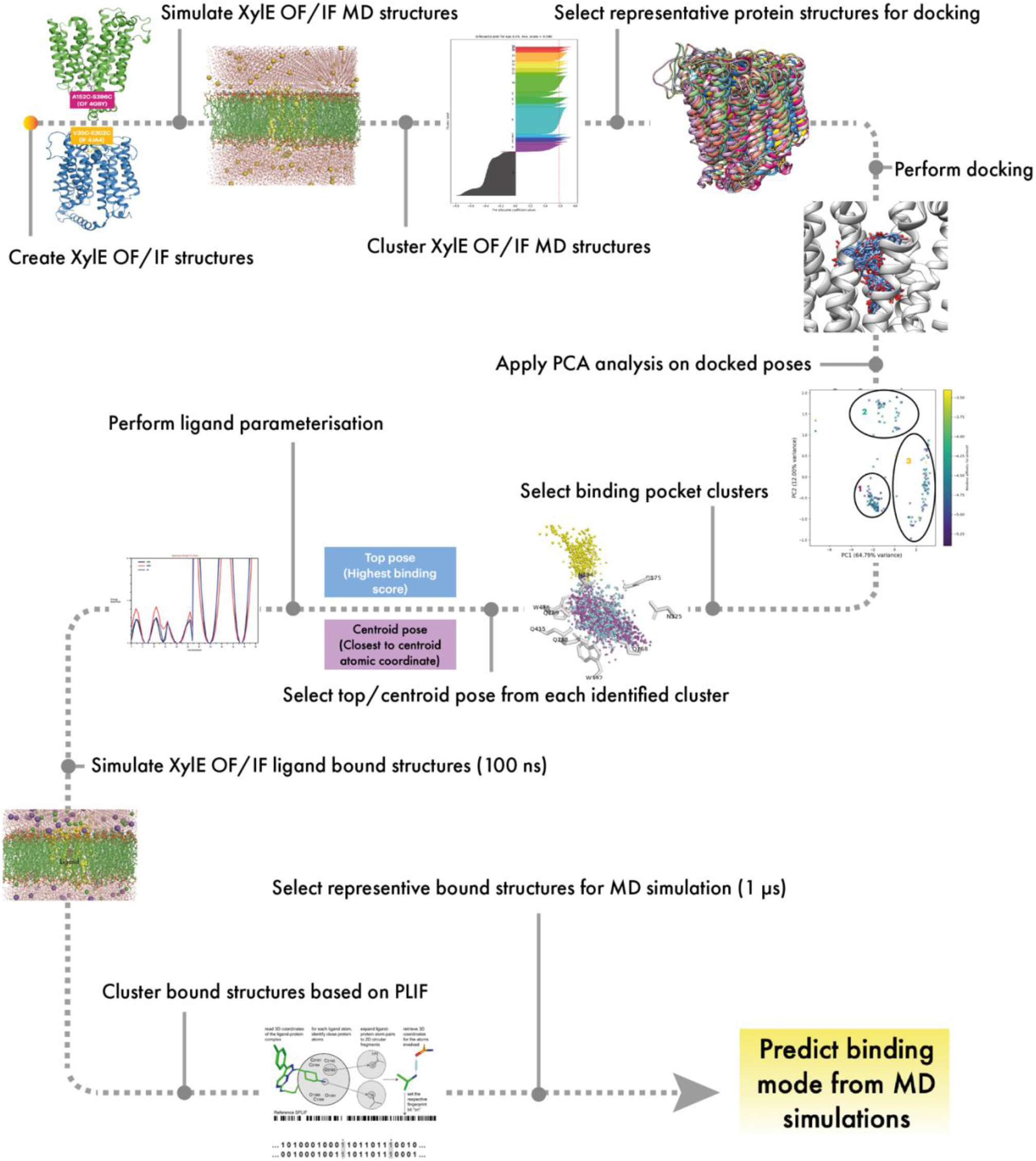
MD workflow of generating XylE phloretin- and phloridzin-bound structures. The workflow comprises generating apo receptor structures; Rigid docking of ligand to representative receptor structures; Dimensionality reduction and structural clustering of docked poses; Simulation of representative/highest scoring docked poses; Analysis of pose dynamic and selection of suitable binding pose by stability, repeated sampling and biological relevance criteria.

Analyses of the lengthier MD simulations suggest that protein and ligands remain stably bound in OF and IF structures during the vast majority of 1 µs simulations. (**Figure 5a, b & Figure S9**). A closer inspection of the data however reveals interesting differences between phloretin and phloridzin. The sugar moiety in phloridzin appears to have a prominent function in binding to XylE. Crystallographic studies have previously shown three glutamines (Gln168, Gln175 and Gln415) in the ligand-binding site are critical in D-glucose recognition by XylE. In particular, Gln168 forms three hydrogen bonds with D-glucose, while it only has one single hydrogen bond with D-xylose. Additionally, Gln175 is hydrogen-bonded with the 6-hydroxyl group of D-glucose but not involved in D-xylose binding^17^. Therefore, we performed hydrogen bond analysis for glucose-, phloretin-, and phloridzin-bound structures, where the number of hydrogen bonds was calculated over the 1 µs simulation time. Hydrogen bond interactions between ligand and residue Gln168, Gln175, and Gln415 were analysed separately (**Figure 5c & Figure S10**). Our MD simulations show that all three residues form hydrogen bond interactions with phloridzin in a similar pattern to D-glucose-bound structures in both OF and IF conformations. In contrast, phloretin-bound structures show substantially fewer H-bonds than D-glucose- and phloridzin-bound structures under all analysed conditions. The results suggest that the glucose-moiety in phloridzin plays a pivotal role in binding with XylE. Overall, our computational pose prediction workflow generated stably bound poses consistent with available experimental interaction information and provided new structural insights into the inhibitory function of phloretin and phloridzin upon binding to XylE.

**Figure 5.**
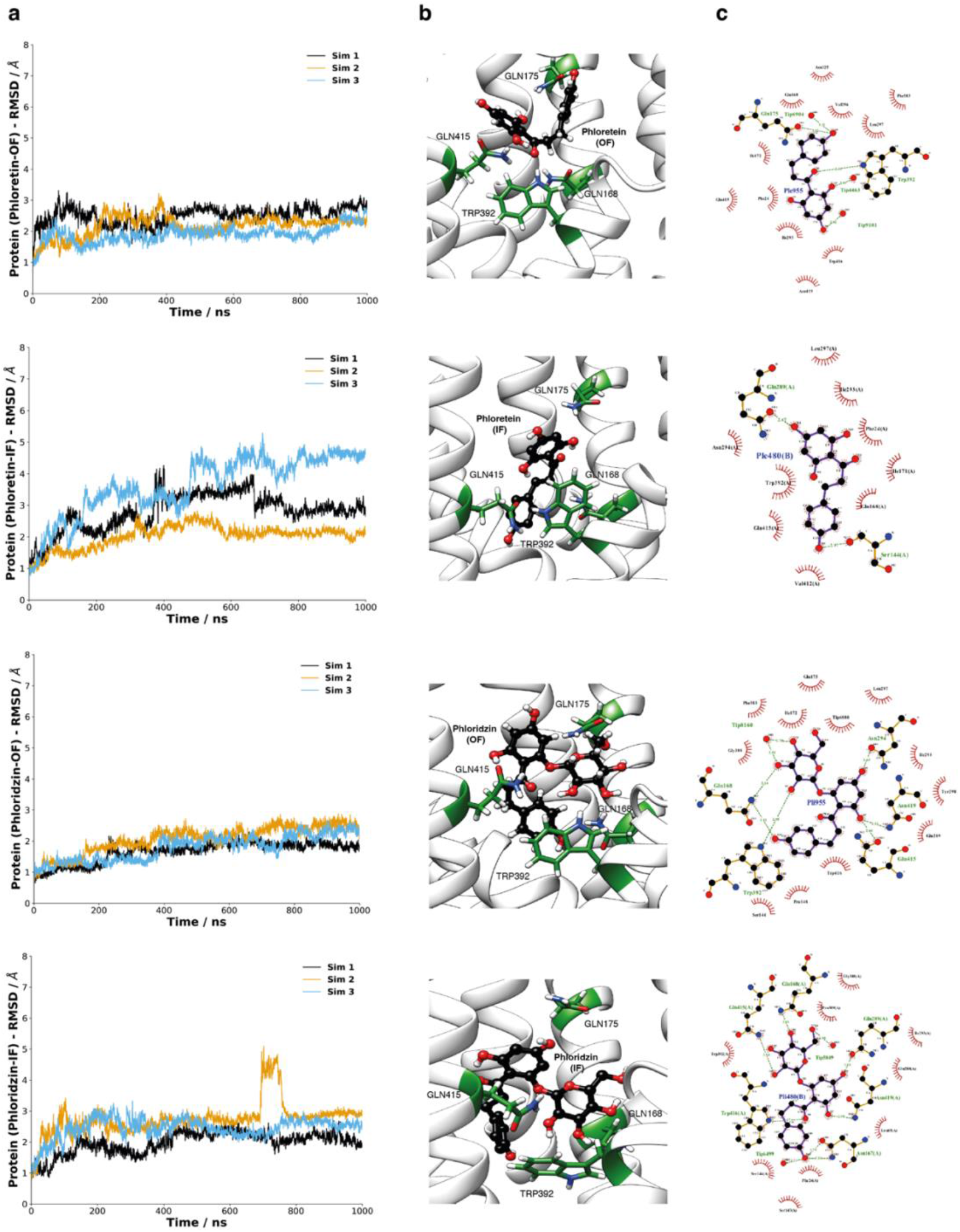
MD simulations of XylE phloretin- and phloridzin-bound structures in the OF and IF conformations. **(a)** RMSD plots of XylE backbone in bound phloretin and phloridzin (OF and IF) conformations through three independent 1µs long simulations. **(b)** Coordination of phloretin and phloridzin by XylE in OF and IF structures. Phloretin and phloridzin are shown in black balls and sticks. The binding site residues in XylE are coloured green. **(c)** 2D protein-ligand interaction diagram generated by LigPlot+^18^. Phloretin and phloridzin are shown in black balls and sticks, hydrogen bonds are shown as green dotted lines, while the red eyelash diagram represents hydrophobic interactions.

### HDX reweighting approaches quantify conformational population

Having gained insights into XylE-inhibitor binding modes by MD simulations, we sought to probe the conformational landscape of XylE in apo and ligand-bound states. Ensemble structures generated by MD simulations were used to predict HDX-MS deuterated fractions for peptide segments corresponding to our experimental HDX-MS data. The calculated deuteration fraction of each time point was compared with the corresponding experimental data. Peptides spanning residue 1-5 or 479 onwards (missing in the crystal structure) or with negative deuteration from experimental data (attributed to experimental noise) were excluded from reweighting analyses.

We performed HDXer analyses of each experimental HDX-MS dataset separately. Initially, for each HDXer analysis, we used a candidate ensemble comprising only the “state-specific” XylE states (i.e. apo simulations fit to apo HDX-MS data, and xylose-bound simulations fit to xylose-bound HDX-MS data, etc.) in a 50:50 mixture of OF:IF conformational populations. HDXer was then applied to each candidate ensemble to obtain a reweighted ensemble with improved correlation to experimental data (**Figure S11**). We applied a *γ* value (tightness of fit) corresponding to a reweighting apparent work *W*_*app*_ of ∼ 5 kJ/mol to initial ensemble structures to avoid overfitting (**Figure S12**). As such, the same *W*_*app*_ was then assigned in each individual reweighting, ensuring equivalent bias was applied to each initial ensemble of structures, and therefore that results for each state were comparable^19^.

Consistent with experimental HDX-MS data, binding of xylose and glucose resulted in an OF favoured conformational equilibrium shift compared to the apo state. Interestingly, the shift is significantly more prominent for the glucose-bound state for which the final reweighting ensemble results in a 55.29% OF population (**Figure 6a & Table S3**). Strikingly, phloretin-bound structures exhibit a dramatically different conformational landscape to xylose/glucose bound structures. The reweighted phloretin-bound ensemble consisted of 72.70% IF structures, consistent with the original visual interpretation of the conformational fingerprint observed in the experimental HDX-MS data (**Figure 3b**). Interestingly and consistently with HDX-MS experiments, phloridzin-bound structures suggest a different fractional population (55.44% OF) in the final reweighting ensemble compared to phloretin-bound structures. Whereas this may be reflective of a more occluded like structure as suggested by the HDX-MS fingerprint (**Figure 3b**), we do not have fully occluded structures in the simulation ensembles to confirm that.

**Figure 6.**
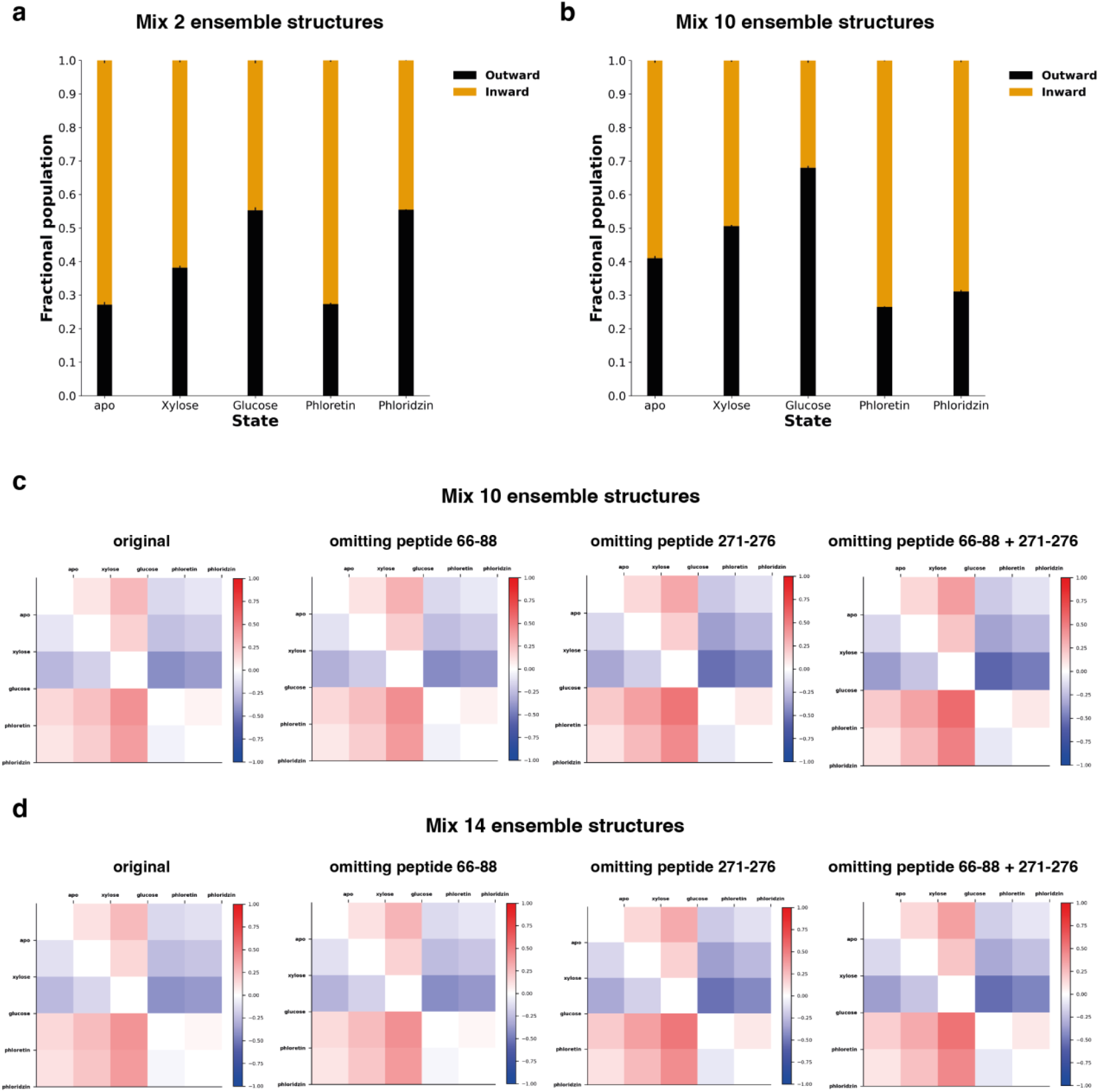
Ensemble reweighting of XylE structures in apo and ligand-bound states. **(a)** The fractional population of the final reweighting ensemble with a mixed 2 (OF/IF) ensemble structure fitted to each state-specific experimental HDX-MS data (apo, xylose-, glucose- phloretin-, phloridzin-bound state). **(b)** Fractional population of the final reweighting ensemble with a mixed 10 (OF/IF) ensemble fitted to each experimental HDX-MS data (apo, xylose-, glucose-, phloretin- and phloridzin-bound state). Standard deviations for each fitting are plotted as errors (n=3). **(c,d)** Heatmap of relative fraction difference between states. Red indicates more OF, and blue indicates more IF.

Next, we mixed all 10 ensemble structures covering 5 protein states, each with 2 ensembles in both OF and IF conformations. Noticeably, mixing all 10 ensemble structures to fit experimental data results in a slightly better agreement to target data for all protein states (**Table S4**), implying some conformations in the “alternate state” structures are better at describing experiment data. However, the overall trend of percentage population for apo, xylose-, glucose- and phloretin-bound structures remains the same compared to the previous ensemble with only “state-specific” structures. Surprisingly, reweighting of phloridzin-bound structures leads to a different percentage population compared to glucose-bound structures, but largely similar population to that observed in phloretin bound (73.51% IF) structures, with 68.90% IF conformers in the final reweighting ensemble (**Figure 6b & Table S4**).

To investigate further the source of the discrepancy, we carried out additional reweighting experiments. Initially, we assessed the impact of experimental noise. RMSE values for each peptide segment were calculated from the ensemble mixture before and after reweighting by HDXer. Peptide 271-276 was identified as the most error-prone peptide across all states (**Table S5**). On top of that, due to an observed decrease and increase of deuterium uptake in different time points for phloretin-bound structures, peptide 66-88 was also included as an error-prone peptide to investigate how it affects the reweighting results using HDXer (**Figure S13**). We then carried out reweighting by omitting peptides 66-88 or 271-276 or both from the analysis. Our results suggest that the errors from these peptides are unlikely to impact the observations from the original dataset. Pairwise comparison of relative OF fraction population was plotted for apo and ligand-bound states (**Figure S14a & Figure S15**). The overall trend of difference in the conformational population remains the same across all tested conditions (original and peptides excluded) (**Figure 6c & Figure S14b**).

We then set out to assess the effect of sampling errors from insufficient conformational sampling. An additional four atomistic MD simulations of phloretin- and phloridzin-bound structures in both OF and IF were performed (**Figure S16**), adding up to a mixture of 14 ensemble structures for HDXer reweighting. We first checked the fractional populations after reweighting for previously generated ensemble structures, newly generated ensemble structures or both for phloretin and phloridzin separately (**Figure S14d**). No distinguishable difference was observed. We then repeated reweighting for the full set of 14 mixed ensemble structures under the same conditions. Interestingly, the mixed ensemble displayed a similar relative percentage population regardless of introducing additional ensemble structures or omitting peptides (**Figure 6d & Figure S14c & Table S6**). It is worth noting that our validation so far was carried out using the same Best and Vendruscolo predictive model^20^, and any inaccuracies related to the model will systematically be reflected across all the reweighting processes. Therefore, potential inaccuracies in the predictive model should not affect one state more than the other.

## Discussion

In summary, we have presented a detailed workflow for quantifying the conformational landscape of the sugar transporter XylE by integrative modelling with HDX-MS data and molecular simulations in three steps. Initially, we generated a wide-ranging candidate ensemble of potential XylE structures, encompassing OF and IF transporter states. Subsequently, HDX-MS experiments together with ensemble reweighting suggest that the GLUT inhibitors phloretin and phloridzin exhibit a substantially different mode of action to the endogenous ligands (xylose and glucose). The final reweighted structural ensembles, fitted to each experimental dataset independently, allow us to probe this mechanistic difference at the atomistic level.

Our HDX-MS results indicate that the binding of phloridzin causes overall protection to the protein while its aglycone, phloretin shifts the protein conformational equilibrium towards IF. By combining MD simulations with a *post hoc* ensemble reweighting approach, we were able to quantify and visualise conformational changes incurred upon inhibitor binding. Overall, our results point to the hypothesis that phloridzin-bound structures cannot be assumed as an ensemble of structures occupying simple OF and IF representations. This hypothesis is supported by occluded HDX-MS pattern (phloridzin-bound structure vs apo) and in-between (glucose and phloretin) percentage population after reweighting with only using “state-specific” OF and IF ensemble structures. Our predictive models further revealed the different conformational changes of inhibitor binding and pointed to the key residue contributions for inhibitor binding (Gln168, Gln175 and Gln415). Overall, this interplay between different ligands and proteins offers an entirely new view of the mechanism of action for GLUT inhibitor binding.

Furthermore, by fitting a mixture of XylE WT and G58W ensemble structures to XylE WT experimental data, we observed differences in the final reweighted ensemble between previously published data, which led to 95.6% IF, and newly generated data, which led to 72.9% IF. It is worth noting that, as previous HDX-MS data were not associated with a MaxD control, we back-exchange corrected them with a newly acquired MaxD control to enable the data for reweighting. In contrast, new HDX-MS data and associated MaxD were performed at the same time. By comparing these two sets of data, we observed consistently more deuterium uptake in newly generated data than in previously published data, however, the deuterium incorporation level in two MaxD showed only negligible difference (**Table S7**), indicating that the newly performed MaxD did not perfectly match previously acquired data. This introduced a small but consistent bias in the calculated relative fractional uptake (%) for previously published data and is responsible for the discrepancy in the reweighted ensemble. Our results demonstrated the capability of our integrative approach to accurately reflect conformational effects and capture the differences in the target experimental data.

## Conclusion

In this study, we also examined the source of errors by eliminating possible error-prone peptides and introducing additional ensemble structures. This work suggested the robustness, consistency and accuracy of our combined experimental and computational strategy.

Concluding, this combinatorial workflow and robust strategy offer a new way to infer dynamic information from HDX-MS experiments. It complements and improves existing methods to interpret ensemble-averaged experimental observables and adds crucial high-resolution detail to our inferences of the MFS transporter mechanism from HDX-MS experiments. More importantly, the methodology could readily be expanded to any proteins of interest for which HDX-MS experiments are available.

## Methods

### XylE expression and purification

XylE was overexpressed in *E. coli* BL21-AI (DE3) (Invitrogen), transformed with the XylE WT gene and cloned in the (30 µg/ml) kanamycin-resistant pET28-a plasmid (Novagen) modified with a C-terminal 10-histidine tag. Bacteria were grown in 6 baffled flasks each containing 1 L of LB media at 37 °C 220 rpm to an OD_600_ of 0.8. Expression was induced with 1 mM Isopropy-b-D-1-thiogalactopyranoside (IPTG) and 0.1% (w/v) L-arabinose, and growth continued until the value of OD_600_ was flat. The cells were harvested by centrifugation, washed in 200 mL phosphate-buffered saline (PBS) buffer and centrifuged again for 20 min at 4,200 rpm in a Beckman JLA-16.250 rotor. The pellet was then resuspended in 50 mL PBS buffer with 10 mM β-mercaptoethanol and 1 cOmplete protease inhibitor tablet and frozen at −70 °C before purification. Cells were defrosted and incubated with 1.5 µL benzonase nuclease (ThermoFisher) for 10 min at room temperature before passing through a constant cell disrupter at 25 kPsi and 4 °C. Then the ice-chilled membranes were isolated by ultracentrifugation for 30 minutes at 38,000 rpm in a Beckman Ti45 rotor, 4 °C. Membrane pellets were solubilised for 2 hours with mixing in solubilisation buffer [50 mM sodium phosphate pH 7.4, 200 mM NaCl, 10% (v/v) glycerol, 20 mM imidazole, 10 mM β-mercaptoethanol, and 2% n-Dodecyl β-D-maltoside (β-DDM, Anatrace), 0.1 mM phenylmethylsulfonyl fluoride (PMSF) and EDTA free protease inhibitor tablet (Roche)] at 4 °C. The protein solution was then isolated by centrifugation for 30 min at 38,000 rpm in a Beckman Ti70 rotor to remove insoluble material. The supernatant was filtered using 0.45 µm filter and applied to a Ni-NTA column equilibrated in 96% SEC purification buffer [50 mM sodium phosphate pH 7.4, 10% (v/v) glycerol, 2 mM β-mercaptoethanol, and 0.05% β-DDM (Anatrace), 0.1 mM phenylmethylsulfonyl fluoride] and 4% elution buffer [50 mM sodium phosphate pH 7.4, 500 mM imidazole, 10% (v/v) glycerol, 10 mM β-mercaptoethanol, 0.1 mM phenylmethylsulfonyl fluoride (PMSF) and 0.05% β-DDM (Anatrace)]. The bound protein was washed with 50 mL 85% SEC purification buffer – 15% elution buffer and eluted with 2 mL of 100% elution buffer. The eluate was collected for further size exclusion chromatography (SEC). The SEC purification was conducted with a Superdex 16/600 GL SEC column, equilibrated with SEC purification buffer. The elution fraction containing XylE was collected and concentrated with a Vivaspin concentrator (100 kDa cutoff) (**Figure S17**). The samples were either flash frozen and kept at −70 °C until use or used directly for HDX-MS experiments.

### Ligand solution preparation

D-xylose was purchased from Santa Cruz Biotechnology. D-glucose (>99.5%) was bought from SIGMA life science. Phloretin and phloridzin were purchased from Merck life science. All ligands were first dissolved in 100% DMSO as phloretin and phloridzin were insoluble in pure water. Subsequently, the solubility of ligands in 10% and 1% DMSO was tested separately to ensure the solubility in equilibration and deuterium labelling conditions (10 times dilution).

### Peptide identification

Peptide identification of XylE was performed by liquid chromatography-tandem mass spectrometry (LC-MS/MS) analysis using a Synapt G2-Si HDMS coupled to nanoACQUITY UPLC (Waters). XylE in detergent micelles were prepared at a concentration of around 15 µM using 100 kDa cutoff Vivaspin concentrators. The protein sample (2.25 μL) was incubated with 0.25 μL DMSO and 22.5 μL equilibration buffer E (10 mM potassium phosphate in H_2_O pH 7.0) and added with 25 μL ice-cold buffer Q (100mM potassium phosphate in formic acid pH 2.5), simulating labelling conditions. Then, protein samples were injected into the LC system and digested online with a self-packed pepsin column at 20 °C. Peptides were trapped for 3 min using an Acquity BEH C18 1.7 μm VANGUARD pre-column at a 200 μL/min flow rate in solvent A (0.1% formic acid in HPLC water, pH 2.5) before eluted to an Acquity UPLC BEH C18 1.7 μm analytical column with a linear gradient (8-40%) of solvent B (0.1% formic acid in acetonitrile) at a flow rate of 40 μL/min. All trapping and chromatography were kept at 0 °C. Then, peptides went through electrospray ionization in positive ion mode and were analysed using a Synapt G2-Si mass spectrometer (Waters) within the mass range 100-2000 m/z. Leucine Enkephalin was applied for mass accuracy correction and sodium iodide was used as calibration for the mass spectrometer. MS^E^ data were collected by fragmenting with a 20-30 V trap collision energy ramp. Five protein injections were performed for peptide identification. To minimize peptide carryover, the pepsin column was washed once between injections using a pepsin wash solution (1.5 M Gu-HCl, 4% (v/v) MeOH, 0.8% (v/v) formic acid) and an LC run with a sawtooth gradient was conducted between each sample injection to wash the analytical segment.

### Continuous deuterium labelling of XylE

XylE in detergent micelles were prepared at a concentration of around 15 µM using a 100 kDa cutoff Vivaspin concentrator. Protein and ligands (xylose, glucose, phloretin and phloridzin solubilised in DMSO) were incubated at ratio enabling about 90% binding occupancy (**Table S1**) according to **Equation 2**, for an estimation of the fraction of the protein in bound forms (*f*_*B*_) from the starting concentration of protein ([*P*_0_]) and ligand ([*L*_0_]) with binding affinity (*K*_*d*_). To enable HDX, the protein alone and the protein incubated with ligands were 10-fold diluted with the deuterium labelling buffer L (10 mM potassium phosphate buffer in D_2_O, pD 7.0) to initiate the exchange reaction. An aliquot of 25 μL (containing 2.5 μL of protein) was withdrawn from the labelling mixture at various time points (30 s, 5 min, and 30 min) and quenched 1:1 with 25 μL ice-cold buffer Q. After quenching, samples were left on ice for 10 s before flash freezing them in liquid nitrogen and kept at −70 °C until LC-MS analysis. Technical triplicates were performed at every time point and condition studied.

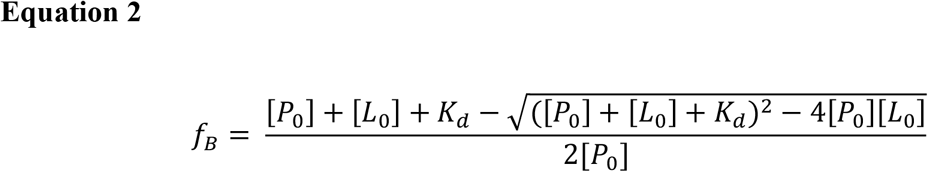

### LC-MS analysis of HDX samples

Frozen quenched samples were rapidly thawed and injected onto a Waters nanoACQUITY UPLC system with all trapping and chromatography elements set at 0 °C. Then, the protein was digested with a self-packed pepsin column with pepsin immobilized on agarose resin beads at 20 °C. The pepsin column was washed 4 times between injections using the pepsin wash solution. Two sawtooth runs were done between each sample run to reduce peptide carry-over. Peptides were trapped for 3 min using an Acquity BEH C18 1.7 μm VANGUARD pre-column at a 200 μL/min flow rate in solvent A before eluted to an Acquity UPLC BEH C18 1.7 μm analytical column with a linear gradient (8-40%) of solvent B at a flow rate of 40 μL/min. Then peptides went through electrospray ionization progress in a positive ion mode using Synapt G2-Si mass spectrometer (Waters). The mass range for MS was m/z 100 to 2000 in positive ion mode. Leucine Enkephalin was applied for mass accuracy correction and sodium iodide was used as calibration for the mass spectrometer. MS data were collected by a 20-30 V trap collision energy ramp. All the isotope labelling time points were performed in triplicate.

### Maximally deuterated control

XylE (in DDM micelles) was concentrated to 20 µM in 50 mM sodium phosphate, 100 mM NaCl, 10% (v/v) glycerol, 1 mM β-mercaptoethanol, 0.02% β-DDM pH 7.4. 2 µL of protein sample were diluted in solvent A and repeatedly injected into the LC system. Following online pepsin digestion, peptides eluting from the pepsin column were collected. Three injections were repeated, the eluted fractions mixed together and then split again into three samples. Peptides were lyophilized on a freeze dryer. Each aliquot of lyophilized peptides was incubated with 2.5 μL of buffer E and diluted 10-fold with 6 M deuterated urea in D_2_O, resulting in identical deuterium fraction as for labelled proteins. The reaction was allowed to proceed for 3 hr, before quenching with 25 μL buffer Q. Quenched samples were kept on ice for 10 s, then flash-frozen and stored at −70 °C until LC-MS analysis. Analysis of the maximally deuterated samples was carried with the pepsin column replaced with a metal union and LC-MS analysis was conducted as for standard deuterated samples.

### HDX data evaluation and statistical analysis

Peptide identification was performed by processing the acquired MS^E^ data with PLGS (ProteinLynx Global Server 2.5.1, Waters). DynamX v.3.0 (Waters) was used to further filter peptides with 0.25 fragments per amino acid and identified in at least 4 out of 5 acquired MS/MS files. To obtain a final peptide map, spectra were further visually inspected in DynamX to exclude peptides of insufficient quality or misidentified. Then all the MS data including undeuterated references and deuterated samples were processed by DynamX for calculation of deuterium incorporation. Peptides with a statistically significant difference in HDX were determined by in-house Deuteros 2.0 software^21^ with a hybrid significance test with a 99% confidence interval.

### Preparation of XylE structures for MD simulations

Initial protein structures were downloaded from the Protein Data Bank (PDB) (https://www.rcsb.org/) with PDB ID 4GBY for outward open partially occluded conformation and 4JA4 for IF conformation. Missing residues in the IF XylE structure (built from 4JA4 chain A) were rebuilt using Modeller 9.24^22^. The 4JA4 sequence with gaps was first aligned to that of 4GBY so that the same residues (5-479) were present in both final structures. 10 initial models of the missing loop sequences were generated, using 4GBY as a template for the loop structures and 4JA4 as a template for the remaining parts of the protein. For each initial model, residues in the loops were remodelled using the loop model functionality, to give 5 alternate loop structures. From the initial model with the lowest molpdf score and DOPE score, the loop conformation with the lowest subsequent molpdf score and the DOPE score was chosen as the initial IF structure for MD simulations. Restrictions using cysteine cross-linking (A152C/S396C or V35C/E302C) were applied to the protein in separate simulations. Site-directed mutations were introduced into structures using PyMol with structural-guided identification of loci for cysteine mutations. An intracellular cysteine pair A152C and S396C was introduced in the outward-open partially occluded XylE structure (PDB 4GBY), and an extracellular cysteine pair (V35C and E302C) in the inward-open XylE structure (PDB 4JA4), with a sulfur-sulfur distance of 2.1 Å in both structures. These restrictions have been previously validated experimentally to lock the protein in OF or IF states^13^. Possible Asn/Gln/His sidechain flips were checked by uploading to MolProbity^23^, followed by manual visual inspection on PyMol where sidechain flips were determined in **Table S8**.

### Setup of XylE cross-linked states in POPE bilayer system

The construction of XylE structures with cross-linked mutations in the membrane system was prepared by CHARMM-GUI^24, 25^. Terminal group patches were set to ACE and CT3 for the N-terminus and C-terminus, respectively. POPE lipids were selected at a 1:1 ratio for bilayer construction, the final upper leaflet number is 177 and the lower leaflet is 178. The protein was aligned along the z-axis and the bilayer in the xy-plane. Structures were oriented in the membrane by the PPM web server^26^. The system was then solvated with 25124 TIP3P water molecules, resulting in a cuboid periodic box of 110 Å x 110 Å x 112 Å. Na^+^ and Cl^−^ ions were added to neutralise the system charge and to create an ionic atmosphere of 150 mM NaCl. Protonation states of the titratable residues were assigned by the H++ server^27^ (http://biophysics.cs.vt.edu/). Specifically, E206 was protonated, consistent with the previous study, and all remaining residues were assigned to their standard protonation states at pH 7.0.

### MD simulations of XylE cross-linked systems

Protein simulations were performed in Gromacs (2020.1) on the research computing facility at King’s College London, Rosalind (https://rosalind.kcl.ac.uk/), Jade (https://www.jade.ac.uk/) and HPC cluster Gravity (https://apps.nms.kcl.ac.uk/wiki/gravity/). The whole system was first energy-minimized for 5000 steps without any positional or dihedral restraint using a steepest-descent method to relax any steric clashes. The system was then equilibrated to 303.15 K and 1 bar pressure over a 6-stage equilibration protocol with a total simulation time of 100 ns. Positional restraints on the protein and lipids and dihedral restraints on lipid head groups were added to maintain geometry and chirality as the box size was equilibrated. The force constants of positional or dihedral restraints are specified in **Table S9**. Restraints were gradually decreased across the first 5 stages, allowing the protein sidechains to move first, then reducing the force constant on the backbone restraints, and finally only the protein Cα atoms were restrained in step 5 (**Table S9**). In the final stage – before production runs, an unbiased 60 ns NPT equilibration was carried out to prepare the system for production MD (**Figure S18**). A 100 ns production run was then performed at constant temperature (303.15 K) and constant pressure (1 bar) using a Nose-Hoover thermostat and Parrinello-Rahman barastat^28^ in a timestep of 2 fs. After completion of production runs, trajectories were read in MDTraj (https://www.mdtraj.org/1.9.5/index.html) and imported into Visual Molecular Dynamics (VMD v1.9.4) (https://www.ks.uiuc.edu/Research/vmd/) for visual inspection.

### Extraction of representative MD structures

Representative protein structures from apo-state MD simulations were extracted using the DBSCAN (Density-based spatial clustering of application with noise) clustering method based on the chi1 dihedral value of 10 binding site residues (Phe24, Asn294, Asn325, Gln168, Gln175, Gln288, Gln289, Gln415, Trp392, Trp416). Three replicates of the simulations for OF/IF structures were converted in one trajectory with 12003 frames in total. Dihedral angles of the aforementioned residues were firstly calculated using gmx tools (gmx angle). Each dihedral angle was then represented by the sine and cosine of the angle, which turns the angles into a point on the unit circle. The distances between points (represented as the chord length) were clustered with minPts set as 60 and epsilon ranging from 0.50 to 1.35 at 0.05 intervals. Each epsilon was assessed with a Silhouette score to choose an optimal cluster size and number of clusters. Epsilon values of 0.55 for both OF/IF structures were chosen. The mean value of the distance between points in each cluster was calculated, and the frame closest to this mean value was then extracted as the representative frame. In total, for OF conformation, 19 representative MD structures were chosen, whereas 11 were selected for IF conformation.

### XylE-ligand binding mode prediction

Autodock Vina was used to predict the mode of binding of the ligands discussed in this study. Before docking, the structures of the GLUT inhibitors (phloretin and phloridzin) were downloaded from PubChem^29^. Xylose and glucose were extracted from corresponding crystal structures from Protein Data Bank (4GBY; 4GBZ). Both the representative protein frames extracted from MD simulation and original XylE crystal structures (4GBY; 4GBZ; 4JA4) were used as receptors. Receptor structures were prepared using Autodock Tools. The initial ligand and protein structure file (pdb) including the desired positioning of hydrogens was created with Autodock Tools. For the ligand structure, a torsion tree (a list of the rotatable torsions ready for sampling) was created for a selection of rotatable bonds. The completed protonated and torsion-parameterised ligand and protonated protein (receptor) structures were saved as a pdbqt file. A grid box used to define the centre and size of each docking run was defined as a dimension of 22.5 x 22.5 x 30 Å with a box centre sitting at the CG atom of residue Trp392. For each receptor structure, a maximum of 9 possible poses of each ligand were generated.

### Dock xylose/glucose to crystal structures

To test the capability of Autodock Vina, it was applied to dock xylose/glucose back to crystal protein structures (4GBY; 4GBZ). Xylose/glucose docking was tested with two levels of ‘exhaustiveness’ (conformational sampling) specified in the configuration file, with the exhaustiveness value set at 8 and 96. Docked poses for xylose/glucose into crystal structure (4GBY/4GBZ) were converted from .pdbqt to .pdb file using Open Babel^30^ for further analysis. Each docked pose generated from several distinct representative protein structures was first aligned to the crystal structures 4GBY and 4JA4 respectively using a custom VMD script. Given that the ligand atom order from original crystal structures is different from the docked poses, the atom order of xylose and glucose in crystal structure was edited to match exactly with the docked poses achieved from Autodock Vina using a custom python script. Both xylose and glucose were successfully docked into the ligand-binding pocket of XylE since the top-scoring docking pose showed an almost identical binding pattern to the crystal structure (**Figure S19**).

### Dock xylose/glucose to representative MD structures

Rigid docking (receptor structure treated as rigid-body) was carried out for xylose/glucose into representative protein structures (OF: 19; IF: 11) from MD simulations. For each ligand, up to 171 (OF) and 108 (IF), possible protein-ligand structures were generated, covering a range of different binding poses. The RMSD of the conformation between docked poses of xylose/glucose and their poses inside crystal structure was calculated using a custom RMSD calculation python script, each pose was assigned with an RMSD value. The distribution of the RMSD values between docked poses and ligand from crystal structure was plotted using a custom python script (**Figure S20a**). Multiple docking poses with high variance were then clustered using DBSCAN and hierarchical clustering methods. The top pose from the binding pocket cluster using DBSCAN and hierarchical clustering methods is identical. RMSD value between the top pose and crystal ligand structure was calculated as 1.626 Å (**Figure S20b**).

### Dock phloretin/phloridzin to representative MD structures

Two GLUT inhibitors (phloretin and phloridzin) were docked into crosslinked XylE OF and IF structures extracted from MD simulations using the same protocol for xylose/glucose. Docked poses for each inhibitor were clustered using Principal Component Analysis (PCA) based on atomic coordinates to identify representative structures of each binding pose, and visual inspection ensured that the selected representative structures were within the XylE binding pocket (**Figure S21**). One top pose with the highest binding affinity and one cluster centroid pose from clusters within the binding pocket for each inhibitor were selected as the initial XylE-inhibitor bound structures for short MD simulations (100 ns).

### Ligand parameterization of ligand molecules

Force field parameters for small molecules are not covered by the CHARMM36m bimolecular force field. The CHARMM General Force Field program (CGenFF)^31–33^ was used to perform atom typing and assignment of parameters for phloretin and phloridzin structures. Quantum-Mechanical (QM) calculations were carried out using Gaussian 09^34^ to validate and optimize parameters that exhibited high ‘penalty scores’ (>10) from CgenFF. A detailed parametrization protocol was followed using a plugin for the VMD software package known as Force Field Toolkit (ffTK)^35^ to make sure the parameters were fitted to data from high-level QM calculations. Conformations of the ligand molecules with minimum energy were obtained from PubChem. An initial set of parameters was assigned with the CGenFF server, creating a parameter file (.str) containing the assigned parameters beyond those in the normal CGenFF parameter file (.prm). A Gaussian input file for geometry optimisation at MP2/6-31g* level was created using the “Opt. geometry” tab in ffTK, and the initial molecular conformation was then optimised and saved as a QM geometry optimised conformation (pdb file) for subsequent charge and dihedral optimisation. QM optimization of the charge was carried out on Gaussian 09 where input files were generated via ffTK using the “Water Int” tab, in which water molecules were oriented to interact with functional groups in the molecule^32^. The orientation of each interacting water molecule depends upon whether the nearby atom is an H-bond donor or an H-bond acceptor based on their chemical environment. H-bonding interactions are predominantly based on electrostatics in molecular mechanical force fields. Positively charged atoms prefer to interact with the water oxygen atoms as H-bond donors while negatively charged atoms will interact with the water hydrogen atoms as H-bond acceptors, and carbons are assigned as both H-bond donors and H-bond acceptors in two Gaussian input files. Gaussian calculations were performed at the HF/6-31g* level for water interactions. Long-distance (> 4 Å) interaction data was removed from fitting QM calculation to MM data in the charge optimisation process, where water molecules fly off during optimisation due to steric clashes or unfavourable interactions. The target Gaussian water interaction data was read in ffTK via “Opt.Charges” tab, the Tolerance is set as default (0.005) which determines the convergence criterion of the fitting process. The Distance and Dipole weights, determining the relative importance of matching MM Distance and Dipole to QM data, were set at a relative weight of 2:1:1 Energy:Distance: Dipole in which the Distance and Dipole weighting were each set to 0.5. After the first iteration of optimisation, the charges were re-optimised at a smaller value of Tolerance (e.g., 0.001), the iterations were repeated until the final charges no longer change from step to next and then saved as an updated parameter file with the new charges. Like charge optimisation, dihedral optimisation requires computing the target data from high-level QM calculations first, then fitting MM dihedral parameters to target QM data. Initial dihedrals within the molecule of interest can be identified automatically in ffTK. In the Gaussian files, a torsional scan was performed by rotating each dihedral angle in steps until it rotates 180 degrees; the remainder of the molecule is allowed to relax at each step, to isolate the contribution of the energy associated with the dihedral of interest. Parameters with penalty scores under 10 from the initial CGenFF parameter assignment were removed from dihedral optimisation for fitting each dihedral at MM level to the QM target energy profile. After the first iteration of optimisation with tolerance set as 0.01 and energy cut-off at 10.0 kcal/mol under simulated annealing optimisation mode, further refinement of the dihedral parameters was performed until the fitting converge using energy profile visualization in ffTK^36^, and optimized dihedral parameters were updated in the final ligand parameter file for MD simulation runs.

### Protein-ligand complex simulation setup

The selected phloretin/phloridzin structures from docking and the associated representative apo-state MD protein structures were first merged together using a PyMol script. The membrane system was built using CHARMM-GUI membrane bilayer builder. Both protein and ligand were selected and uploaded into the CHARMM-GUI input generator. To create a CHARMM-GUI recognizable ligand parameter file, an initial rtf file was generated using the CHARMM-GUI ligand reader & modeler with the same mol2 file used for CGenFF. The charges in the topology (.rtf) file were manually updated with the optimised charged values from ffTK ligand parameterisation, and the ligand atom names and orders in the merged pdb file were manually updated to match those in the mol2 file. The final parameter file (.par) with optimised dihedral angles is recognizable to CHARMM-GUI, which is therefore directly uploaded using the “upload CHARMM top & par hetero chain” option together with updated topology (.rtf) file for the ligand. The POPE bilayer construction and system preparation for MD simulations was done in CHARMM-GUI as described in the previous section (preparation of XylE in the POPE system).

### Short MD simulations (100ns) of selected XylE-inhibitor bound structures

MD simulation protocol was similar to previously described for apo-state protein structures, except for some minor changes on restrains which are summarised in **Table S10**.

### Protein ligand interaction fingerprints

Protein ligand interaction fingerprints (PLIFs) were used to fully characterize the possible protein-inhibitor binding modes from MD simulations. Calculation of PLIFs was performed using the python package Open Drug Discovery Toolkit (ODDT)^37^ across the trajectories. Pairwise Tanimoto coefficient (**Equation 3**) was then calculated based on interaction fingerprints, where |*A* ∩ *B*| represents the number of the common ON bits presented in both string *A* and *B*, and |*A* ∪ *B*| is the union set, where the number of ON bits present in either string *A* or *B* is counted.

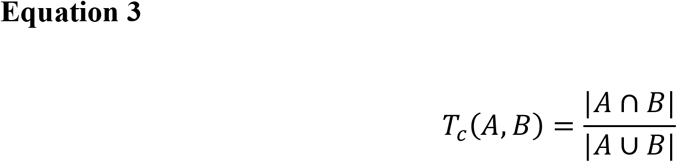

The distribution of the similarity score was first visualized in a heat map to identify converged clusters using a custom python script (**Figure S22**). Due to high variance, the hierarchical clustering method was then applied to identify clusters to select the most representative XylE-inhibitor bound structures from each short MD simulation of possible binding mode. Identified clusters were chosen for further analysis where all the following 3 requirements were met: (1). The size of the cluster was at least 10% of all MD frames. (2). The cluster contained at least one frame from trajectories initiated from each possible binding pose. (3). The majority of ligands inside the cluster were within 1.5 Å RMSD to the average structure (**Table S11**). The mean value of the distance between ligand conformation in each identified cluster was calculated, and the frame closest to this mean value was then extracted as the representative frame. Representative phloretin-and phloridzin-bound structures from identified clusters were extracted directly from previous short MD simulations (100 ns).

### Long MD simulation of XylE-ligand bound structures (1 µs)

Long MD simulations of phloretin- and phloridzin-bound structures were performed using the representative structure selected from the 100 ns simulation directly following the same MD simulation protocol described in the previous section (100 ns simulation protocol), only extending the production stage for 1 µs (**Table S10**). To compare phloretin- and phloridzin-bound structures with xylose- and glucose-bound structures, additional 1 µs MD simulations were performed. For xylose- and glucose-bound XylE OF structures, pdb structures (4GBY and 4GBZ) were used directly (substrate/inhibitor-bound protein structure only) following the simulation set-up preparation described in the previous section (Setup of XylE structures). Xylose- and glucose-bound IF structures were generated using Autodock Vina by docking xylose and glucose separately into IF apo-state crystal structure (4JA4). MD simulation protocols were used the same as phloretin- and phloridzin-bound structures.

### Analysis of 1 µs XylE-ligand bound simulations

Trajectory files were read using gmx tools, and 10 separate input trajectory files, each accounting for 100 ns, were concatenated with the gmx trjcat tool in sorted order while frames with identical time stamps were removed. The concatenated trajectory was further modified with gmx trjconv, where atoms were centred in the box and frames were saved at a time step of 250 ps. The RMSD of ligand structure was computed using gmx rms for each 1 µs trajectory with 4001 frames in total. Each frame from the trajectory was compared to the first frame with structures fitted onto protein transmembrane regions, RMSD calculation was done based on ligand heavy atoms. Output RMSD calculation results were read in a python script and plotted for visual inspection. Xylose in IF structure (4JA4) was dissociated over 1 µs simulation time, therefore XylE xylose-bound IF simulation was treated as XylE apo IF ensemble structures for further analysis. The number of hydrogen bond contacts between ligand (glucose, phloretin and phloridzin) and residues (Gln168, Gln175, and Gln415) were calculated using gmx hbond. Output from contact calculations (number of hydrogen bonds as a function of time) was read in a custom python script and plotted for visualization.

### MD simulations of XylE WT apo and G58W (1 µs)

Simulations of WT apo XylE were initiated from PDB entry 4JA4. Simulations of G58W XylE were initiated from 6N3I with the L315W mutation reverted to the wild-type leucine sequence. Following initial structure preparation, systems were protonated, minimised, and equilibrated following an identical 6-step protocol to the inhibitor-bound structures, followed by a 1 µs production simulation. Analysis of simulations was carried out as described in the aforementioned section.

### MD simulations data preparation to run HDXer

An atomistic ensemble of protein structures from MD trajectories was generated using gmx trjconv, with the system centred in the box and the centre of mass of molecules put in the box. For each 1 μs simulation, 4001 frames were saved at the interval of 250 ps. Peptide relative fractional uptakes were corrected for the deuterium uptake of the maximally deuterated control and then extracted using a python script for defined peptide segments for each deuterium labelling time. A list of peptide segments present in the target HDX-MS data was extracted using a custom python script from extracted deuterated fractions data.

### HDX data calculation using HDXer

Within HDXer, the Best and Vendruscolo method^20^ was used to estimate protection factors based on ensemble structures from MD simulations. The distance cut-offs were set as 0.65 nm and 0.24 nm to count heavy atom contact and hydrogen bonds between backbone amides and surrounding atoms including protein atoms, POPE lipids atoms, N terminus and bound ligand if applicable. The scaling factors are set as default to 0.35 for *β*_*c*_ and 2.0 for *β*_*H*_. Residue intrinsic exchange rates were calculated at the experimental conditions, in which pD is 7.0 and temperature is 293 K, with reference acid, base, and water catalysis parameters set to 1.62, 10.18 and −1.50 respectively^38^.

### Reweighting ensemble experiments

Ensemble reweighting was carried out using the output files of per-residue contacts and H-bonds created by HDX data calculation via HDXer. A mixed candidate ensemble, initially composed of 50% OF and 50% IF structures, was fitted using HDXer to each target HDX-MS dataset with absolute HDX-MS data. The *γ* value parameter in ensemble reweighting was first explored in a wide range from 1–10^−1^ to 9 x 10^2^. A “decision” plot was generated using a custom python script to investigate the effect of the choices of *γ* value upon the mean-square deviation (MSD) of the fitted data to target experimental data. In each decision plot, overfitting was classified as a rapid increase of the applied apparent work (*W*_*app*_) for limited improvement of MSD. Typically, this occurred at *W*_*app*_ over 5 kJ/mol. To adjust the initial ensemble without overfitting and allow equivalent comparison of ensembles reweighted to different datasets, we compared results at the exact *γ* value that corresponded as closely as possible to *W*_*app*_ of 5 kJ/mol. To investigate the effect of target experimental noise on final ensemble structures, a “leave one out” method was applied, in which peptide segments with high errors are systematically removed from the target data for each round of reweighting.

## Data availability

Data supporting the findings of this paper are available from the corresponding author upon reasonable request. All the deuterium uptake plots of the experiments presented for XylE are available on figshare data repository using the following link: (https://figshare.com/s/4f24c0aded2b5f51cd1d). Spectrometry proteomics data have been deposited to the ProteomeXchange Consortium via the PRIDE partner repository with the dataset identifier PXD034387.

## Author contributions

A.P. and R.T.B. conceived and designed the research; R.J. prepared protein samples; R.J., R.T.B, V.C. performed and analysed HDX-MS experiments; R.J., R.T.B. performed and analysed computational experiments; A.P. supervised the project; A.P. and R.J. wrote the paper. All authors commented on the final draft of the paper.

## Notes

The authors declare no competing interest.

## Acknowledgements

This work was supported by the Leverhulme Trust (RPG-2019-178) to A.P. A.P. is supported by an EPSRC Research Fellowship (EP/V011715/1). R.J. acknowledges receiving PhD studentship from King’s College London and China Scholarship Council.

## Notes

### Competing Interest Statement

The authors have declared no competing interest.

