## Supplementary information for "Integrative HDX-MS enables quantification of the conformational landscape of the sugar transporter XylE"

† Authors contributed equally

#### Supporting Figures

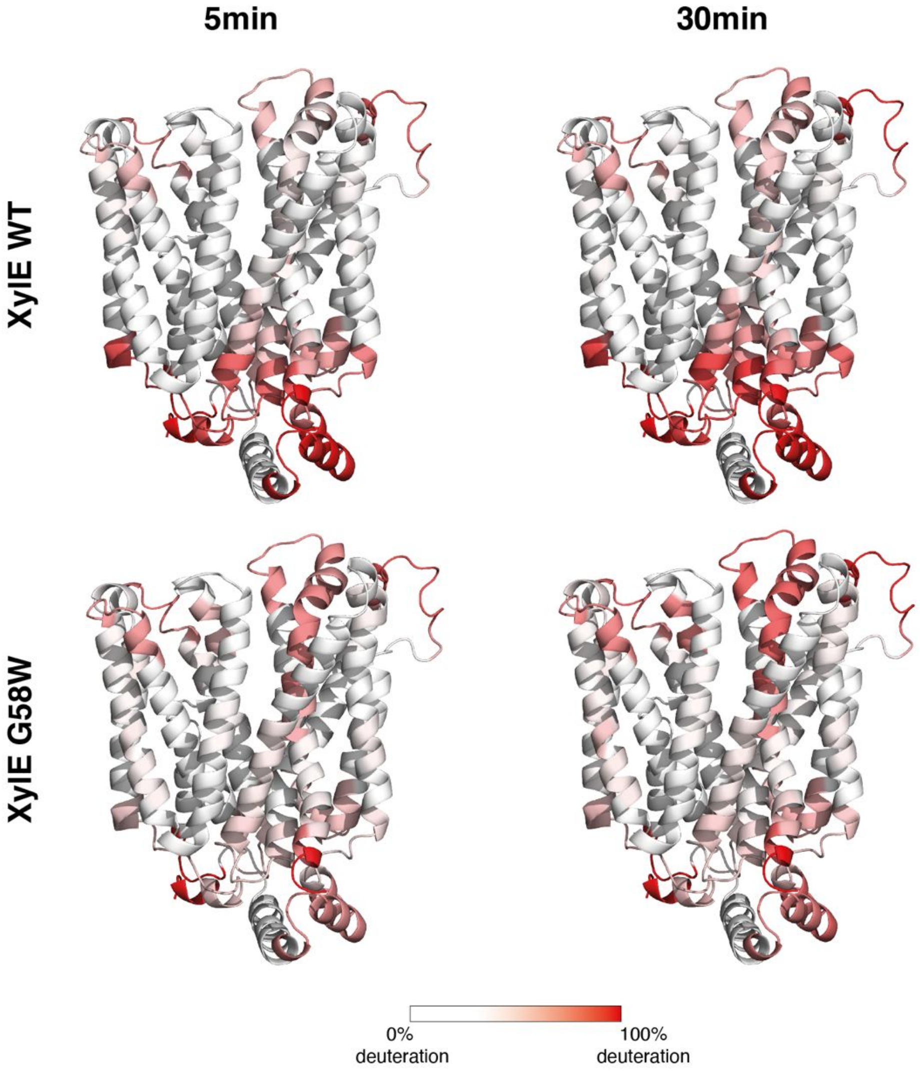

**Figure S1 Absolute deuteration of XylE WT and G58W.** Peptides with significant differences were analysed using the hybrid significance test incorporated in Deuteros 2.0[1] at a 99% confidence level.

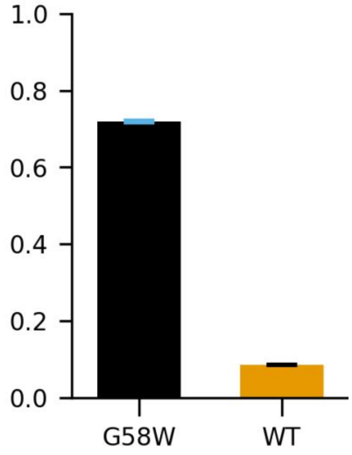

**Figure S2 Computational reweighting of a mixed OF/IF ensemble using a parametrized predictive model with $\beta_C$= 0.29, $\beta_H$= 3.9 still results in a clear separation of the structures present in each experimental dataset.** WT XylE remains mostly inward-facing, G58W mostly outward-facing.

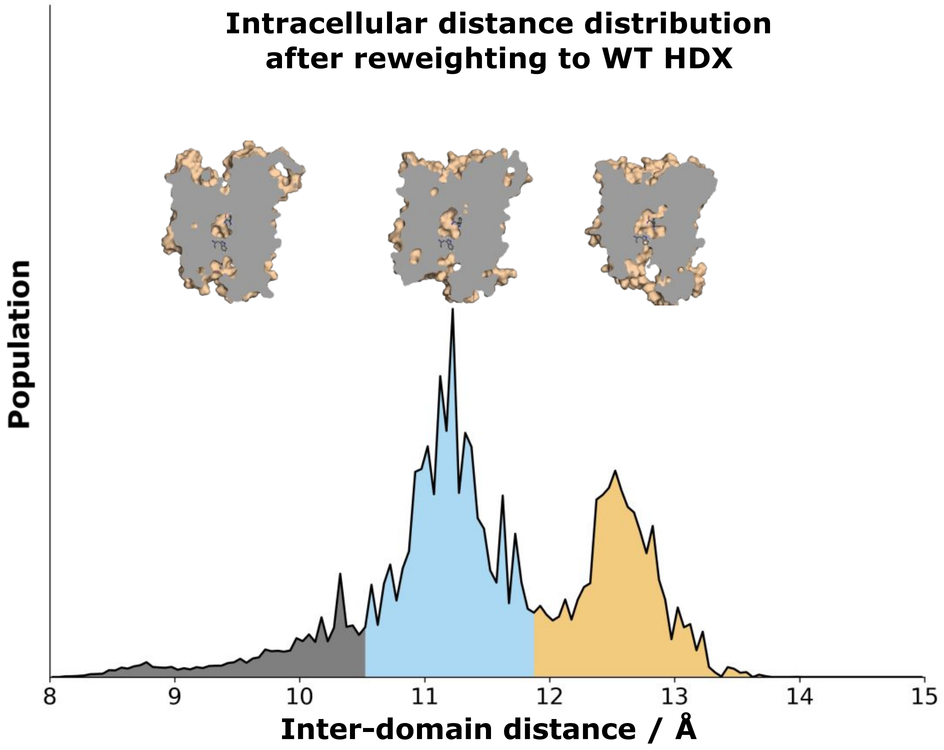

**Figure S3 Representative structures associated with final ensemble after reweighting to fit WT HDX-MS data.** The final ensemble contains substantial populations of both a partially-inward-open (blue cluster) and fully-inward-open (orange cluster) state. The extent of opening, defined by the inter-domain distance, is particularly correlated with the motion of intracellular loops 1-4, visible at the lower right of the structures.

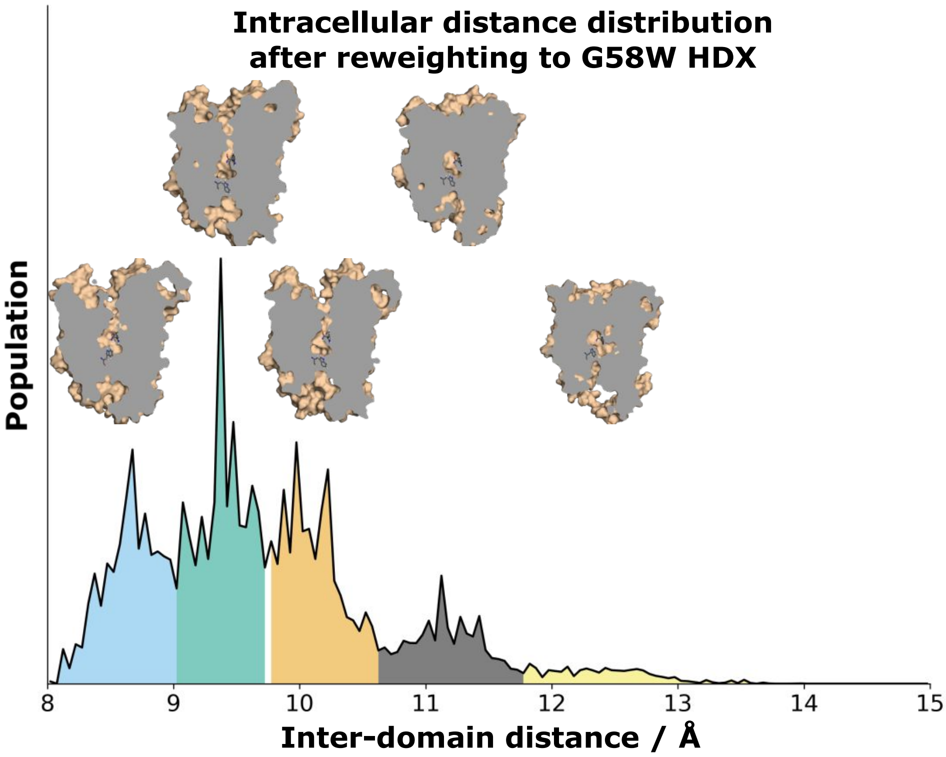

**Figure S4 Representative structures associated with final ensemble after reweighting to fit G58W HDX-MS data.** The final ensemble contains mostly outward-facing populations (blue, green, and orange clusters), with a minor population of a partially inward-open state (black cluster). The fully-inward-open (yellow cluster) state has a negligible population. Outward-facing structures also show flexibility in intracellular loops 1-4, which results in the broad distribution of intracellular inter-domain distances between 8.0 – 10.6 Å.

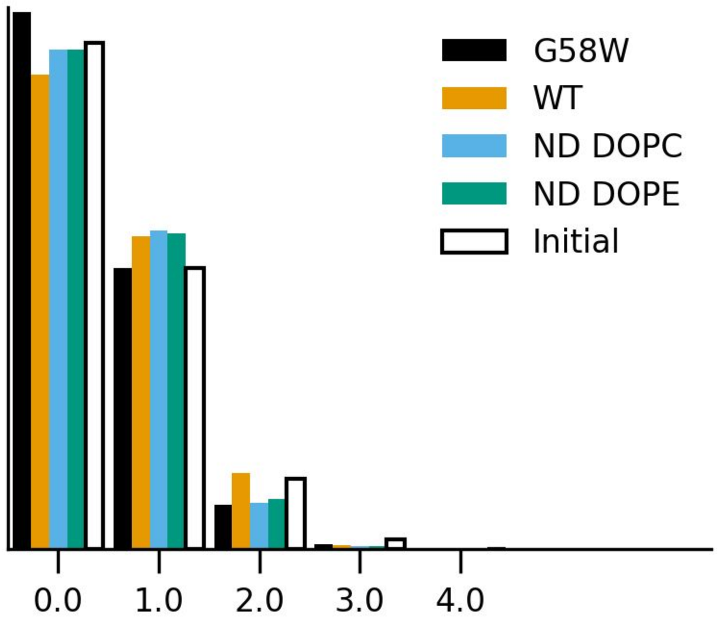

**Figure S5 Sum of lipid contacts between the PE headgroup and E153, D337, or E397, before (white) and after reweighting to G58W (black), WT (orange), DOPC nanodisc (blue) or DOPE nanodisc (green) HDX-MS data.**

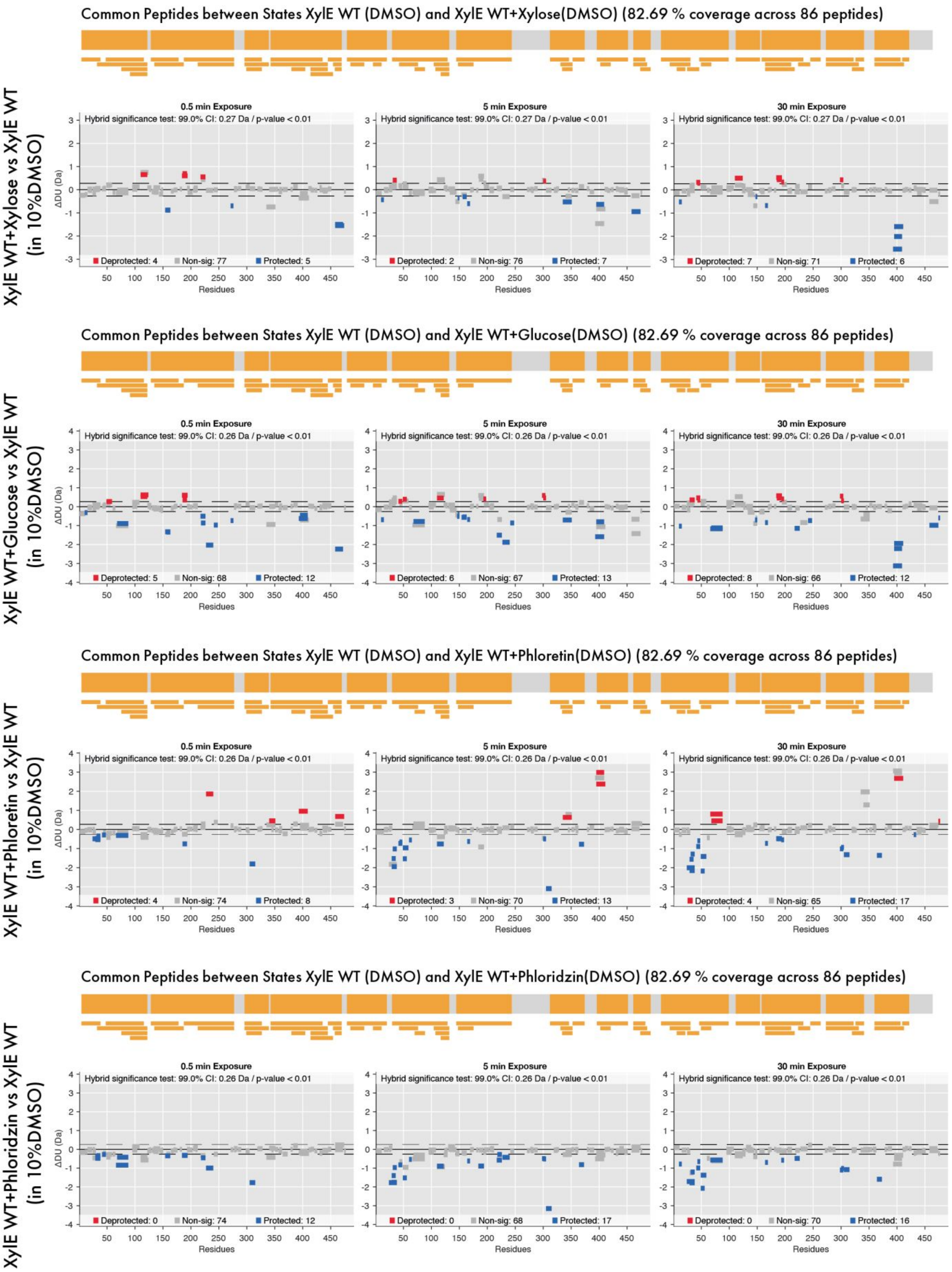

**Figure S6 Woods plots and common sequence coverage maps obtained from differential HDX-MS of XylE apo and ligand-bound states.** Each bar represents a single peptide with peptide length indicated by the bar length. Common peptides between two different protein states are indicated as orange.

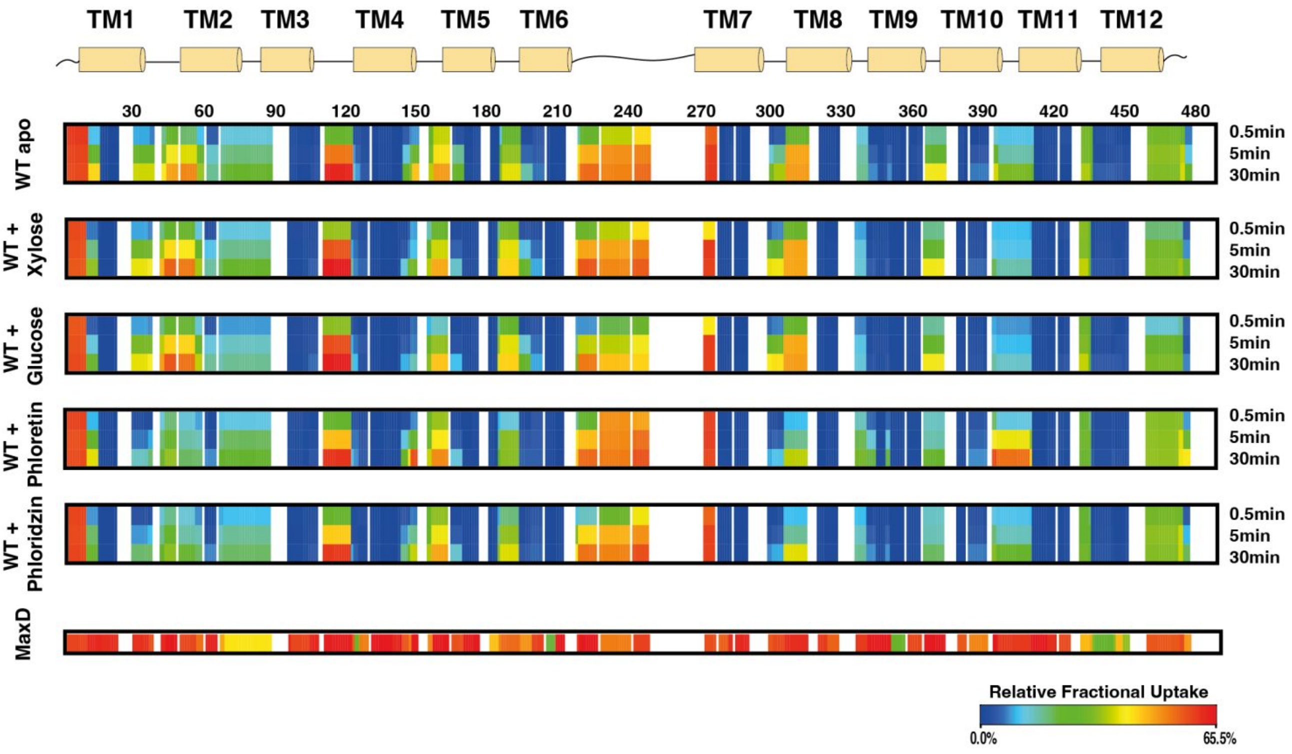

**Figure S7 Heatmap of all protein states in triplicates.** Protein states include XylE apo, WT + xylose, WT + glucose, WT + phloretin, WT + phloridzin and maximally deuterated control (MaxD).

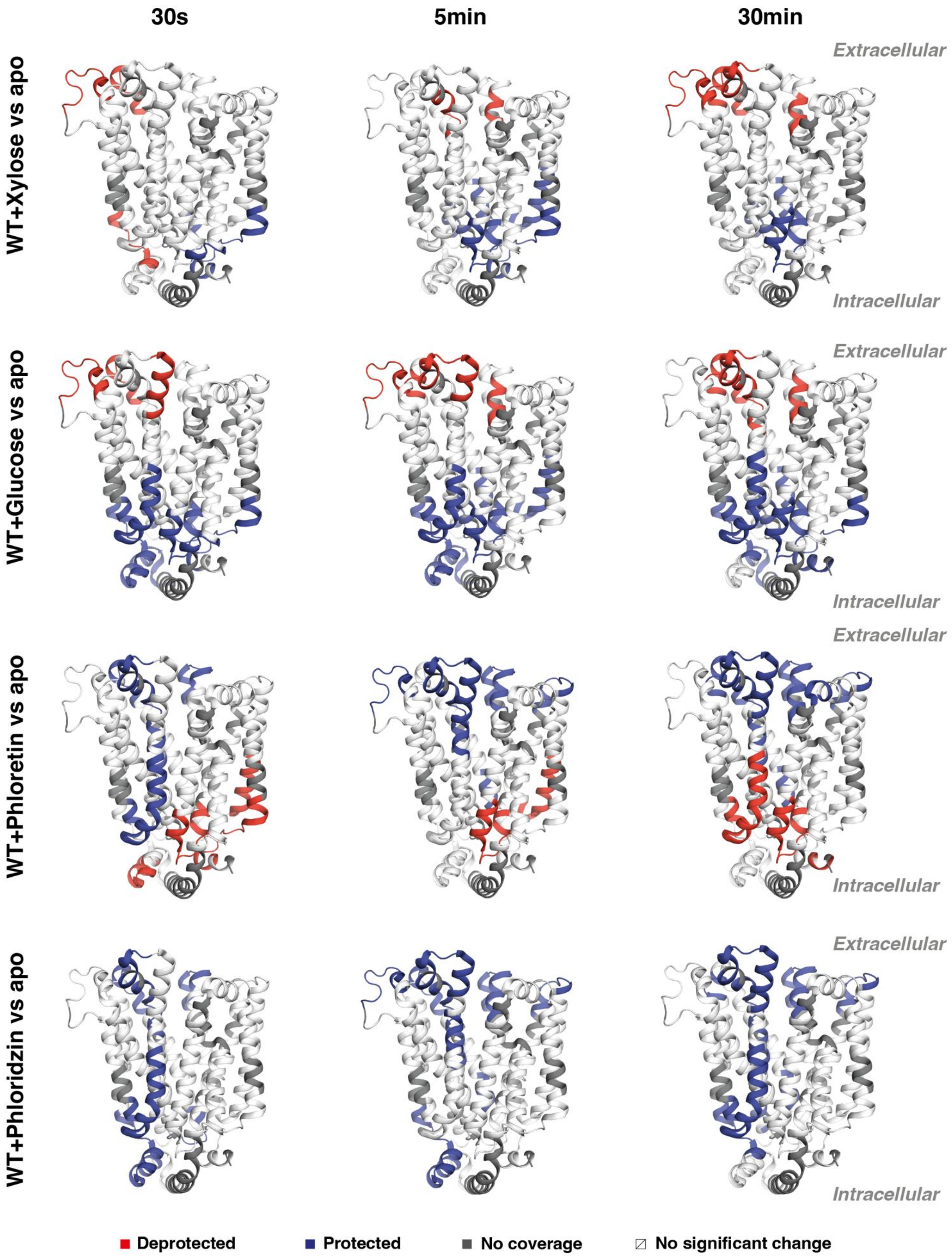

30s
5min
30min
WT+Xylose vs apo
WT+Glucose vs apo
WT+Phloretin vs apo
WT+Phloridzin vs apo
Extracellular
Intracellular
Extracellular
Intracellular
Extracellular
Intracellular
Extracellular
Intracellular
Deprotected
Protected
No coverage
No significant change

**Figure S8 Differential deuterium uptake plotted onto 3D protein structure (PDB: 4GBY) at 30s, 5min and 30min for all protein states.** Protein states include XylE apo, WT + xylose, WT + glucose, WT + phloretin and WT + phloridzin

**a**

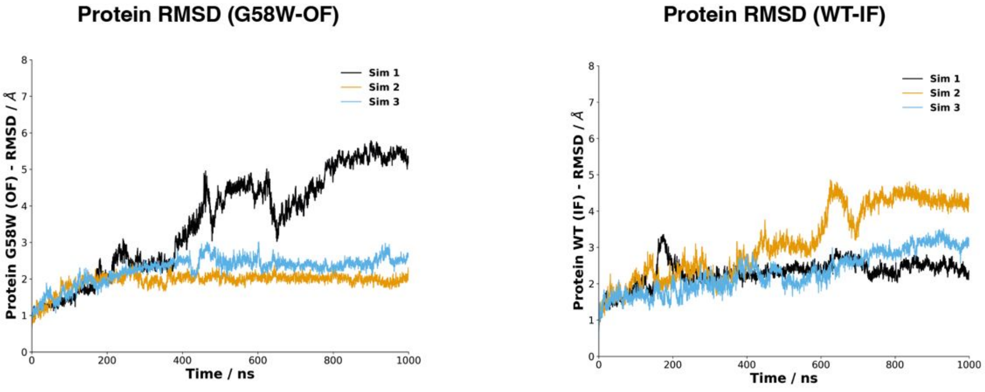

**b**

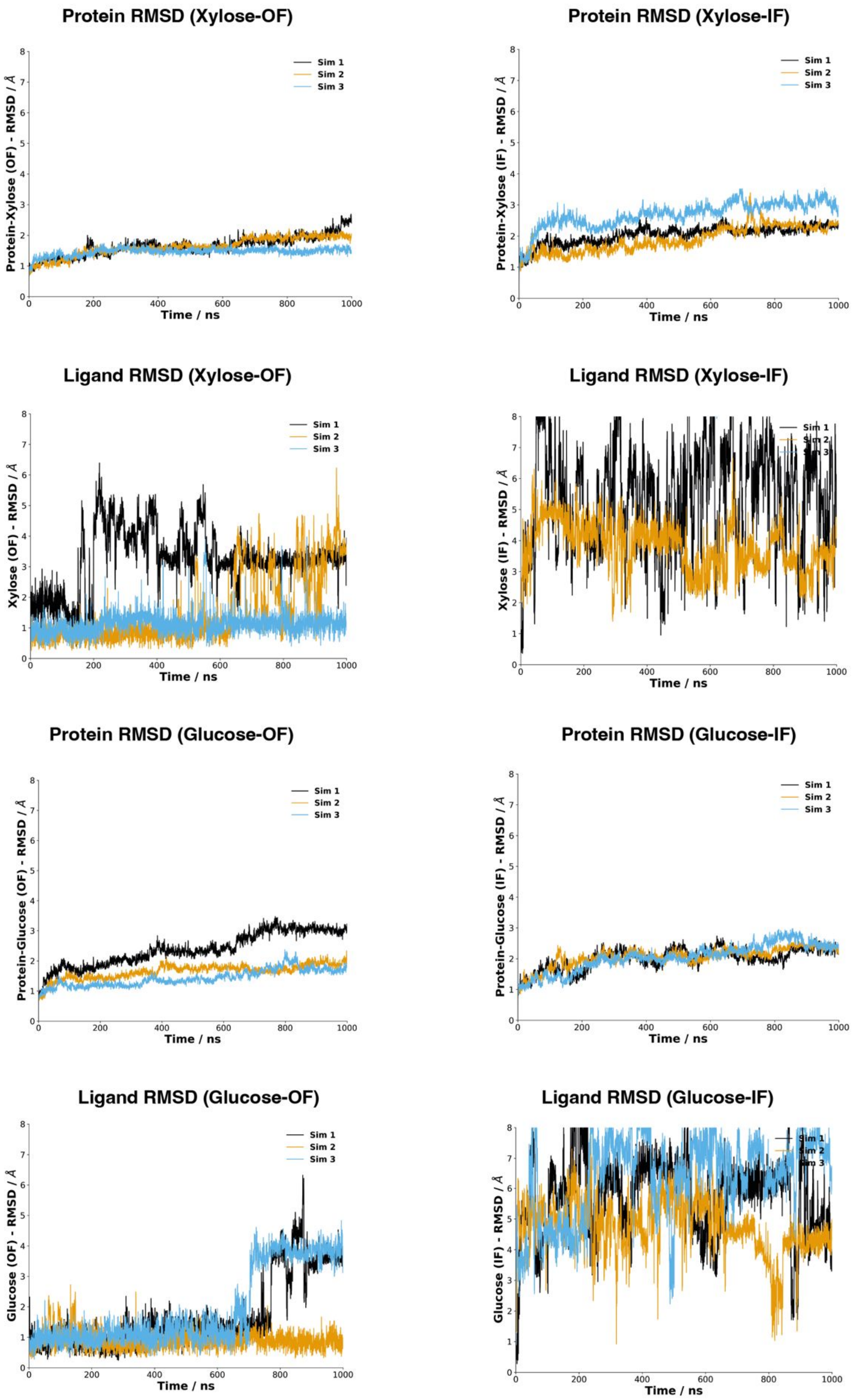
Protein RMSD (Xylose-OF)
Protein-Xylose (OF) - RMSD / Å
Protein RMSD (Xylose-IF)
Protein-Xylose (IF) - RMSD / Å
Ligand RMSD (Xylose-OF)
Xylose (OF) - RMSD / Å
Ligand RMSD (Xylose-IF)
Xylose (IF) - RMSD / Å
Protein RMSD (Glucose-OF)
Protein-Glucose (OF) - RMSD / Å
Protein RMSD (Glucose-IF)
Protein-Glucose (IF) - RMSD / Å
Ligand RMSD (Glucose-OF)
Glucose (OF) - RMSD / Å
Ligand RMSD (Glucose-IF)
Glucose (IF) - RMSD / Å
Time / ns
Sim 1
Sim 2
Sim 3

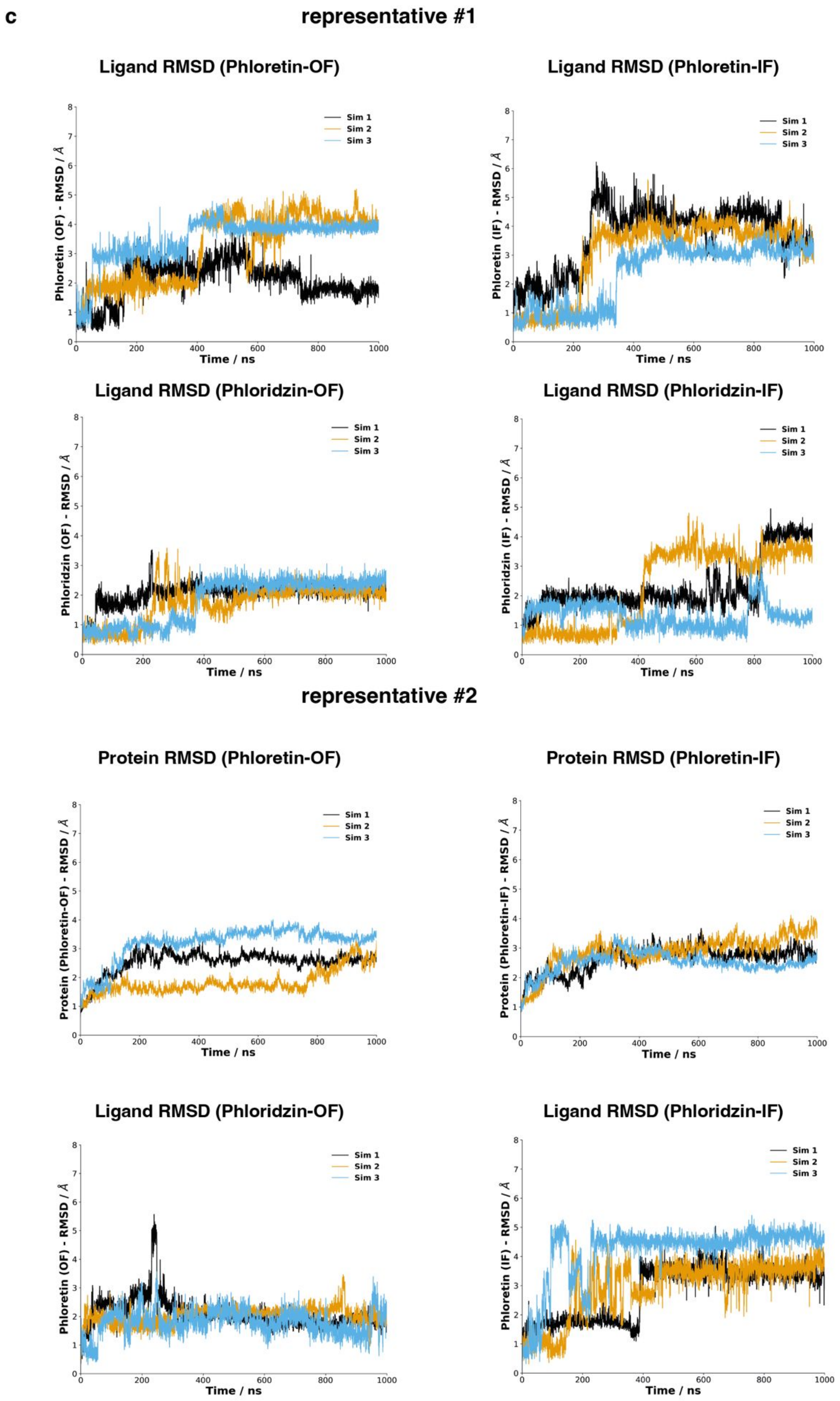
c
representative #1
Ligand RMSD (Phloretin-OF)
Ligand RMSD (Phloretin-IF)
Phloretin (OF) - RMSD / Å
Phloretin (IF) - RMSD / Å
Sim 1
Sim 2
Sim 3
Time / ns
Ligand RMSD (Phloridzin-OF)
Ligand RMSD (Phloridzin-IF)
Phloridzin (OF) - RMSD / Å
Phloridzin (IF) - RMSD / Å
representative #2
Protein RMSD (Phloretin-OF)
Protein RMSD (Phloretin-IF)
Protein (Phloretin-OF) - RMSD / Å
Protein (Phloretin-IF) - RMSD / Å
Ligand RMSD (Phloridzin-OF)
Ligand RMSD (Phloridzin-IF)
Phloretin (OF) - RMSD / Å
Phloretin (IF) - RMSD / Å

**Figure S9 RMSD plots of protein and ligand structures in OF and IF conformation over the 1 μs simulation time**. (**a**)apo state (4GBY and 4JA4). (**b**) Xylose (4GBY and 4JA4)-, glucose (4GBZ and 4JA4)-bound states. (**c**) Phloretin (4GBY and 4JA4) phloridzin (4GBY and 4JA4))-bound states. Simulations of (**a**) and (**b**) were generated using crystal structures directly or docked ligands into crystal structures, and (**c**) were generated by docking ligands into representative MD structures. Three independent simulation runs are shown in black, orange and blue, respectively.

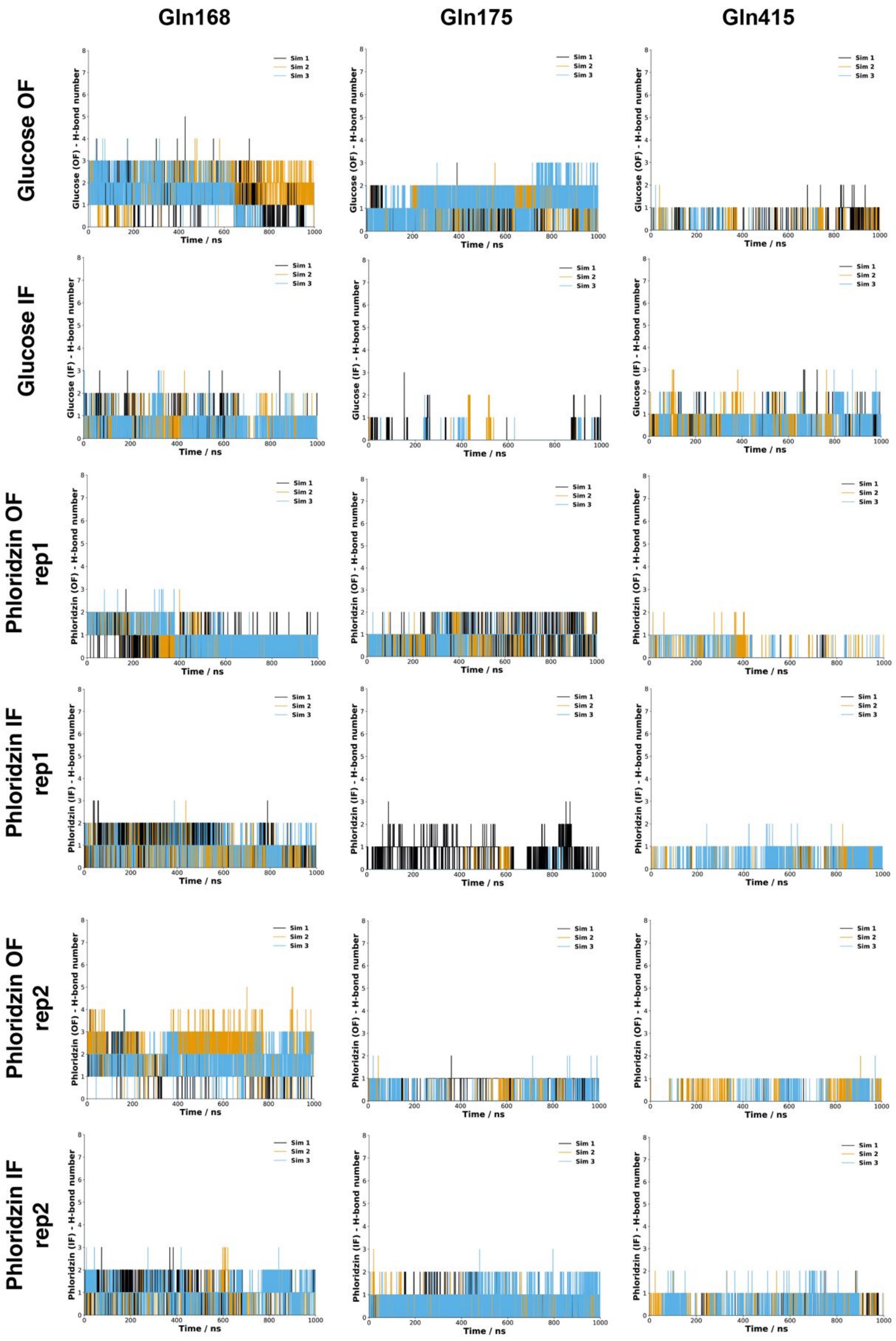
Gln168
Gln175
Gln415
Glucose OF
Glucose IF
Phloridzin OF rep1
Phloridzin IF rep1
Phloridzin OF rep2
Phloridzin IF rep2
Sim 1
Sim 2
Sim 3
Glucose (OF) - H-bond number
Glucose (IF) - H-bond number
Phloridzin (OF) - H-bond number
Phloridzin (IF) - H-bond number
Time / ns

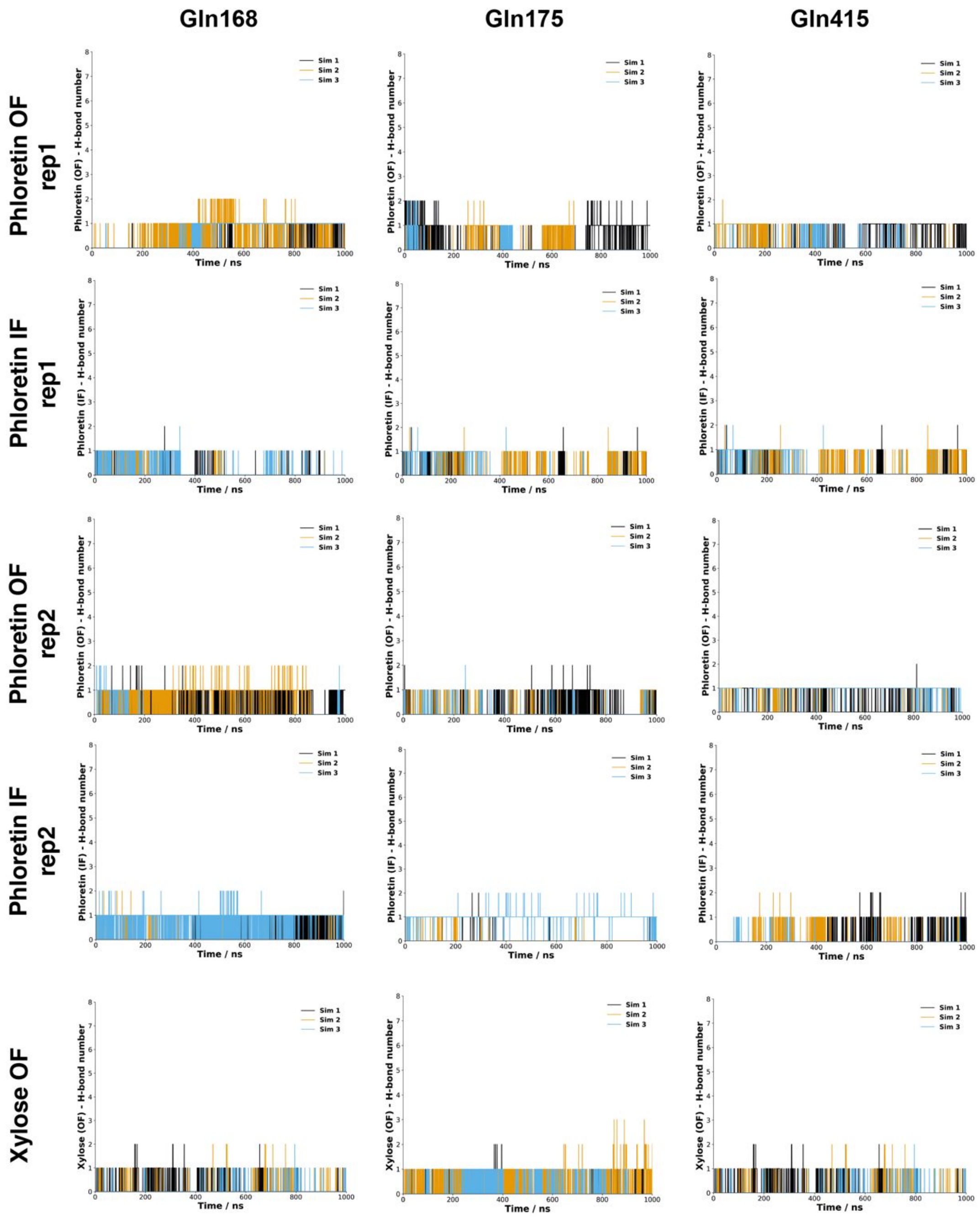

**Figure S10 Hydrogen bond interaction analysis for glucose-, phloridzin-, phloretin-, and xylose-bound structures.** The number of hydrogen bond interactions between ligand (glucose, phloridzin, phloretin and xylose) with residue Gln168, Gln175 and Gln415 over simulation time were plotted separately for three independent simulations in outward and inward-facing conformation. Xylose was dissociated from xylose-bound structures in inward-facing conformation from MD simulations, therefore, it was excluded from the H-bond analysis.

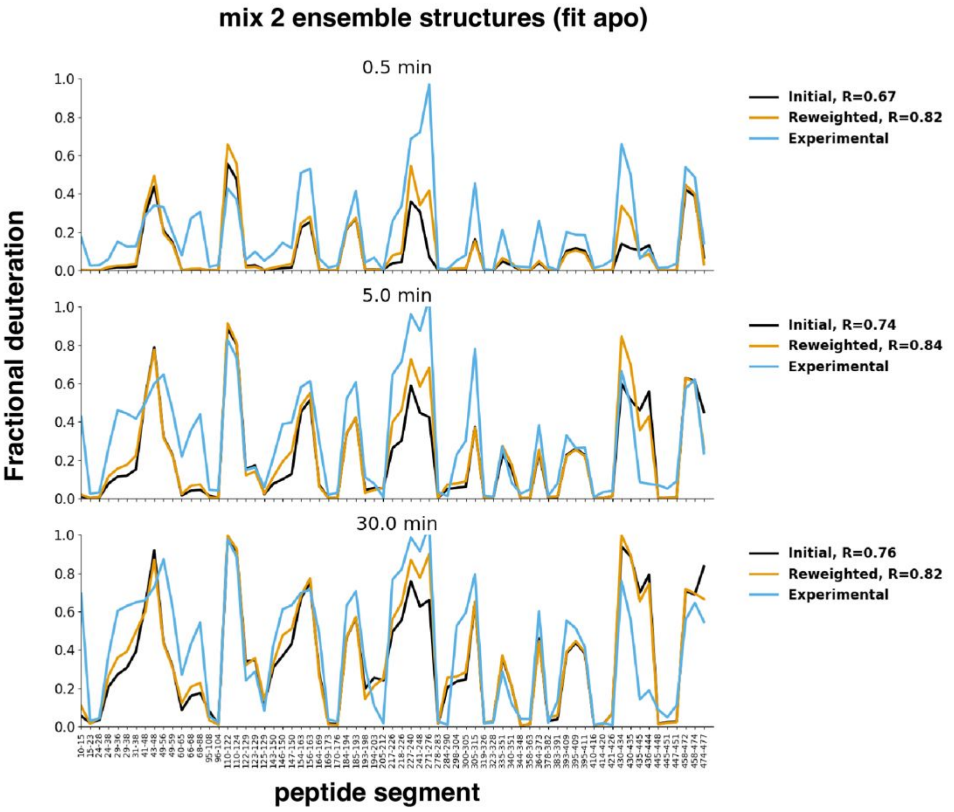
mix 2 ensemble structures (fit apo)
0.5 min
Initial, R=0.67
Reweighted, R=0.82
Experimental
5.0 min
Initial, R=0.74
Reweighted, R=0.84
Experimental
30.0 min
Initial, R=0.76
Reweighted, R=0.82
Experimental
Fractional deuteration
peptide segment

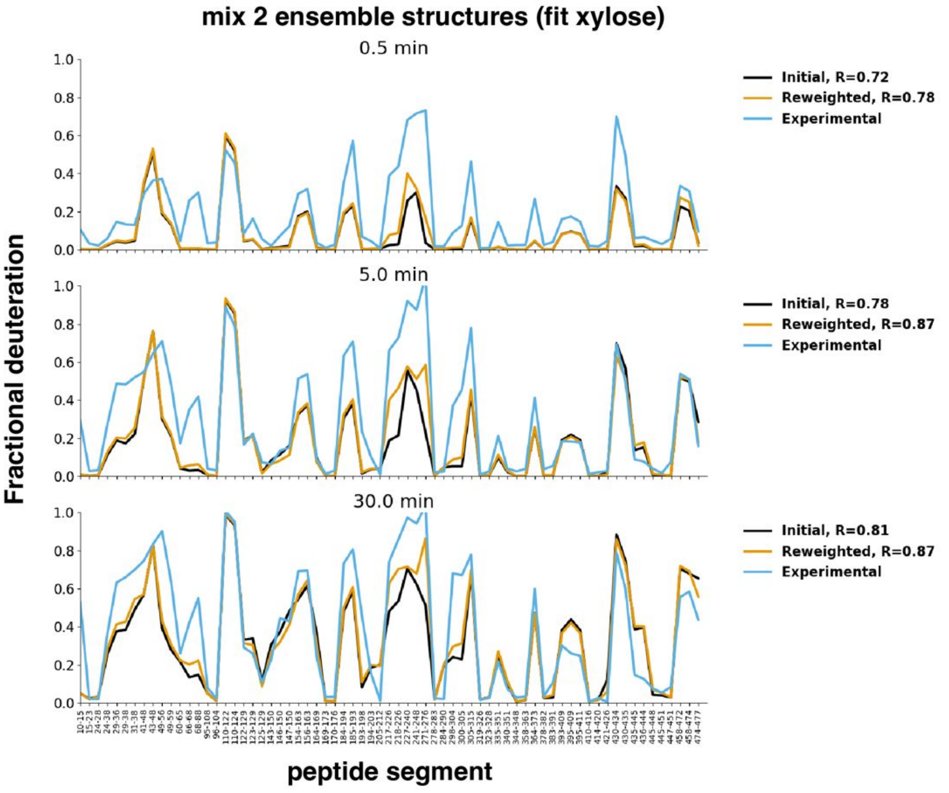
mix 2 ensemble structures (fit xylose)
0.5 min
Initial, R=0.72
Reweighted, R=0.78
Experimental
5.0 min
Initial, R=0.78
Reweighted, R=0.87
Experimental
30.0 min
Initial, R=0.81
Reweighted, R=0.87
Experimental
Fractional deuteration
peptide segment

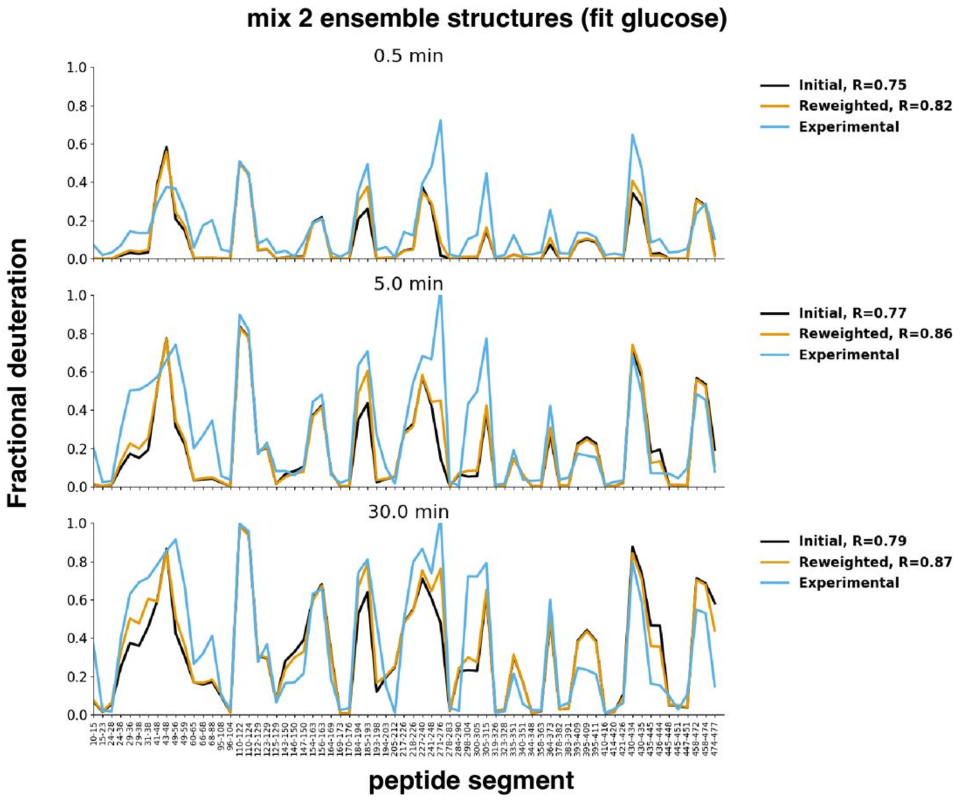
mix 2 ensemble structures (fit glucose)
0.5 min
5.0 min
30.0 min
Initial, R=0.75
Reweighted, R=0.82
Experimental
Initial, R=0.77
Reweighted, R=0.86
Experimental
Initial, R=0.79
Reweighted, R=0.87
Experimental
Fractional deuteration
peptide segment

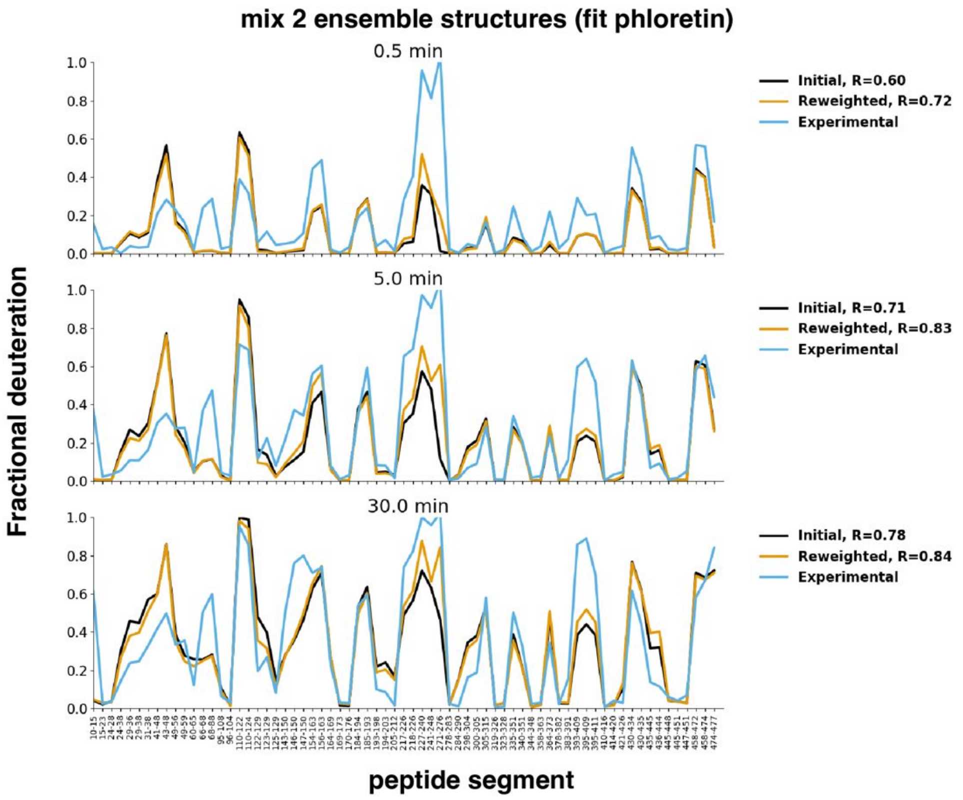
mix 2 ensemble structures (fit phloretin)
0.5 min
5.0 min
30.0 min
Initial, R=0.60
Reweighted, R=0.72
Experimental
Initial, R=0.71
Reweighted, R=0.83
Experimental
Initial, R=0.78
Reweighted, R=0.84
Experimental
Fractional deuteration
peptide segment

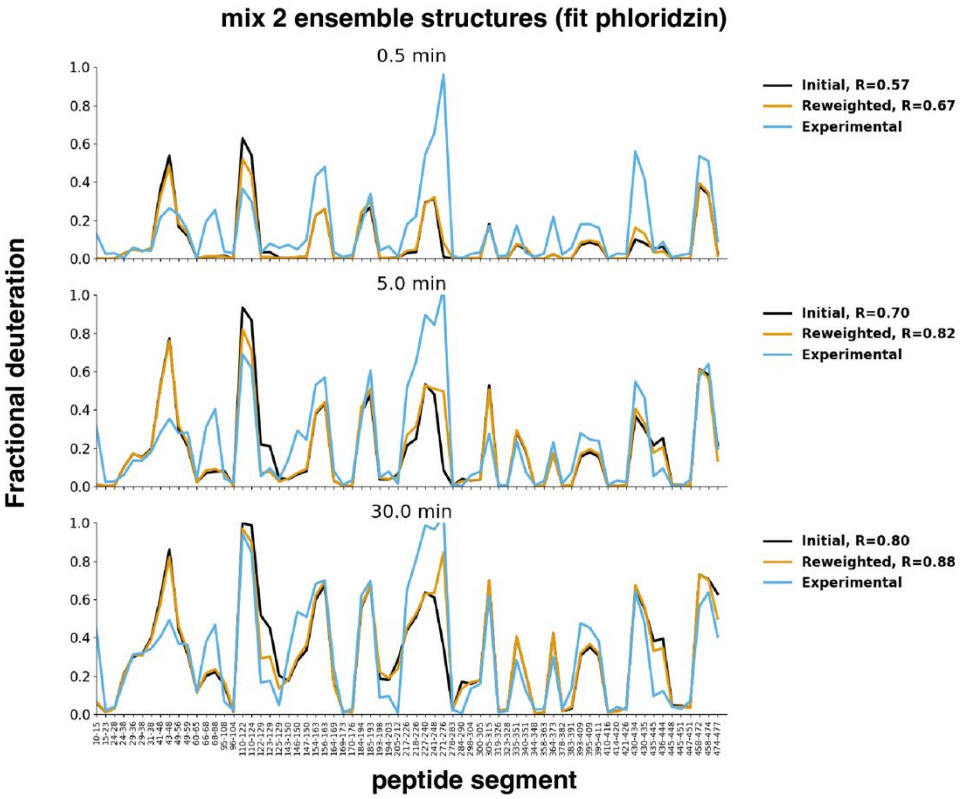

**Figure S11 Fractional deuteration before and after reweighting.** Fractional deuteration for initial predicted, reweighted and experimental HDX data were shown in black, orange and blue, respectively.

a

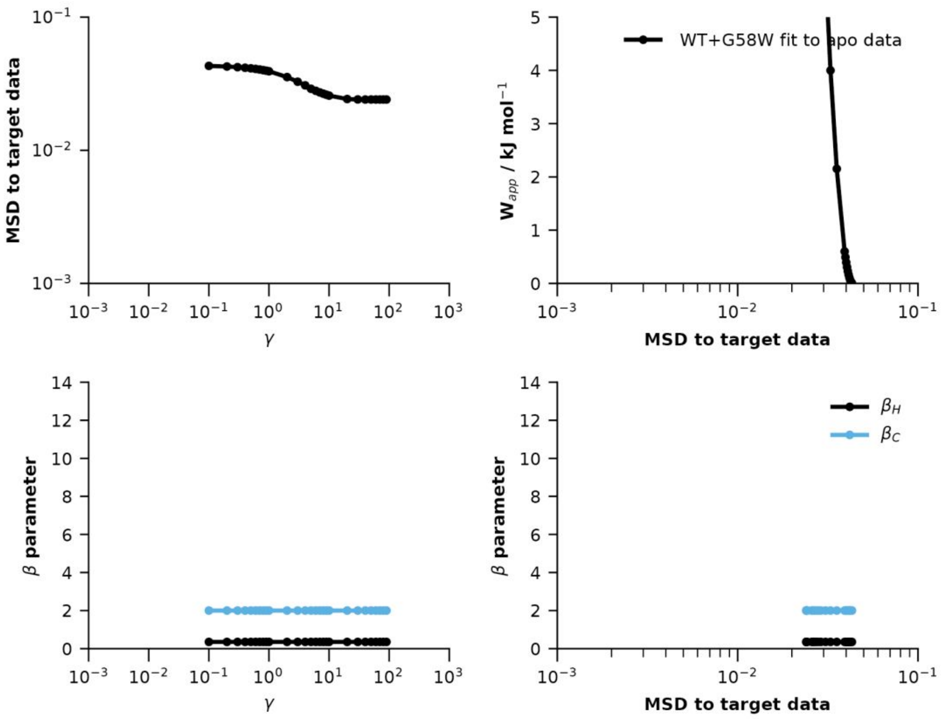
MSD to target data
γ
WT+G58W fit to apo data
W$_{app}$ / kJ mol$^{-1}$
MSD to target data
β parameter
γ
β parameter
$\beta_H$
$\beta_C$
MSD to target data

b

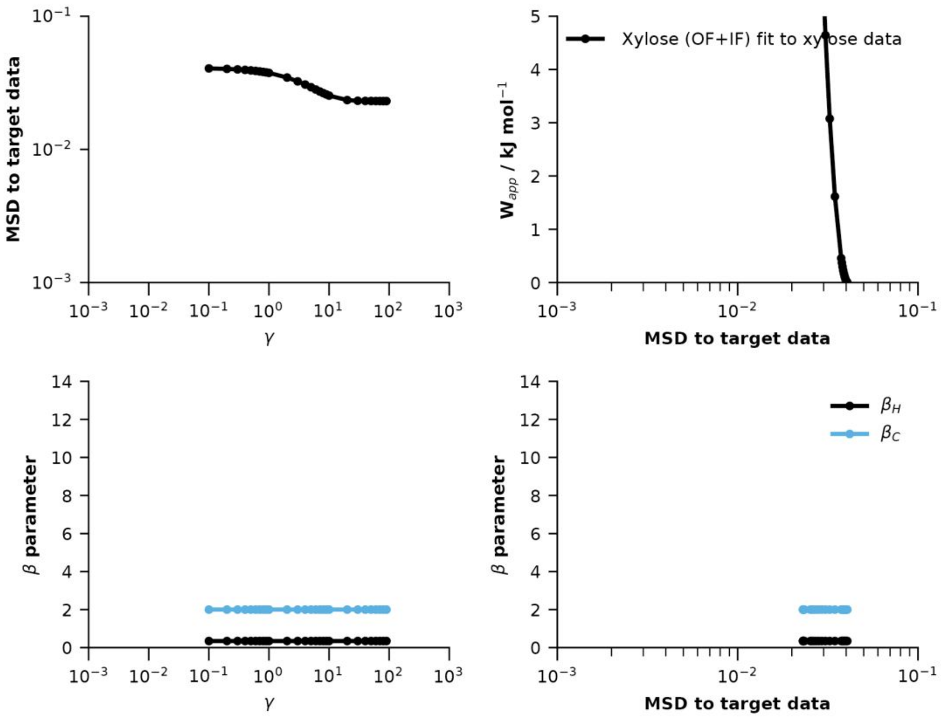
MSD to target data
γ
Xylose (OF+IF) fit to xylose data
W$_{app}$ / kJ mol$^{-1}$
MSD to target data
β parameter
γ
β parameter
$\beta_H$
$\beta_C$
MSD to target data

**c**

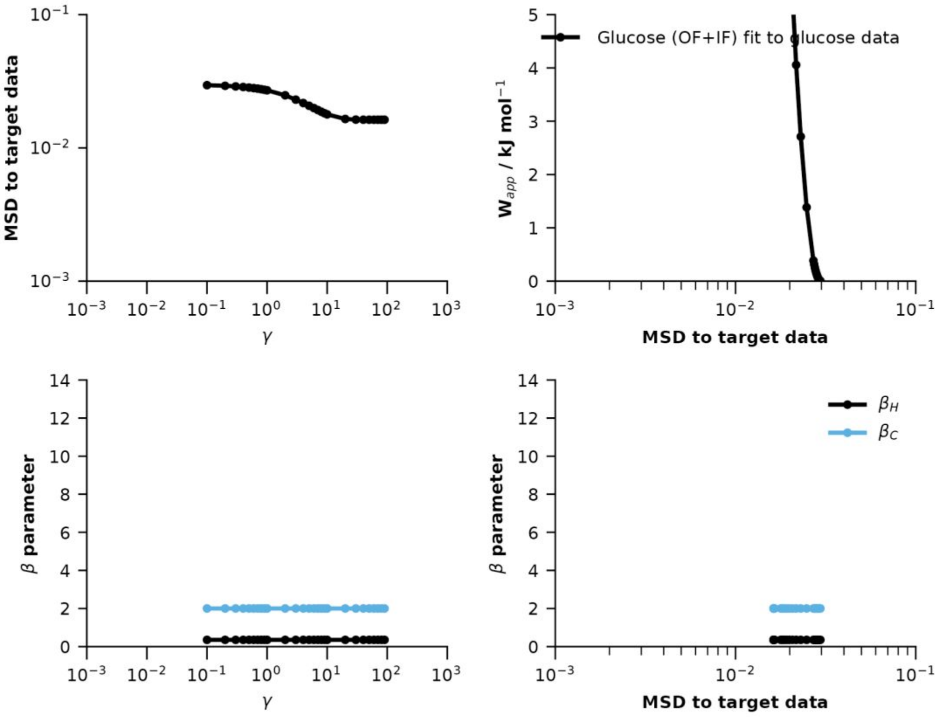

MSD to target data
γ
Glucose (OF+IF) fit to glucose data
W_app / kJ mol^-1
β parameter
β_H
β_C

**d**

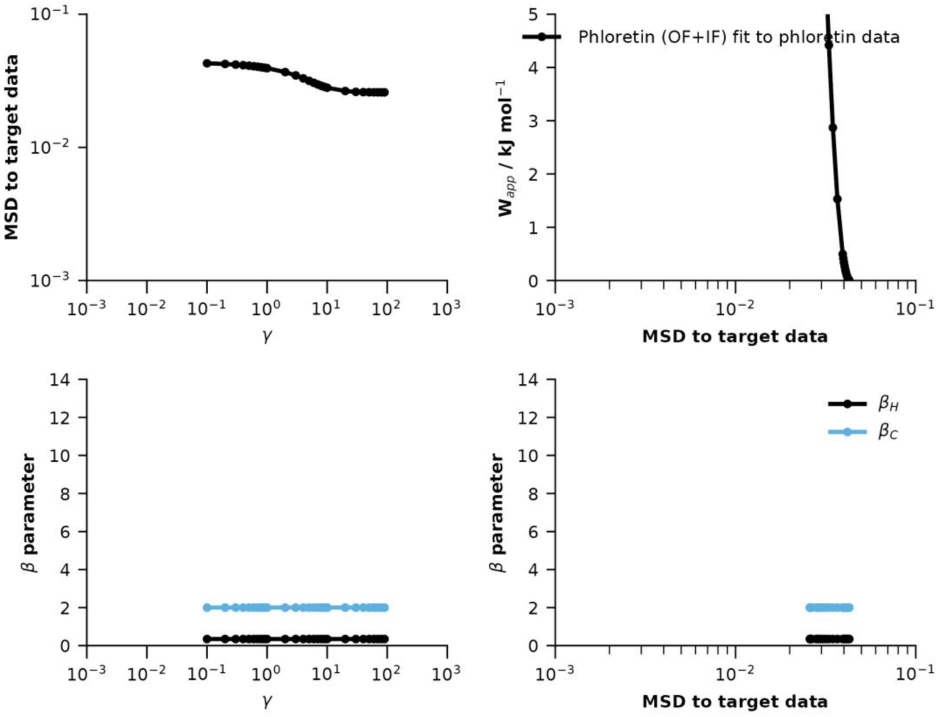

MSD to target data
γ
Phloretin (OF+IF) fit to phloretin data
W_app / kJ mol^-1
β parameter
β_H
β_C

**e**

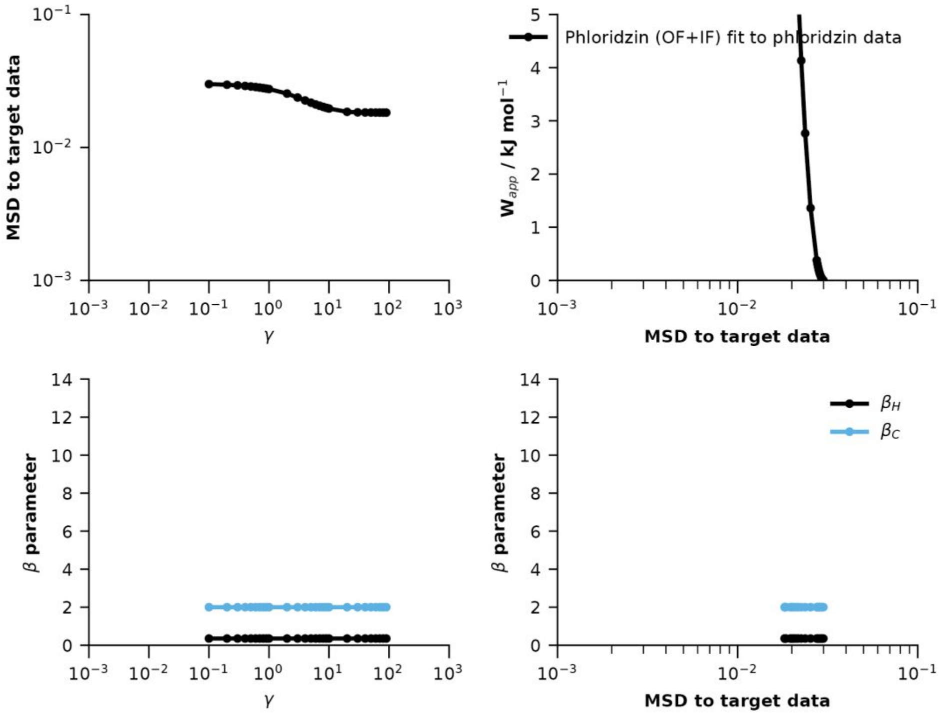

**Figure S12 Decision plots on ensemble reweighting.** Ensemble reweighting was carried out by fitting ensemble structures (**a**) WT and G58W, (**b**) Xylose-bound (OF and IF), (**c**) Glucose-bound (OF and IF), (**d**) Phloretin-bound (OF and IF) and (**e**)Phloridzin-bound (OF and IF) to experimental XylE (**a**) apo, (**b**) Xylose-, (**c**) Glucose-, (**d**) Phloretin- and (**e**) Phloridzin-bound data, respectively. Top left: Relationship between mean-square deviation (MSD) to target data and $\gamma$. Top right: Relationship between applied work (kJ/mol) and MSD to target data. Bottom left: Relationship between scaling factor ($\beta_H$ and$\beta_C$) and $\gamma$. Bottom right: Relationship between scaling factor ($\beta_H$ and$\beta_C$) and MSD to target data.

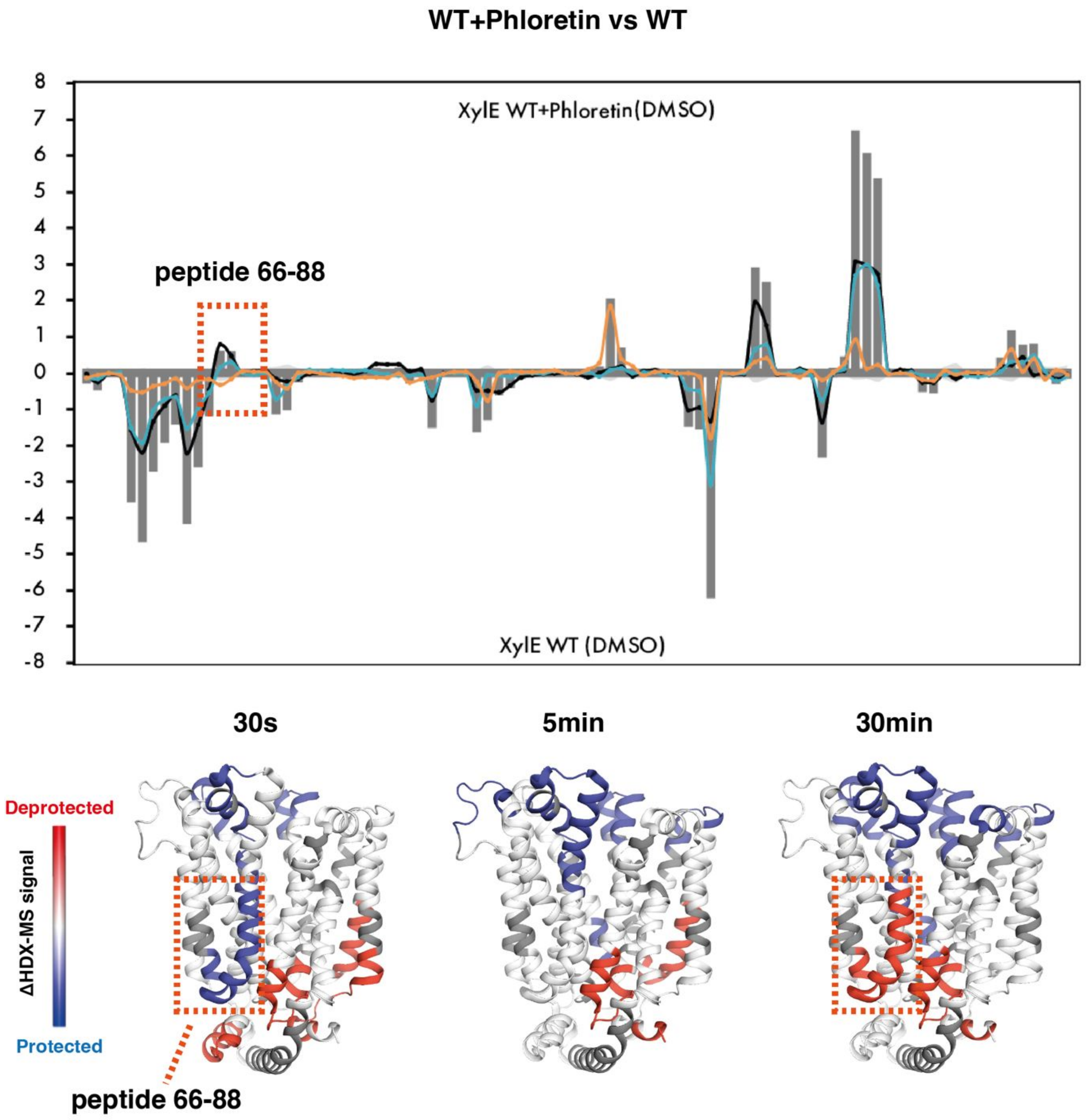

**Figure S13 Butterfly plot of comparison between phloretin-bound and WT structures**. Peptide 66-88 presents different HDX patterns at the 30s and 30min time points.

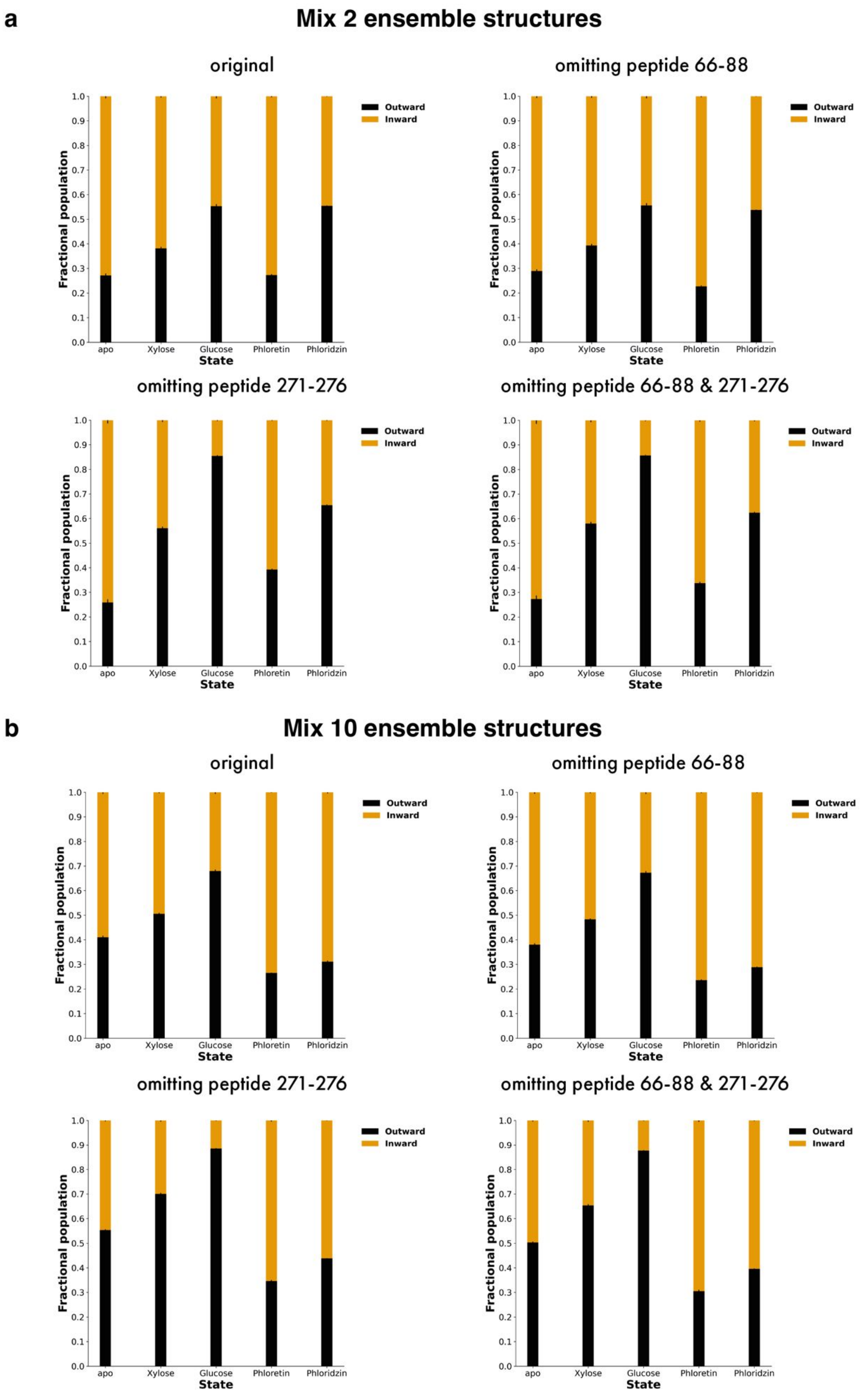
a
Mix 2 ensemble structures
original
omitting peptide 66-88
omitting peptide 271-276
omitting peptide 66-88 & 271-276
b
Mix 10 ensemble structures
original
omitting peptide 66-88
omitting peptide 271-276
omitting peptide 66-88 & 271-276
Fractional population
State
apo
Xylose
Glucose
Phloretin
Phloridzin
Outward
Inward
0.0
0.1
0.2
0.3
0.4
0.5
0.6
0.7
0.8
0.9
1.0

**c** **Mix 14 ensemble structures**

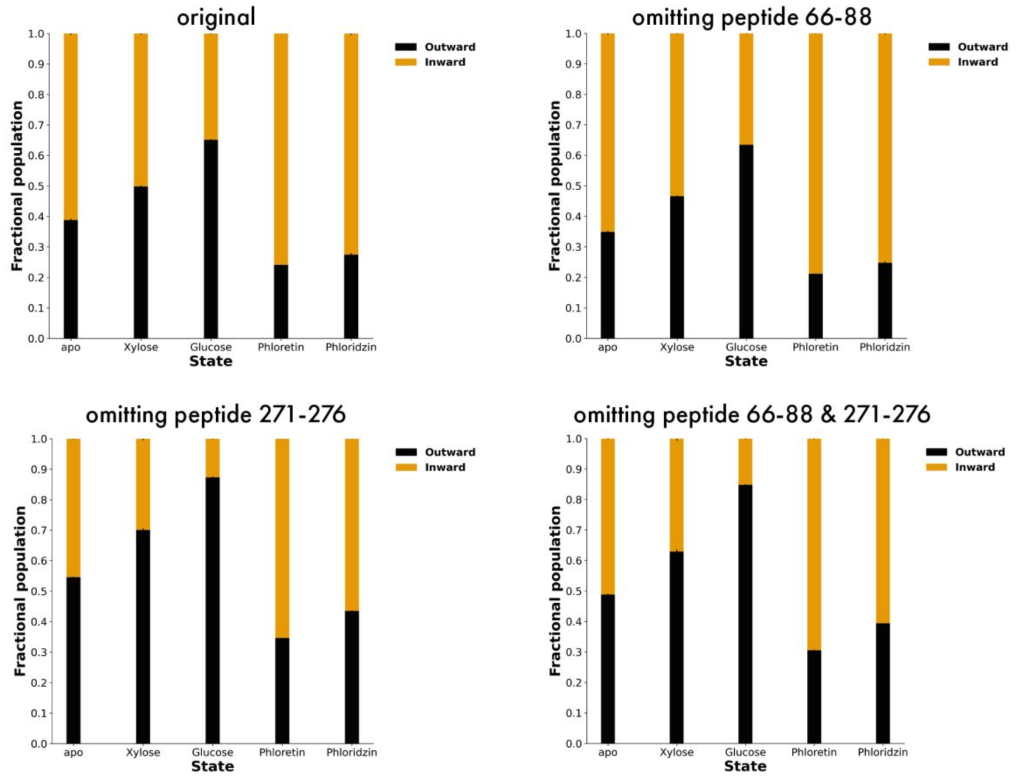

**d** **Compare representative ensemble structures (Phloretin/Phloridzin)**

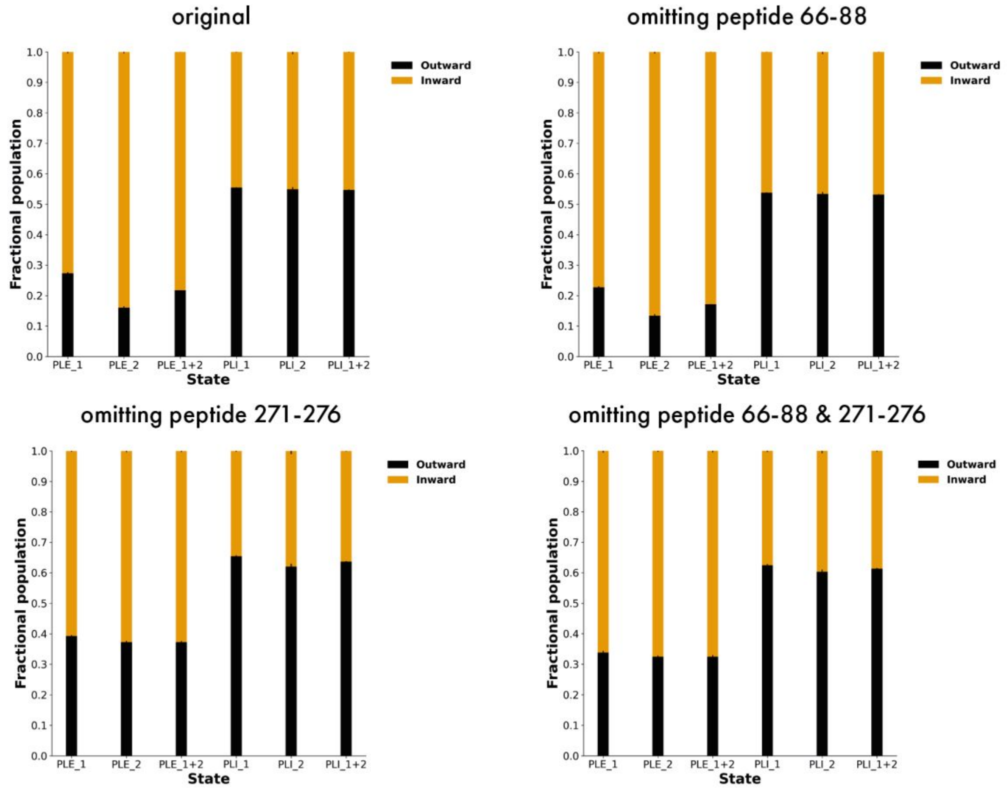

**Figure S14 Bar charts of fractional population after reweighting under 4 conditions (original, omitting peptide 66-88, peptides 271-276 or both).** (**a**) Mix 2 ensemble structures. (**b**) Mix 10 ensemble structures. (**c**) Mix 14 ensemble structures. (**d**) Compare fractional population using ensemble structures for phloretin and phloridzin (representative structures 1, 2 and both).

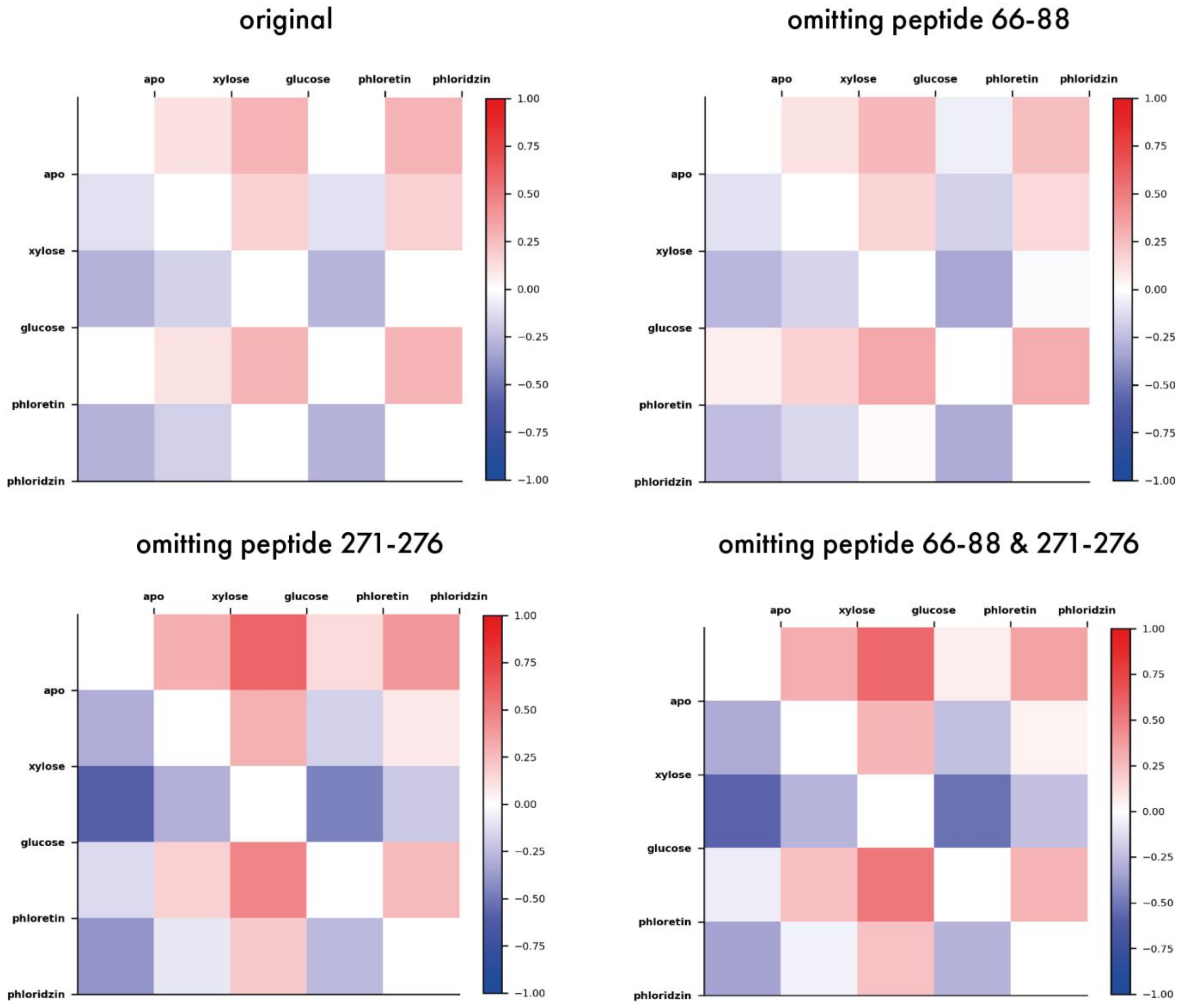

**Figure S15 Heap map of relative fractional population of mixing "state-specific" states between WT and ligand-bound states under four conditions (original, omitting peptide 66-88, peptides 271-276 or both) for mixing 2 ensemble structures.** Red indicates relatively outward-facing, and blue indicates relatively inward-facing.

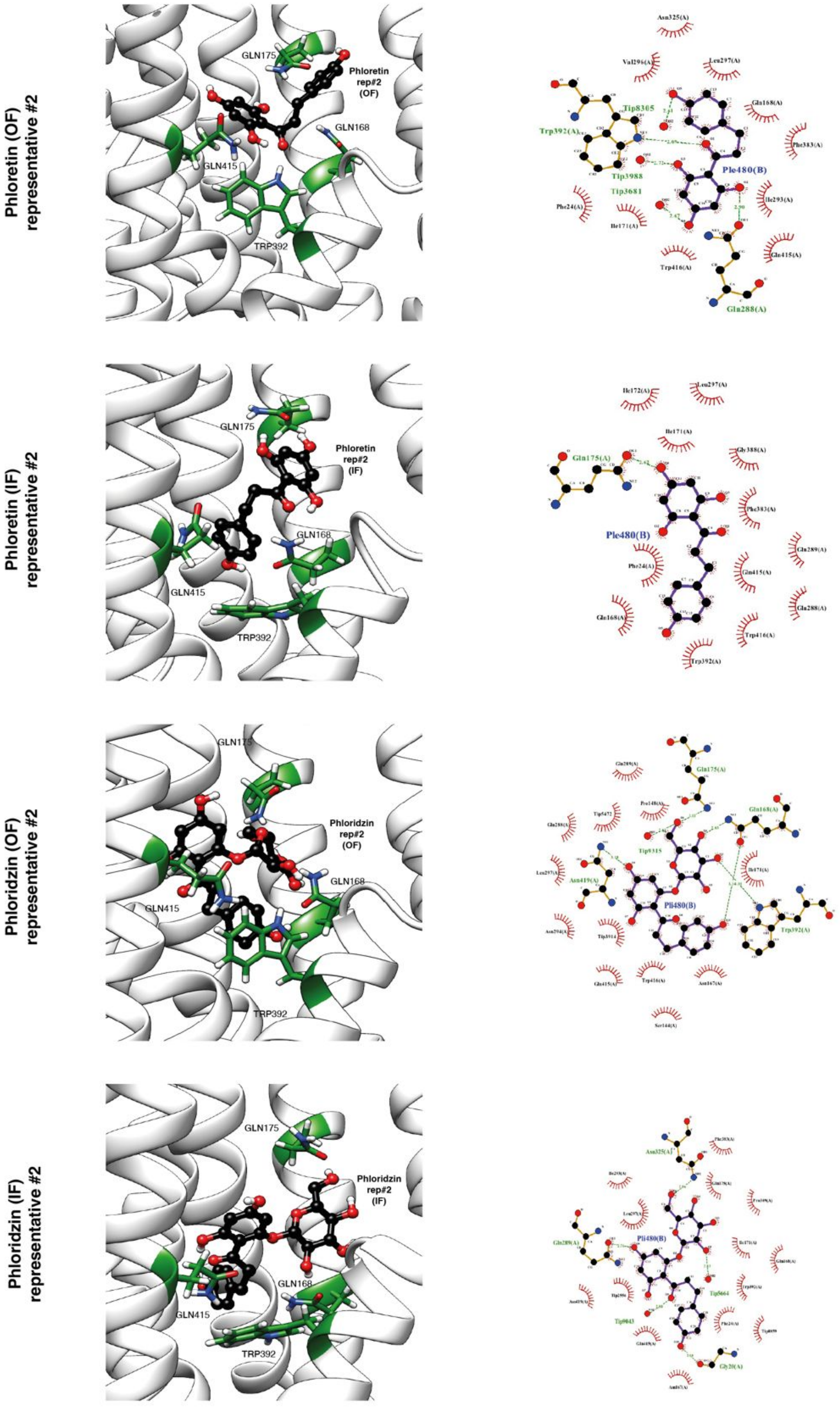

**Figure S16 Coordination of second representative phloretin and phloridzin by XylE in outward-facing and inward-facing structures and 2D protein-ligand interaction diagram.** Phloretin and phloridzin were shown in black balls and sticks. The binding site residues in XylE were coloured green. The protein-ligand interaction plots were generated by LigPlot+[2]. Hydrogen bonds were shown as green dotted lines, hydrophobic interactions were represented by red eyelashes.

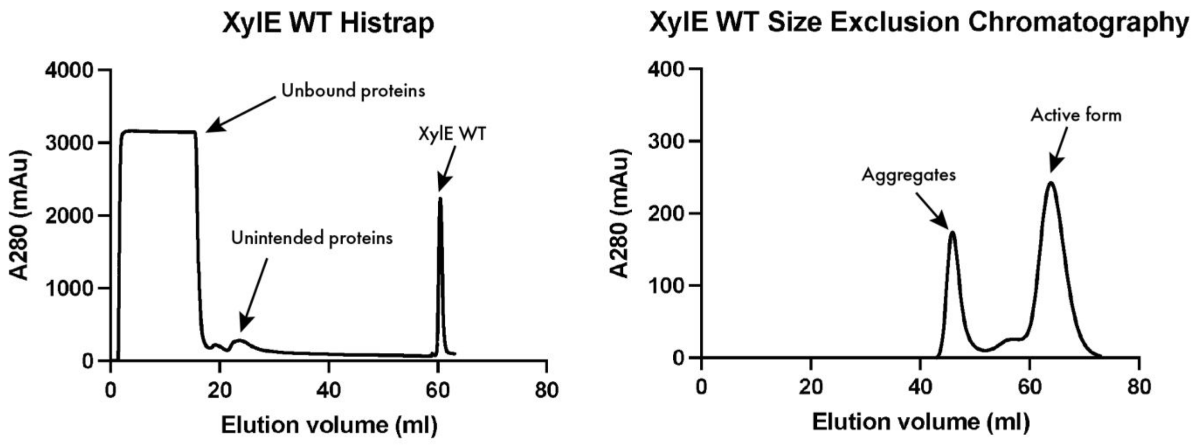

**Figure S17 XylE WT chromatogram of Histrap and Size Exclusion Chromatography.**

a

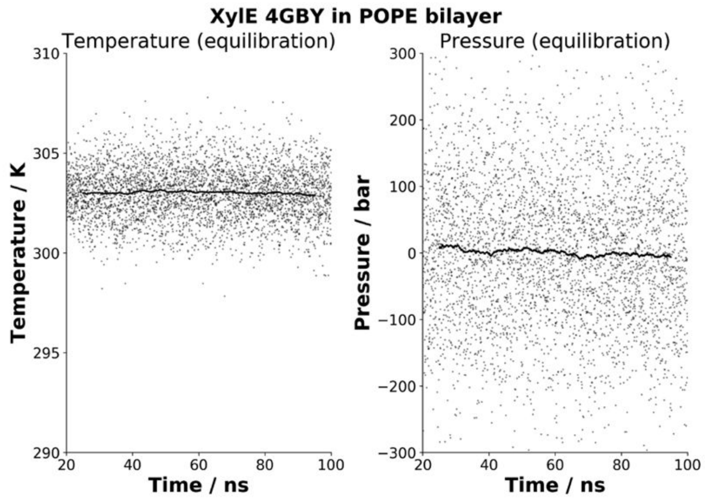

b

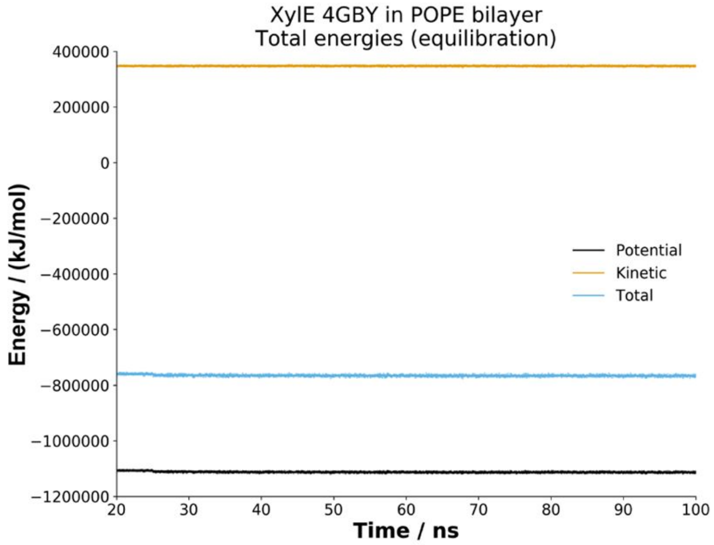

c

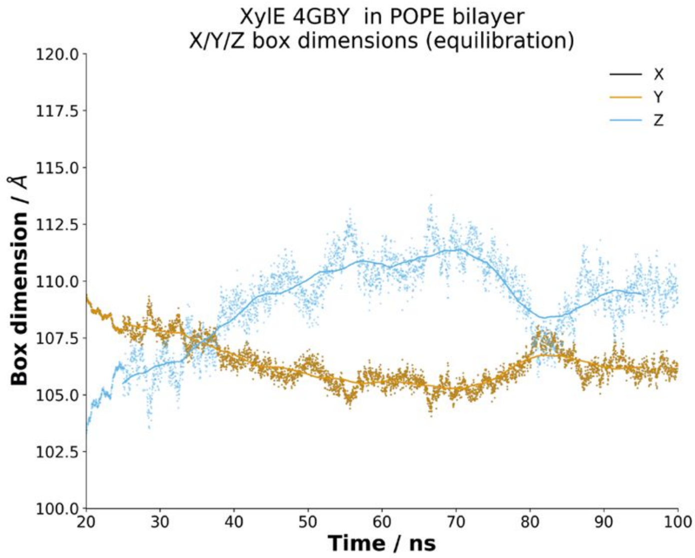

**Figure S18 Trajectory analysis of MD simulations at the equilibrium stage.** (**a**) Temperature and pressure fluctuate over time. (**b**)Potential, kinetic and total energy over simulation time. (**c**) Box dimension.

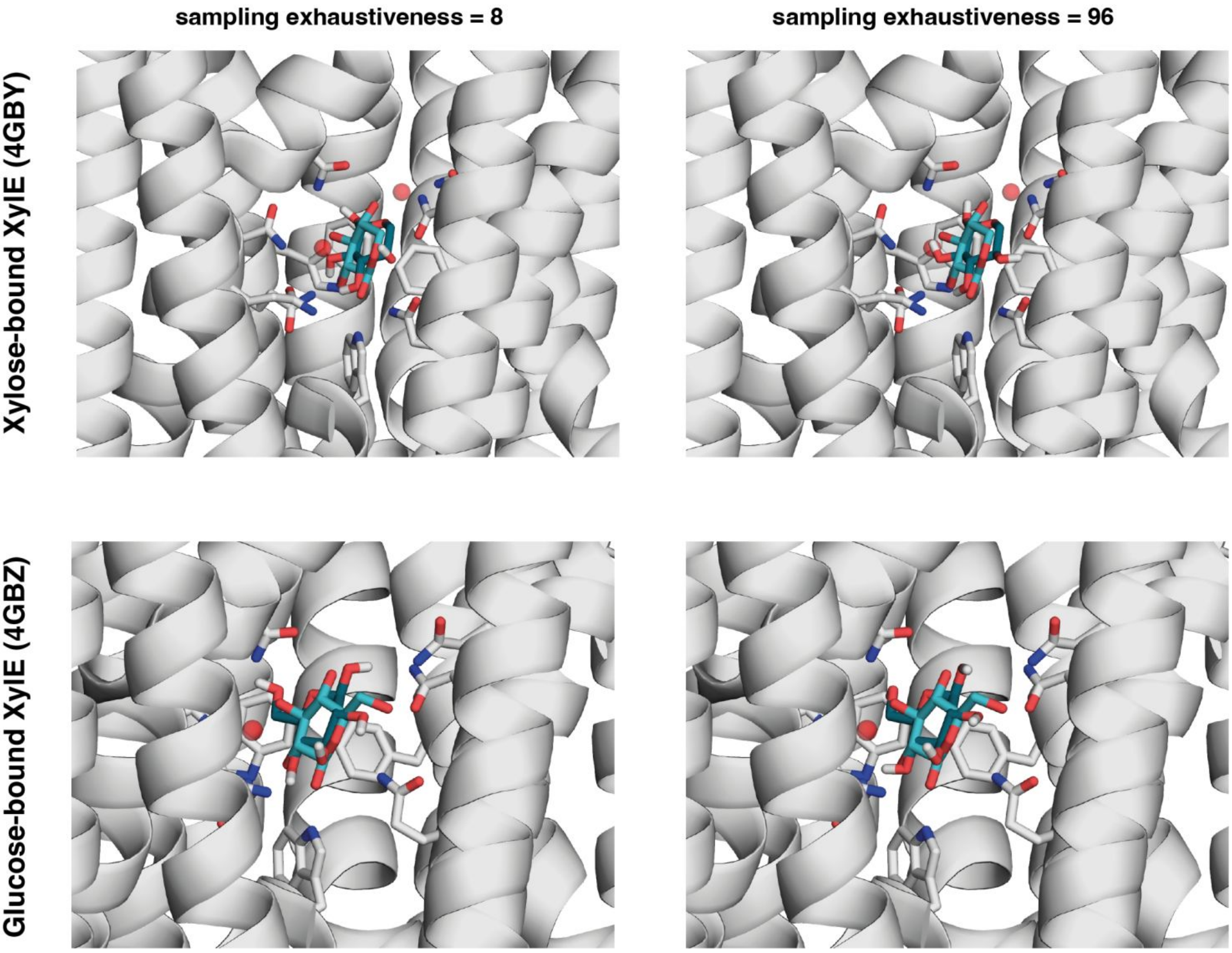

**Figure S19 Evaluation of docking xylose/glucose to crystal structure using Autodock Vina.** Two levels of exhaustiveness (8 and 96) were tested for docking xylose and glucose to their associated crystal structure (4GBY and 4GBZ). Crystal ligand structures were shown in cyan; the top pose from docking was shown in dark green. Binding site residues (F24, Q168, Q175, Q288, Q289, N294, N325, W392, Q415 and W416) were shown in white.

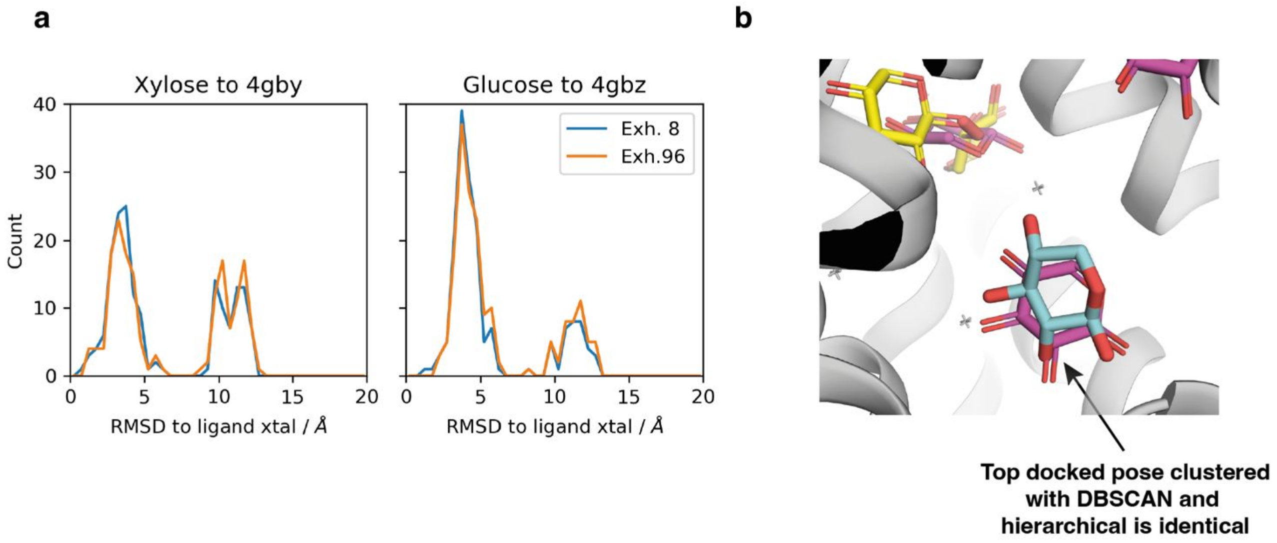

**Figure S20 Docking analysis of xylose/glucose structures.** (**a**) Distribution plots for rigid docking poses (xylose docked to XylE OF (4GBY) and glucose docked to XylE OF (4GBZ)). Each docking generated 9 poses, which led to 171 docked poses for 19 representative protein structures from MD simulations. Rigid docking was tested in two levels of exhaustiveness (8 and 96). (**b**) Representative poses from docking were clustered using DBSCAN and hierarchical clustering methods for rigid docking. Xylose pose from crystal structure was shown in cyan, docked xylose (clustered using DBSCAN) was shown in yellow, and docked xylose (clustered using hierarchical clustering method) was shown in magenta. the top docked pose in the binding was identical from DBSCAN and hierarchical clustering, with an RMSD value of 1.626 Å to ligand pose in the crystal structure.

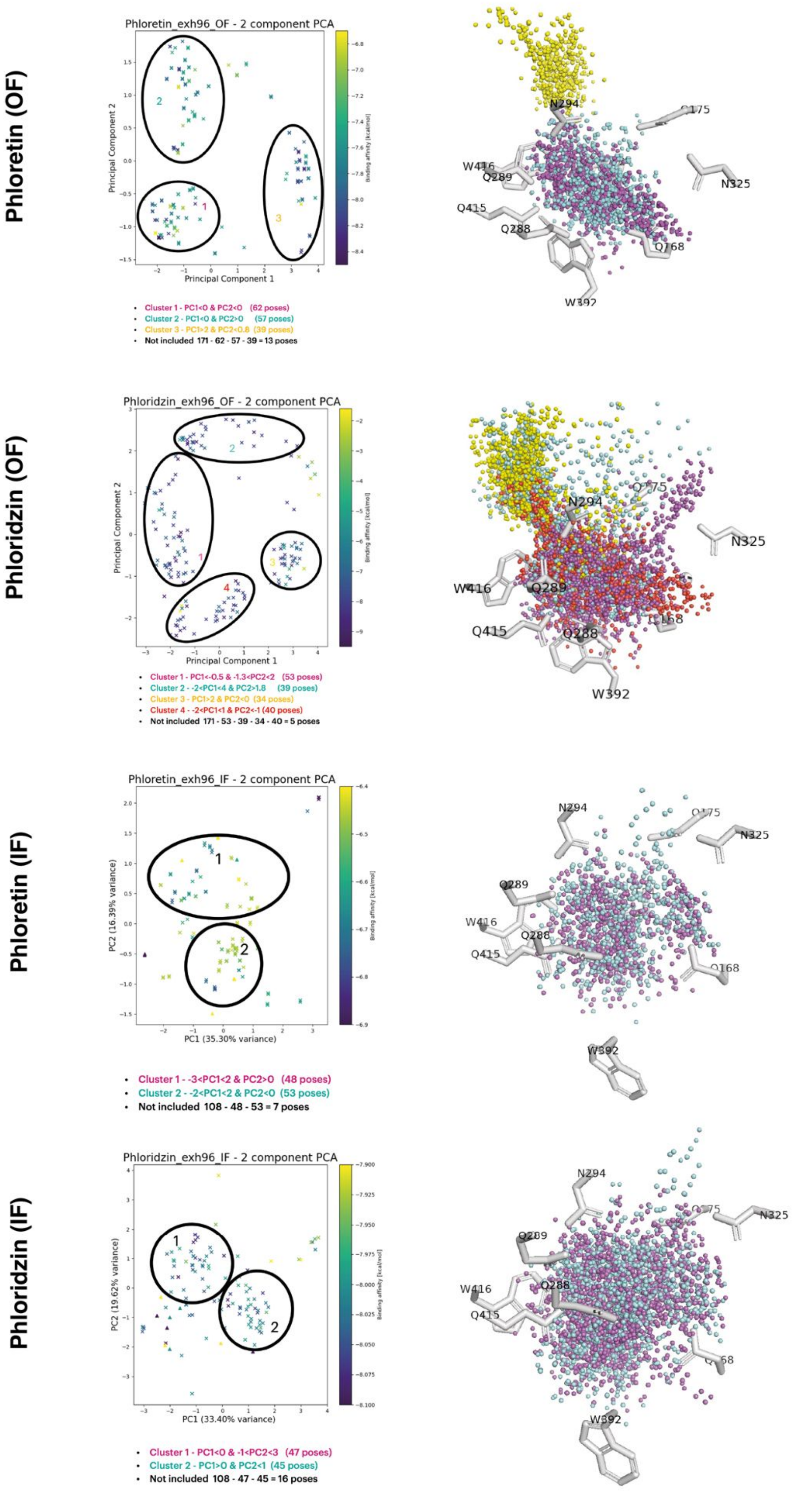

Phloretin (OF)
Phloretin_exh96_OF - 2 component PCA
Principal Component 1
Principal Component 2
Binding affinity [kcal/mol]
Cluster 1 - PC1<0 & PC2<0 (62 poses)
Cluster 2 - PC1<0 & PC2>0 (57 poses)
Cluster 3 - PC1>2 & PC2<0.8 (39 poses)
Not included 171 - 62 - 57 - 39 = 13 poses
N294
W416
Q289
Q415
Q288
N325
Q168
W392
Phloridzin (OF)
Phloridzin_exh96_OF - 2 component PCA
Principal Component 1
Principal Component 2
Binding affinity [kcal/mol]
Cluster 1 - PC1<-0.5 & -1.3<PC2<2 (53 poses)
Cluster 2 - -2<PC1<4 & PC2>1.8 (39 poses)
Cluster 3 - PC1>2 & PC2<0 (34 poses)
Cluster 4 - -2<PC1<1 & PC2<-1 (40 poses)
Not included 171 - 53 - 39 - 34 - 40 = 5 poses
N294
N325
W416
Q289
Q415
Q288
W392
Phloretin (IF)
Phloretin_exh96_IF - 2 component PCA
PC1 (35.30% variance)
PC2 (16.39% variance)
Binding affinity [kcal/mol]
Cluster 1 - -3<PC1<2 & PC2>0 (48 poses)
Cluster 2 - -2<PC1<2 & PC2<0 (53 poses)
Not included 108 - 48 - 53 = 7 poses
N294
N325
Q289
W416
Q288
Q415
Q168
Phloridzin (IF)
Phloridzin_exh96_IF - 2 component PCA
PC1 (33.40% variance)
PC2 (19.62% variance)
Binding affinity [kcal/mol]
Cluster 1 - PC1<0 & -1<PC2<3 (47 poses)
Cluster 2 - PC1>0 & PC2<1 (45 poses)
Not included 108 - 47 - 45 = 16 poses
N294
N325
Q289
W416
Q288
Q415

**Figure S21 PCA analysis of docked poses for phloretin and phloridzin.** Two-component PCA analysis was performed for docked poses of phloretin and phloridzin structures. PCA plots were generated using a custom python script. Clusters were coloured in magenta, cyan, yellow, red and black for clusters 1, 2, 3, 4 and "not included", respectively. Binding site residues were shown in white stick.

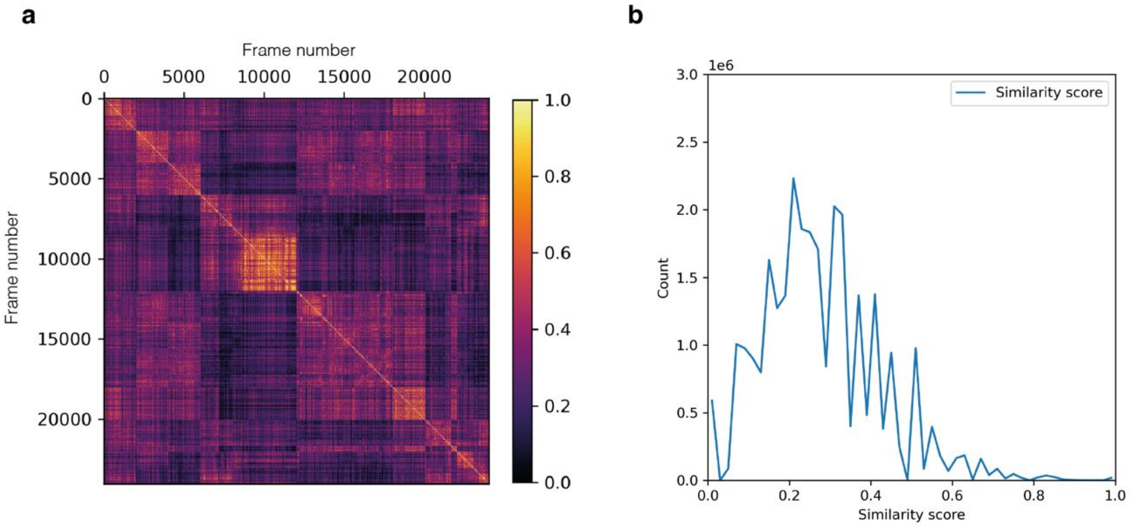

**Figure S22 2D similarity score for protein-ligand interaction fingerprint analysis.** (**a**) Pairwise Tanimoto coefficient of XylE-phloretin fingerprint (24012 MD frames) in OF conformation was calculated and plotted using a custom python script. Frames were coloured in gradient, yellow indicates a high similarity score, and black indicates a low similarity score. (**b**) Histogram of similarity score distribution for XylE-phloretin fingerprint (24012 MD frames) in OF conformation. The similarity score was displayed on the x-axis and the number of frames count ($1e^6$) was displayed on the y-axis.

### Supporting Tables

**Table S1 Binding occupancy of XylE WT binding to xylose, glucose, phloretin and phloridzin.**

| | Kd[3] (μM) | Protein (μM) | | Ligand (μM) | | % Protein bound | |
|---|---|---|---|---|---|---|---|
| | | Equilibration solution | Labelling solution | Equilibration solution | Labelling solution | Equilibration solution | Labelling solution |
| **Xylose** | 350 | 13.86 | 1.37 | 30000.00 | 3000.0 | 98.85% | 89.55% |
| **Glucose** | 770 | 13.86 | 1.37 | 60000.00 | 6000.00 | 98.73% | 88.62% |
| **Phloretin** | 33.9 | 13.86 | 1.37 | 3000.00 | 300.0 | 98.88% | 89.81% |
| **Phloridzin** | 258 | 13.86 | 1.37 | 20000.00 | 2000.0 | 98.73% | 88.57% |

**Table S2 HDX-MS measurements.**

| **1. ΔHDX = (XylE WT+ Xylose) vs (XylE WT) in 10%DMSO** | | |
|---|---|---|
| | XylE WT + Xylose | XylE WT |
| **HDX reaction details** | 10mM potassium phosphate in $H_2O$ pH 7.0, 0.02%DDM | |
| **HDX time course (min)** | 0.5, 5, and 30 minutes | |
| **Number of peptides** | 86 | 86 |
| **Sequence Coverage** | 82.7% | 82.7% |
| **Average peptide length / Redundancy** | 8.9 / 1.6 | 8.9 / 1.6 |
| **Replicates (biological or technical)** | 3 (technical) | 3 (technical) |
| **Repeatability (average SD)** | 0.046 | 0.043 |
| **Significant differences in sum ΔHDX** | CI 99% ± 0.27 Da | |

| 2. ΔHDX = (XylE WT+ Glucose) vs (XylE WT) in 10%DMSO | | |
|---|---|---|
| | XylE WT + Glucose | XylE WT |
| **HDX reaction details** | 10mM potassium phosphate in $H_2O$ pH 7.0, 0.02%DDM | |
| **HDX time course (min)** | 0.5, 5, and 30 minutes | |
| **Number of peptides** | 86 | 86 |
| **Sequence Coverage** | 82.7% | 82.7% |
| **Average peptide length / Redundancy** | 8.9 / 1.6 | 8.9 / 1.6 |
| **Replicates (biological or technical)** | 3 (technical) | 3 (technical) |
| **Repeatability (average SD)** | 0.044 | 0.043 |
| **Significant differences in sum ΔHDX** | CI 99% ± 0.26 Da | |

| 3. ΔHDX = (XylE WT+ Phloretin) vs (XylE WT) in 10%DMSO | | |
|---|---|---|
| | XylE WT + Phloretin | XylE WT |
| **HDX reaction details** | 10mM potassium phosphate in $H_2O$ pH 7.0, 0.02%DDM | |
| **HDX time course (min)** | 0.5, 5, and 30 minutes | |
| **Number of peptides** | 86 | 86 |
| **Sequence Coverage** | 82.7% | 82.7% |
| **Average peptide length / Redundancy** | 8.9 / 1.6 | 8.9 / 1.6 |
| **Replicates (biological or technical)** | 3 (technical) | 3 (technical) |
| **Repeatability (average SD)** | 0.044 | 0.043 |
| **Significant differences in sum ΔHDX** | CI 99% ± 0.26 Da | |

| 4. ΔHDX = (XylE WT+ Phloridzin) vs (XylE WT) in 10%DMSO | | |
|---|---|---|
| | XylE WT + Phloridzin | XylE WT |
| **HDX reaction details** | 10mM potassium phosphate in $H_2O$ pH 7.0, 0.02%DDM | |
| **HDX time course (min)** | 0.5, 5, and 30 minutes | |
| **Number of peptides** | 86 | 86 |
| **Sequence Coverage** | 82.7% | 82.7% |
| **Average peptide length / Redundancy** | 8.9 / 1.6 | 8.9 / 1.6 |
| **Replicates (biological or technical)** | 3 (technical) | 3 (technical) |
| **Repeatability (average SD)** | 0.044 | 0.043 |
| **Significant differences in sum ΔHDX** | CI 99% ± 0.26 Da | |

**Table S3 HDX-MS reweighting data of mixing "state-specific" ensemble structures.**

| Mix IF+OF (apo) | $\gamma$ | Weighted RMSE to target | Applied work (kJ/mol) | Conformational population (IF:OF) |
|---|---|---|---|---|
| **Original** | 3.5 | 0.177942 | 4.94481 | 0.72870:0.27130 |
| **Omitting 66-88** | 3.6 | 0.163209 | 5.01186 | 0.71076:0.28924 |
| **Omitting 271-276** | 4.6 | 0.171485 | 5.06270 | 0.74165:0.25835 |
| **Omitting 66-88 + 271-276** | 4.6 | 0.154807 | 4.95933 | 0.72677:0.27323 |

| Mix IF+OF (xylose) | $\gamma$ | Weighted RMSE to target | Applied work (kJ/mol) | Conformational population (IF:OF) |
|---|---|---|---|---|
| **Original** | 4.2 | 0.174327 | 4.96328 | 0.61804:0.38196 |
| **Omitting 66-88** | 4.4 | 0.158985 | 5.06920 | 0.60683:0.39317 |
| **Omitting 271-276** | 5.5 | 0.164855 | 5.00366 | 0.43961:0.56039 |
| **Omitting 66-88 + 271-276** | 5.8 | 0.148635 | 5.05329 | 0.42037:0.57963 |

| Mix IF+OF (glucose) | $\gamma$ | Weighted RMSE to target | Applied work (kJ/mol) | Conformational population (IF:OF) |
|---|---|---|---|---|
| **Original** | 4.7 | 0.144861 | 4.98085 | 0.44715:0.55285 |
| **Omitting 66-88** | 4.7 | 0.137190 | 4.95480 | 0.44391:0.55609 |
| **Omitting 271-276** | 6.5 | 0.128285 | 4.96765 | 0.14566:0.85434 |
| **Omitting 66-88 + 271-276** | 6.8 | 0.118516 | 5.04470 | 0.14327:0.85673 |

| Mix IF+OF (Phloretin) Representative #1 | $\gamma$ | Weighted RMSE to target | Applied work (kJ/mol) | Conformational population (IF:OF) |
|---|---|---|---|---|
| **Original** | 4.4 | 0.179977 | 5.03989 | 0.72698:0.27302 |
| **Omitting 66-88** | 4.5 | 0.167363 | 4.98114 | 0.77315:0.22685 |
| **Omitting 271-276** | 6.9 | 0.165217 | 5.04423 | 0.60736:0.39264 |
| **Omitting 66-88 + 271-276** | 7.2 | 0.150428 | 5.04090 | 0.66203:0.33797 |

| Mix IF+OF (Phloretin) Representative #2 | $\gamma$ | Weighted RMSE to target | Applied work (kJ/mol) | Conformational population (IF:OF) |
|---|---|---|---|---|
| **Original** | 4.9 | 0.181900 | 4.96968 | 0.83969:0.16031 |
| **Omitting 66-88** | 4.9 | 0.165580 | 4.99049 | 0.86608:0.13392 |
| **Omitting 271-276** | 5.7 | 0.165947 | 4.93941 | 0.83969:0.16031 |
| **Omitting 66-88 + 271-276** | 6.5 | 0.151093 | 4.97699 | 0.67538:0.32462 |

| Mix IF+OF (Phloretin) Representative #1+2 | $\gamma$ | Weighted RMSE to target | Applied work (kJ/mol) | Conformational population (IF:OF) |
|---|---|---|---|---|
| **Original** | 4.6 | 0.180121 | 4.98044 | 0.78256:0.21744 |
| **Omitting 66-88** | 4.7 | 0.165396 | 4.94515 | 0.82852:0.17148 |
| **Omitting 271-276** | 6.2 | 0.164322 | 5.04684 | 0.62753:0.37247 |
| **Omitting 66-88 + 271-276** | 6.7 | 0.149445 | 5.03621 | 0.67511:0.32489 |

| Mix IF+OF (Phloridzin) Representative #1 | $\gamma$ | Weighted RMSE to target | Applied work (kJ/mol) | Conformational population (IF:OF) |
|---|---|---|---|---|
| **Original** | 4.7 | 0.148382 | 5.04179 | 0.44563:0.55437 |
| **Omitting 66-88** | 4.7 | 0.138700 | 4.97599 | 0.46246:0.53754 |
| **Omitting 271-276** | 7.6 | 0.124186 | 4.95055 | 0.34598:0.65402 |
| **Omitting 66-88 + 271-276** | 8.5 | 0.116034 | 5.04881 | 0.37590:0.62410 |

| Mix IF+OF (Phloridzin) Representative #2 | $\gamma$ | Weighted RMSE to target | Applied work (kJ/mol) | Conformational population (IF:OF) |
|---|---|---|---|---|
| **Original** | 5.3 | 0.154079 | 4.95580 | 0.45107:0.54893 |
| **Omitting 66-88** | 5.6 | 0.142263 | 5.05621 | 0.46646:0.53354 |
| **Omitting 271-276** | 9.2 | 0.132222 | 5.03193 | 0.38013:0.61987 |
| **Omitting 66-88 + 271-276** | 9.7 | 0.116802 | 5.04263 | 0.39750:0.60250 |

| Mix IF+OF (Phloridzin) Representative #1+2 | $\gamma$ | Weighted RMSE to target | Applied work (kJ/mol) | Conformational population (IF:OF) |
|---|---|---|---|---|
| **Original** | 4.8 | 0.149978 | 4.98607 | 0.45319:0.54681 |
| **Omitting 66-88** | 4.9 | 0.139437 | 4.97173 | 0.46834:0.53166 |
| **Omitting 271-276** | 8.6 | 0.1225890 | 5.01158 | 0.36350:0.63650 |
| **Omitting 66-88 + 271-276** | 9.4 | 0.115578 | 4.99555 | 0.38712:0.61288 |

**Table S4 HDX-MS reweighting experiments of mixing 10 ensemble structures.**

| Mix 10 to fit apo | $\gamma$ | Weighted RMSE to target | Applied work (kJ/mol) | Conformational population (IF:OF) |
|---|---|---|---|---|
| **Original** | 3.8 | 0.167557 | 5.07162 | 0.58981:0.41019 |
| **Omitting 66-88** | 4.0 | 0.156539 | 4.97036 | 0.62006:0.37994 |
| **Omitting 271-276** | 5.1 | 0.152565 | 4.95515 | 0.44674:0.55326 |
| **Omitting 66-88 + 271-276** | 5.6 | 0.140929 | 5.00836 | 0.49493:0.50307 |

| Mix 10 to fit xylose | $\gamma$ | Weighted RMSE to target | Applied work (kJ/mol) | Conformational population (IF:OF) |
|---|---|---|---|---|
| **Original** | 3.7 | 0.163952 | 4.92331 | 0.49399:0.50601 |
| **Omitting 66-88** | 4.0 | 0.153689 | 4.98495 | 0.51704:0.48296 |
| **Omitting 271-276** | 5.0 | 0.150334 | 5.06030 | 0.29957:0.70043 |
| **Omitting 66-88 + 271-276** | 5.4 | 0.141076 | 5.02943 | 0.34622:0.65378 |

| Mix 10 to fit glucose | $\gamma$ | Weighted RMSE to target | Applied work (kJ/mol) | Conformational population (IF:OF) |
|---|---|---|---|---|
| **Original** | 4.2 | 0.140658 | 5.07313 | 0.32059:0.67941 |
| **Omitting 66-88** | 4.3 | 0.136612 | 5.02650 | 0.32732:0.67268 |
| **Omitting 271-276** | 5.4 | 0.121775 | 4.96685 | 0.11498:0,88502 |
| **Omitting 66-88 + 271-276** | 5.8 | 0.118668 | 5.01107 | 0.12321:0.87679 |

| Mix 10 to fit phloretin | $\gamma$ | Weighted RMSE to target | Applied work (kJ/mol) | Conformational population (IF:OF) |
|---|---|---|---|---|
| **Original** | 3.6 | 0.179290 | 4.95745 | 0.73512:0.26488 |
| **Omitting 66-88** | 3.8 | 0.164210 | 5.01612 | 0.76382:0.23618 |
| **Omitting 271-276** | 5.2 | 0.164627 | 5.01602 | 0.65402:0.34598 |
| **Omitting 66-88 + 271-276** | 5.6 | 0.148582 | 5.02416 | 0.69496:0.34504 |

| Mix 10 to fit phloridzin | $\gamma$ | Weighted RMSE to target | Applied work (kJ/mol) | Conformational population (IF:OF) |
|---|---|---|---|---|
| **Original** | 4.5 | 0.129704 | 4.96615 | 0.68902:0.31098 |
| **Omitting 66-88** | 4.7 | 0.117545 | 5.04124 | 0.71142:0.28858 |
| **Omitting 271-276** | 7.5 | 0.110742 | 4.96434 | 0.56190:0.43810 |
| **Omitting 66-88 + 271-276** | 8.5 | 0.098408 | 5.03106 | 0.60442:0.39558 |

**Table S5 RMSE value of peptides after reweighting.**

| | Peptides with RMSE over 0.3 | Common peptides with RMSE over 0.3 | Unique peptides with RMSE over 0.3 | Union peptides with RMSE over 0.3 |
|---|---|---|---|---|
| **Mix IF+OF (apo)** | 10-15 (0.35606); 49-56 (0.32942); 271-276(0.45235); 305-315(0.32282); 435-445(0.33012); 436-444 (0.37610) | 271-276 | 10-15; 49-56; 305-315; 435-445; 436-444 | 10-15; 43-48; 49-56; 227-240; 241-248; 271-276; 298-304; 300-305; 305-315; 393-409; 395-409; 435-445; 436-444 |
| **Mix IF+OF (xylose)** | 10-15(0.32147; 49-56(0.36862); 241-248(0.33938); 271-276(0.43909); 300-305(0.30507); | | 10-15; 49-56; 241-248; 300-305 | |
| **Mix IF+OF (glucose)** | 49-56(0.33397); 271-276(0.53095); 298-304(0.32483); 300-305(0.35711); | | 49-56; 298-304; 300-305 | |
| **Mix IF+OF (phloretin)** | 10-15(0.39923); 43-48(0.34699); 241-248(0.39807); 271-276(0.56135); 393-409(0.33052); 395-409(0.30392) | | 10-15; 43-48; 241-248; 393-409;395-409 | |
| **Mix IF+OF (phloridzin)** | 43-48 (0.33259); 227-240(0.32147); 241-248(0.33703); 271-276(0.62760); | | 43-48; 227-240; 241-248 | |

**Table S6 HDX-MS reweighting experiments of mixing 14 ensemble structures.**

| Mix 14 to fit apo | $\gamma$ | Weighted RMSE to target | Applied work (kJ/mol) | Conformational population (IF:OF) |
|---|---|---|---|---|
| **Original** | 3.7 | 0.167289 | 5.04337 | 0.61204:0.38796 |
| **Omitting 66-88** | 3.9 | 0.154267 | 5.06030 | 0.65093:0.34907 |
| **Omitting 271-276** | 5.0 | 0.151724 | 5.02817 | 0.45409:0.54591 |
| **Omitting 66-88 + 271-276** | 5.3 | 0.138593 | 5.00156 | 0.51161:0.48839 |

| Mix 14 to fit xylose | $\gamma$ | Weighted RMSE to target | Applied work (kJ/mol) | Conformational population (IF:OF) |
|---|---|---|---|---|
| **Original** | 3.6 | 0.163218 | 4.98732 | 0.50212:0.49788 |
| **Omitting 66-88** | 3.8 | 0.152143 | 4.93235 | 0.53420:0.46580 |
| **Omitting 271-276** | 4.9 | 0.149907 | 5.05656 | 0.29938:0.70062 |
| **Omitting 66-88 + 271-276** | 5.2 | 0.139319 | 4.99465 | 0.37072:0.62928 |

| Mix 14 to fit glucose | $\gamma$ | Weighted RMSE to target | Applied work (kJ/mol) | Conformational population (IF:OF) |
|---|---|---|---|---|
| **Original** | 3.9 | 0.139824 | 4.95374 | 0.34900:0.65100 |
| **Omitting 66-88** | 4.0 | 0.135521 | 4.97450 | 0.36545:0.63455 |
| **Omitting 271-276** | 5.6 | 0.121471 | 5.02286 | 0.12684:0.87316 |
| **Omitting 66-88 + 271-276** | 5.9 | 0.117855 | 5.01597 | 0.15196:0.84804 |

| Mix 14 to fit phloretin | $\gamma$ | Weighted RMSE to target | Applied work (kJ/mol) | Conformational population (IF:OF) |
|---|---|---|---|---|
| **Original** | 3.7 | 0.179584 | 5.07340 | 0.75874:0.24126 |
| **Omitting 66-88** | 3.8 | 0.164504 | 4.94549 | 0.78855:0.21145 |
| **Omitting 271-276** | 5.2 | 0.165276 | 4.99458 | 0.65387:0.34613 |
| **Omitting 66-88 + 271-276** | 5.6 | 0.149153 | 4.95957 | 0.69457:0.30543 |

| Mix 14 to fit phloridzin | gamma | Weighted RMSE to target | Applied work (kJ/mol) | Conformational population (IF:OF) |
|---|---|---|---|---|
| **Original** | 4.6 | 0.129659 | 5.01028 | 0.72563:0.27437 |
| **Omitting 66-88** | 4.8 | 0.116872 | 5.05214 | 0.75229:0.24771 |
| **Omitting 271-276** | 7.5 | 0.110742 | 5.00505 | 0.56482:0.43518 |
| **Omitting 66-88 +271-276** | 8.4 | 0.098166 | 4.98115 | 0.60631:0.39369 |

**Table S7 HDX-MS data comparison between previously published and newly generated data for XylE WT**. MaxD experiments for newly generated data were performed in parallel with the new HDX labelling, but not for previously published data.

| start | end | Uptake (5 min) Previous data | Uptake (5 min) New data | Uptake (30 min) Previous data | Uptake (30 min) New data | MaxD For previous data | MaxD For new data |
|---|---|---|---|---|---|---|---|
| 10 | 15 | 0.988023 | 1.341604 | 1.648423 | 2.178138 | 3.381621 | 3.138548 |
| 16 | 23 | 0.036203 | 0.02243 | -0.04094 | 0.050032 | 4.731473 | 4.337314 |
| 29 | 36 | 0.360692 | 1.983312 | 0.712998 | 2.599542 | 5.022441 | 4.302648 |
| 29 | 38 | 0.47115 | 2.554559 | 1.029097 | 3.625274 | 6.525283 | 5.759576 |
| 31 | 38 | 0.42444 | 1.677353 | 1.06695 | 2.609273 | 4.518544 | 4.027515 |
| 41 | 48 | 0.904589 | 1.864013 | 1.302296 | 2.474225 | 4.039788 | 3.752867 |
| 43 | 48 | 0.82694 | 1.597494 | 1.21237 | 1.940894 | 2.835552 | 2.664079 |
| 49 | 56 | 1.026593 | 2.66125 | 1.03743 | 3.597548 | 4.591334 | 4.115131 |
| 68 | 88 | 3.383246 | 3.761676 | 4.186496 | 4.558251 | 9.930536 | 10.58284 |
| 110 | 122 | 3.891694 | 5.321523 | 5.397606 | 6.291501 | 6.913373 | 6.430672 |
| 110 | 124 | 4.028711 | 5.820736 | 5.891508 | 7.021107 | 8.349839 | 7.944533 |
| 122 | 129 | 0.161974 | 0.501083 | 0.294865 | 0.817584 | 3.950091 | 3.385067 |
| 143 | 150 | 0.506214 | 0.731701 | 1.091809 | 1.460049 | 3.949765 | 3.422853 |
| 146 | 150 | 0.447891 | 0.611255 | 0.805693 | 0.963076 | 1.868529 | 1.573661 |
| 164 | 169 | 0.576433 | 0.863486 | 1.101133 | 1.394248 | 3.424384 | 2.873047 |
| 169 | 173 | 0.052192 | 0.048571 | 0.095737 | 0.092673 | 2.900675 | 2.574969 |
| 217 | 226 | 2.413823 | 2.737934 | 2.843405 | 3.250789 | 5.307931 | 4.237368 |
| 227 | 240 | 6.632008 | 6.57535 | 6.600157 | 6.749862 | 8.180742 | 6.843217 |
| 241 | 248 | 3.545824 | 3.652138 | 3.82323 | 3.822171 | 4.69903 | 4.176144 |
| 298 | 304 | 0.051406 | 0.649032 | 0.06824 | 1.495712 | 3.287716 | 2.846126 |
| 344 | 348 | 0.015812 | 0.067183 | 0.018998 | 0.109736 | 2.837236 | 2.651964 |
| 395 | 409 | 2.281486 | 2.072208 | 3.36335 | 4.056172 | 9.301031 | 7.905648 |

**Table S8 Modification of residue protonation states and Histidine sidechain flips using MolProbity[4]**

| | | | | | |
|---|---|---|---|---|---|
| **Protonation states** | **E206 = neutral** | | | | |
| **Sidechain flips** | HSE158 | HSE258 | HSD262 | HSD438 | HSE440 |

**Table S9 Energy minimisation and equilibration protocol for XylE protein structure.**

| | Length(ns) | Timestep(ps) | Ensemble | Restraint location | Restraint value (kJ $mol^{-1}$ $nm^{-2}$) | Lipid restraint location | Lipid restraint value (kJ $mol^{-1}$ $nm^{-2}$) | Thermostat/barostat |
|---|---|---|---|---|---|---|---|---|
| min | 10k steps | N/A | N/A | None | N/A | N/A | N/A | N/A |
| Eq1a | 10 | 0.001 | NVT | All protein atoms | 4000 | Phosphorus atom positional restraint in the z-direction; dihedral restraints for palmitoyl chirality | Positional:1000; Dihedral:1000 | V-rescale thermostat |
| Eq1b | 10 | 0.002 | NPT | All protein atoms | 4000 | Phosphorus atom positional restraint in the z-direction; dihedral restraints for palmitoyl chirality | Positional:40; Dihedral:100 | V-rescale thermostat; Berendsen barostat |
| Eq2 | 5 | 0.002 | NPT | All protein atoms | 4000 | N/A | N/A | V-rescale thermostat; Berendsen barostat |
| Eq3 | 5 | 0.002 | NPT | Protein backbone | 1000 | N/A | N/A | V-rescale thermostat; Berendsen barostat |
| Eq4 | 5 | 0.002 | NPT | Protein backbone | 200 | N/A | N/A | V-rescale thermostat; Berendsen barostat |
| Eq5 | 5 | 0.002 | NPT | Protein CA | 20 | N/A | N/A | V-rescale thermostat; Berendsen barostat |
| Eq6 | 60 | 0.002 | NPT | None | N/A | N/A | N/A | V-rescale thermostat; Parrinello-Rahman barostat |

**Table S10 Energy minimisation and equilibration for XylE-ligand bound structure**. Solute_all represents all protein and ligand atoms. Solute_bb is for all protein backbone atoms, Solute_sc is all protein sidechain atoms and ligand atoms, and Solute_ca is all protein alpha carbon atoms. Lipid_headgroup is the POPE phosphorus atom and lipid_dihedral represents the C1-C3-C2-O21 and C28-C29-C210-C211 dihedrals in POPE.

| | **Length(ns)** | **Timestep(ps)** | **Ensemble** | **Restraint location & value ((kJ mol$^{-1}$ nm$^{-2}$))** | **Thermostat/barostat** |
|---|---|---|---|---|---|
| **min** | 10k steps | N/A | N/A | None | N/A |
| **Eq1a** | 10 | 0.001 | NVT | Solute_all = 4000; Lipid_headgroup=1000; Lipid_dihedral = 1000 | V-rescale thermostat |
| **Eq1b** | 10 | 0.002 | NPT | Solute_all = 4000; Lipid_headgroup=40; Lipid_dihedral = 100 | V-rescale thermostat; Berendsen barostat |
| **Eq2** | 5 | 0.002 | NPT | Solute_all = 4000 | V-rescale thermostat; Berendsen barostat |
| **Eq3** | 5 | 0.002 | NPT | Solute_bb = 1000; solute_sc = 0 | V-rescale thermostat; Berendsen barostat |
| **Eq4** | 5 | 0.002 | NPT | Solute_bb = 200; solute_sc = 0 | V-rescale thermostat; Berendsen barostat |
| **Eq5** | 5 | 0.002 | NPT | Solute_ca=20 | V-rescale thermostat; Berendsen barostat |
| **Eq6** | 60 | 0.002 | NPT | None | V-rescale thermostat; Parrinello-Rahman barostat |

**Table S11 Representative XylE-inhibitor bound structures selection.**

| | | Phloretin | | Phloridzin | |
|---|---|---|---|---|---|
| | | OF | IF | OF | IF |
| SD_RMSD (nm) | Representative structure #1 | 0.04845 | 0.04166 | 0.04451 | 0.03791 |
| | Representative structure #2 | 0.05323 | 0.08742 | 0.04739 | 0.06654 |
| Number of visited starting structures | Representative structure #1 | 4/4 | 4/4 | 4/4 | 4/4 |
| | Representative structure #2 | 4/4 | 4/4 | 4/4 | 4/4 |
| Population of total frames | Representative structure #1 | 3501/24012 = 14.58% | 3811/24012 = 15.87% | 5743/24012 = 23.92% | 5955/24012 =24.80% |
| | Representative structure #2 | 4601/24012 = 19.16% | 2999/24012 = 12.49% | 4021/24012 =16.75% | 4140/24012 = 17.24% |
| The population of frames inside the cluster shares < 1.5 Å RMSD with the average structure | Representative structure #1 | 3247/3501 = 92.74% | 3771/3811 = 98.95% | 5688/5743 = 99.04% | 5903/5955 = 99.13% |
| | Representative structure #2 | 4295/4601 = 93.35% | 2778/2999 = 92.63% | 3825/4021 = 95.12% | 4091/4140 = 98.82% |
